# *In silico* identification of a molecular circadian system with novel features in the crustacean model organism *Parhyale hawaiensis*

**DOI:** 10.1101/705038

**Authors:** Benjamin James Hunt, Eamonn Mallon, Ezio Rosato

**Author notes:** Correspondence: Dr. Ezio Rosato.

## Abstract

The amphipod *Parhyale hawaiensis* is a model organism of growing importance in the fields of evolutionary development and regeneration. A small, hardy marine crustacean that breeds year-round with a short generation time, it has simple lab culture requirements and an extensive molecular toolkit including the ability to generate targeted genetic mutant lines. Here we identify canonical core and regulatory clock genes using genomic and transcriptomic resources as a first step in establishing this species as a model in the field of chronobiology. The molecular clock of *P. hawaiensis* lacks orthologs of the canonical circadian genes *cryptochrome 1* and *timeless*, in common with the mammalian system but in contrast to many arthropods including *Drosophila melanogaster*. Furthermore the predicted CLOCK peptide is atypical and CRY2 shows an extended 5’ region of unknown function. These results appear to be shared by two other amphipod species.

## Introduction

The subphylum Crustacea has a rich history of chronobiological research. One of the earliest recorded examples of a persistent circadian phenomenon in animals was the colour changes seen in the prawn *Hippolyte varians* (Gamble and Keeble, 1900), while investigations on the circadian and circatidal rhythms of the fiddler crab *Uca pugnax* helped lay the basis for modern chronobiology (Palmer, 1991), with published works on the persistence of chromatophore and locomotor activity rhythms in constant conditions; temperature compensation; persistance of rhythms in isolated tissue; and the phase-shifting effect of light pulses, this latter observation coming a decade before the phase response curve (PRC) was incorporated into circadian concepts (De Coursey, 1960). A variety of endogenously controlled rhythmic phenomena have been documented in wide range of crustaceans, not only circadian rhythms but also tidal, lunar and semi-lunar (Strauss and Dircksen, 2010), under the control of *zeitgebers* such as light, hydrostatic pressure, immersion, wave motion and salinity. Where the clade has historically proved lacking, however, is in providing an organism with a molecular toolkit comparable to that of the fruit fly *Drosophila melanogaster*, in which transgenics, reporter genes and binary expression systems have been used to characterise, locate and dissect the endogenous biological clock at a neurological level (Helfrich-Förster, 2005).

At the centre of the *Drosophila* clock are the bHLH-PAS proteins CLOCK (Allada *et al.*, 1998; Gekakis *et al.*, 1998, CLK) and CYCLE (Rutila *et al.*, 1998, CYC), which form a heterodimer that drives transcription of clock-controlled genes including *period* (Konopka and Benzer, 1971, per) and *timeless* (Sehgal *et al.*, 1994, tim), whose protein products dimerise and interact with CLK:CYC to inhibit its transcription activity and thus their own expression. TIM is degraded by the light-activated CRYPTOCHROME (Emery *et al.*, 1998, CRY), exposing PER to degradation in turn and ending CLK:CYC repression. The transcription-translation feedback loop enacted by these core clock genes is modified to achieve a circadian periodicity of 24 hours through the contributions of numerous regulatory genes such as *doubletime, vrille, shaggy* and *clockwork orange*. In mammals the clock is similarly well understood thanks to work on *Mus musculus* (Reppert and Weaver, 2001), showing that the molecular system is remarkably conserved but not identical. In mammals the CLK:BMAL1 heterodimer (where BMAL1 is an ortholog of CYC) is inhibited by multiple PER and CRY proteins, the latter notably different to the *Drosophila* CRY in lacking photosensitivity. A number of species (Zhu *et al.*, 2005) have been found to have both a *Drosophila*-like CRY (often termed CRY1) and a mammalian-like CRY (CRY2).

In crustaceans, the first core clock gene to be identified was *clock* in the prawn *Macrobrachium rosenbergii* (Yang *et al.*, 2006). The falling cost of next generation sequencing has subsequently brought a wave of such data in crustacean species including the water flea *Daphnia pulex* (Tilden *et al.*, 2011), the copepods *Calanus finmarchicus* (Christie *et al.*, 2013) and *Tigriopus californicus* (Nesbit and Christie, 2014), the lobster *Nephrops norvegicus* (Sbragaglia *et al.*, 2015), the amphipod *Talitrus saltator* (O’Grady *et al.*, 2016) and the Antarctic krill *Euphausia superba* (Hunt *et al.*, 2017), in some cases including identification of rhythmic transcription. The clock of *E. superba* has been further characterised through analysis of the interactions of its protein components (Biscontin *et al.*, 2017), as has that of the isopod *Eurydice pulchra* (Zhang *et al.*, 2013), while tissue-specific identification of clock components has been performed in the American lobster *Homarus americanus* (Christie *et al.*, 2018a,b).

Nevertheless much work remains to be done in determining how these genes generate rhythms, and in what tissues and cells they are active and when, and with which genes they interact. More fine-grained investigation into the crustacean clock requires a model organism of similar power to *D. melanogaster*, and the establishment of such a model also promises to significantly advance our understanding of the animal clock through comparative analysis.

The amphipod *Parhyale hawaiensis* was first isolated for lab culture in 1997 from the filtration system of the John G. Shedd Aquarium, “preselected [for] minimal care”, in the words of the collectors (Browne *et al.*, 2005). A circumtropical marine detritivore found in shallow water mangrove zones, it is a robust, fecund organism that is cheap to maintain and breeds year-round with a quick generation time, making it well-suited to lab culture. Thanks to a reproductive system that permits the collection and manipulation of embryos from a single cell stage this species has become established as a model organism of some significance in the field of development. It has further proven itself to be amenable to molecular analysis and manipulation, including *in situ* hybridisation, RNAi-mediated knockdown, transposon and integrase-based genetic transformation and CRISPR/Cas9 mutagenesis, and recently the 3.6 Gb genome was published (Kao *et al.*, 2016).

Here we describe the creation of a head transcriptome to identify core and regulatory clock genes and use the genome to gain independent verification of our findings, which include the apparent loss of a number of major clock genes and the identification of a peptide feature that may be unique to amphipods.

## Materials and Methods

### Animal husbandry

Animals were maintained in polypropylene containers with lids 3/4 on, with a layer of crushed coral approximately 1 cm deep and filled with 1 L artificial seawater prepared using 30g Tetra Marine Sea Salt per litre of distilled tap water to give a specific gravity of 1.022. Cultures were maintained on a 12:12 light:dark cycle with lights on at 9am, illumination coming from fluorescent room lighting at 300 lux, at 20°C. Animals were fed twice weekly, once with shredded carrot and once with a pinch of standard fish flakes.

### Head transcriptome

At eight time-points across 24 hours beginning at 9am (lights on, *zeitgeber* time (ZT) 0) and every three hours thereafter, a total of 20 adult males were collected per time-point and snap-frozen on a metal plate atop dry ice, whereupon the heads were dissected and stored at −80°C until use. Total RNA was extracted from five heads per time-point using a micropestle to homogenise the tissue in TRIzol reagent (Invitrogen), cleaned using the PureLink Micro Kit (ThermoFisher Scientific) with on-column DNase treatment to remove genomic DNA contamination and eluted with RNase free water. RNA extractions were quality assessed using Nanodrop and Agilent 2100 Bioanalyzer. Samples were required to meet a minimum quality of > 50 ng μl^−1^ concentration; 260/280 ratio 1.8 – 2.2; 260/230 ratio 2.0 - 2.4 for sequencing use, and those that failed were discarded and the extraction process repeated on another five heads from storage.

One sample per time-point was delivered to Glasgow Polyomics (University of Glasgow) and subject to polyA mRNA enrichment with the TruSeq Stranded mRNA Library Prep Kit (Illumina). The resultant libraries were sequenced using the Illumina NextSeq 500 platform to generate 75 bp paired-end strand-specific reads, which were subject to gentle quality control comprising low quality read removal < Q10 and adapter trimming by Glasgow Polyomics. After download of this dataset FastQC 0.11.2 (Andrews, 2010) was used to assess read quality and Cutadapt 1.9.1 (Martin, 2011) used to remove remaining traces of the TruSeq Index and Universal adapters using the sequence “GATCGGAAGAGC” common to both adapter types as criteria. Reads were then filtered using Trimmomatic 0.32 (Bolger *et al.*, 2014) to remove reads with an average quality of less than 20 and/or shorter than 72 bases.

A multi-k-mer, multi-assembler approach was adopted for assembly. Trans-ABySS 1.5.5 (Robertson *et al.*, 2010), Bridger 2014-12-01 (Chang *et al.*, 2015) and Trinity r20140717 (Haas *et al.*, 2013) were used to generate individual assemblies using k-mers ranging from 19 to 71 with a minimum contig length of 200 bp. These were subsequently merged and duplicates and containments were removed at 99% identity using CD-HIT-EST 4.6.6 (Li and Godzik, 2006). TransRate (Smith-Unna *et al.*, 2016) was used to identify well-assembled contigs based on read-mapping evidence. TransDecoder 3.0.0 (Haas *et al.*, 2013) was used to identify predicted peptides from the well-assembled contigs (single_best_orf, -m 90), and CD-HIT used at 90% peptide identity to produce a final list of well-assembled, non-redundant coding contigs that were extracted from the original merged transcriptome assembly and used for all downstream purposes. Transcriptome completeness was assessed using BUSCO (Waterhouse *et al.*, 2018) against the arthropoda lineage dataset.

### Identification of clock genes

The head transcriptome was searched for core and regulatory circadian genes using the tblastn function of BLAST+ with an E-value cutoff of 1E^−5^ (Table 1). The protein sequences inferred from the genome (Kao *et al.*, 2016, see https://doi.org/10.6084/m9.figshare.3498104.v4) were searched in the same manner using blastp. Genes that returned little or no evidence through this approach were searched for in the genome using tblastn and, where evidence was subsequently obtained, the scaffold and coordinates of the alignments were noted. The head transcriptome reads were aligned to the genome using HISAT2 2.1.0 (Kim *et al.*, 2015) and visualised with IGV 2.3.82 (Robinson *et al.*, 2011), and Stringtie 1.3.3b (Pertea *et al.*, 2015) was used to assemble transcripts of interest from these read alignments. Gene sequences identified through BLAST+ searches were extended using these alignments and transcripts where appropriate. Sequence data from two other amphipods, *Hyalella azteca* (GCA_000764305.2) and *Talitrus saltator* (GDUJ00000000) was further used to extend partial transcripts and lend support to atypical results. RNA-seq reads for *H. azteca* (PRJNA312414) were downloaded, aligned and assembled as above. *P. hawaiensis* sequences obtained from from BLAST+ searches and contigs identified as circadian genes in *T. saltator* (O’Grady *et al.*, 2016) were used to search the genomes/transcriptomes of all three species, with results expanded using read mapping evidence and then these extended sequences used to search all resources again until no further extension was possible. Sequences returned by these processes were translated to peptides and subject to validation using the NCBI BLAST web interface (blastp, Database ‘nr’, Entrez Query ‘all[filter] NOT predicted[title] NOT hypothetical[title] NOT putative[title]’) to ascertain their closest match in other species.

**Table 1:** *P. hawaiensis* clock gene candidates identified from transcriptome and genome mining. nt = nucleotides. aa = amino acids. CK = Casein kinase. PDH = Pigment dispersing hormone. PDP = Par domain protein. All searches were conducted using complete query sequences.

| Protein | Query | Species | *P. hawaiiensis* contig | E-value | Length (nt) | Length (aa) |
| --- | --- | --- | --- | --- | --- | --- |
| BMAL1 | ANW48377 | *E. superba* | phaw_30_tra_m.024548 | 2.00E-156 | - | 617 |
| CK II $\alpha$ | AAN11415 | *D. melanogaster* | PH.k29.comp2312_seq0 | 0 | 4,885 | 352 |
| CK II $\beta$ | AAF48093 | *D. melanogaster* | PH.k21.comp3428_seq3 | 5.00E-110 | 2,357 | 222 |
| CIRCADIAN TRIP | AAF52092 | *D. melanogaster* | PH.k31.comp1615_seq0 | 2.00E-111 | 12,125 | 2,996 |
| CLOCKWORK ORANGE | AAF54527 | *D. melanogaster* | PH.k21.comp80764_seq0 | 3.00E-26 | 642 | 212 |
| CRYPTOCHROME 2 | AFV96168 | *T. saltator* | PH.k21.R17911710 | 0 | 3,301 | 931 |
| DOUBLETIME | AAF57110 | *D. melanogaster* | PH.k27.comp42178_seq0 | 0 | 1,599 | 418 |
| JETLAG | AAF52178 | *D. melanogaster* | - | - | - | - |
| MICROTUBULE STAR (PP2A) | AAF52567 | *D. melanogaster* | PH.k21.comp640_seq0 | 0 | 3,538 | 309 |
| NEJIRE | AAF46516 | *D. melanogaster* | PH.k51.J3947083 | 0 | 5,384 | 1,794 |
| NEMO | AAF50497 | *D. melanogaster* | PH.c118708_g1_i2 | 0 | 1,416 | 430 |
| PDP 1 $\epsilon$ | AAF04509 | *D. melanogaster* | phaw_30_tra_m.019467 | 2.00E-35 | - | 591 |
| PERIOD | ALC74274 | *N. norvegicus* | PH.k25.comp125155_seq0 | 4.00E-07 | 280 | 91 |
| PDH | GAJQ01005023 | *H. azteca* | PH.k41.S6600835 | 2.00E-24 | 795 | 103 |
| PDH RECEPTOR | AAF45788 | *D. melanogaster* | PH.k21.comp140546_seq0 | 5.00E-29 | 597 | 191 |
| PROTEIN PHOSPHATASE 1 | CAA39820 | *D. melanogaster* | PH.k27.comp8511_seq1 | 0 | 2,963 | 328 |
| SHAGGY | AAN09084 | *D. melanogaster* | PH.k29.comp7576_seq1 | 0 | 5,244 | 457 |
| SUPERNUMERARY LIMBS | AAF55853 | *D. melanogaster* | PH.k21.comp23826_seq4 | 0 | 4,640 | 642 |
| TIMELESS | ANW48379 | *E. superba* | phaw_30_tra_m.019341 | 1.00E-10 | - | 1,674 |
| TWINS (PP2A) | AAF54498 | *D. melanogaster* | PH.k21.R18132040 | 0 | 2,887 | 444 |
| VRILLE | AAF52237 | *D. melanogaster* | PH.k21.comp5330_seq6 | 3.00E-42 | 3,178 | 480 |
| WIDERBORST (PP2A) | AAF56720 | *D. melanogaster* | PH.k29.comp1930_seq3 | 0 | 9,834 | 458 |

Peptide domain analysis was conducted using SMART (Letunic *et al.*, 2015). Peptide sequences were visualised using DOG 2.0 (Ren, et al, 2009). Phylogenetic trees were created using MEGA 7.0.26 (Kumar *et al.*, 2016); peptide sequences (Table S1) were aligned using MUSCLE at default settings, and neighbour-joining trees generated using 1,000 bootstrap replications and complete deletion.

## Results

### Transcriptome assembly and mining

After quality control the RNA-seq library for each head timepoint sample was approximately 22 million read pairs, with a total of 178,028,148 used in a final assembly of 86,686 contigs with an N50 of 2,797 and contig length ranging from 267 – 35,494. BUSCO analysis indicated 91.1% complete BUSCOs (81.1% single copy, 10% duplicated), 5.1% present but fragmented and 3.8% missing from a dataset of 1,066 groups searched. This Transcriptome Shotgun Assembly project has been deposited at GenBank under the accession GFVL00000000.

Evidence was found for the existence of the core clock genes *bmal1*, *period* and the mammalian-like *cryptochrome 2* in this assembly. Also identified were 16 regulatory clock genes (Table 1, Table 2).

**Table 2:** BLAST search results using *P. hawaiensis* clock gene candidates vs. NCBI non-redundant protein database.

| Protein | Accession | Name | Species | E-value |
| --- | --- | --- | --- | --- |
| Es-BMAL1 | AFV39705 | Bmal1a | *Pacifastacus leniusculus* | 1.00E-161 |
| Es-CASEIN KINASE II $\alpha$ | ARJ31756 | Casein kinase II subunit alpha | *Litopenaeus vannamei* | 0 |
| Es-CASEIN KINASE II $\beta$ | ANO53990 | CKII beta | *Limulus polyphemus* | 1.00E-145 |
| Es-CIRCADIAN TRIP | NP_001084531 | E3 ubiquitin-protein ligase TRIP12 | *Tribolium castaneum* | 0 |
| Es-CLOCKWORK ORANGE | KOC64192 | Hairy/enhancer-of-split related with YRPW motif protein 1 | *Habropoda laboriosa* | 3.00E-34 |
| Es-CRYPTOCHROME 2 | AFV96168 | cryptochrome 2 | *Talitrus saltator* | 0 |
| Es-DOUBLETIME | AUI80371 | Doubletime | *Euphausia superba* | 0 |
| Es-MICROTUBULE STAR (PP2A) | KDR18186 | PP2A catalytic subunit alpha | *Zootermopsis nevadensis* | 0 |
| Es-NEJIRE | CUT08824 | CREB-binding protein | *Blattella germanica* | 0 |
| Es-NEMO | KZS07628 | Mitogen-activated protein kinase | *Daphnia magna* | 0 |
| Es-PAR DOMAIN PROTEIN 1 $\epsilon$ | ABV22507 | PAR domain protein 1 | *Danaus plexippus* | 2.00E-46 |
| Es-PERIOD | ANN13870 | period | *Macrobrachium nipponense* | 4.00E-07 |
| Es-PDH | WP_082125040 | ribonuclease P protein component | *Erythrobacter luteus* | 6.8 |
| Es-PDH RECEPTOR | BAH85843 | Pigment dispersing hormone receptor | *Marsupenaeus japonicus* | 2.00E-103 |
| Es-PROTEIN PHOSPHATASE 1 | AQZ36563 | Serine/threonine protein phosphatase 1 | *Penaeus monodon* | 0 |
| Es-SHAGGY | ALK82316 | glycogen synthase kinase 3 beta | *Procambarus clarkii* | 0 |
| Es-SUPERNUMERARY LIMBS | KDR19729 | F-box/WD repeat-containing protein 1A | *Zootermopsis nevadensis* | 0 |
| Es-TIMELESS | AUI80375 | timeless 2 | *Euphausia superba* | 0 |
| Es-TWINS (PP2A) | AFK24473 | serine/threonine-protein phosphatase PP2A reg. subunit B | *Penaeus monodon* | 0 |
| Es-VRILLE | AUI80373 | Vrille | *Euphausia superba* | 1.00E-57 |
| Es-WIDERBORST (PP2A) | KYB26642 | PP2A regulatory subunit epsilon | *Tribolium castaneum* | 0 |

### BMAL1/CYCLE

*P. hawaiensis bmal1 (Ph-bmal1)*, found in complete form in the genome-inferred peptides (phaw_30_tra_m.024548) and in fragmentary form in the head transcriptome assembly, shows the typical complement of domains when translated (Figure 1A), with a bHLH domain, two PAS domains, a PAC domain immediately following the second PAS, and a BMAL1 C-terminal Region (BCTR), a transactivation domain characteristic of this gene (except in *D. melanogaster* CYCLE). The peptide returned Bmal1a in *Pacifastacus leniusculus* (accession AFV39705; 1E^−161^) from a BLAST confirmation query. *Ph-bmal1* showed very low levels of expression in the head transcriptome (Table S3).

**Figure 1:**
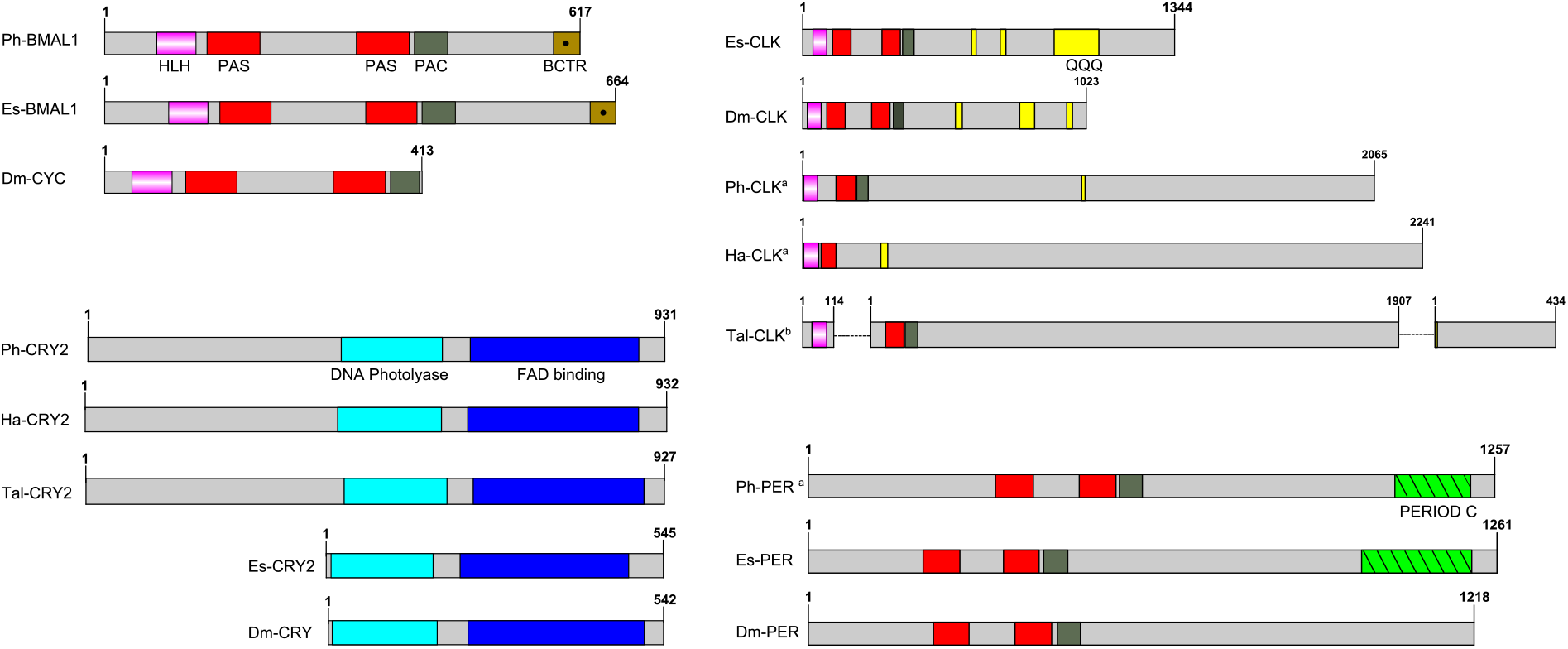
Schematics of predicted *P. hawaiensis* core clock peptides aligned with examples from other species. Ph = *Parhayle hawaiensis*. Dm = *Drosophila melanogaster*. Es = *Euphausia superba*. Tal = *Talitrus saltator*. Ha = *Hyalella azteca*. QQQ = polyglutamine domain (defined as a run of 10 or more amino acids in which 60% or more are glutamine). ^a^ - peptide is predicted on the basis of BLAST alignments and read mapping. ^b^ - peptide is a construct of multiple separate *de novo* assembled contigs. Both Tal-CRY2 and Tal-CLK have been previously reported in a shorter form (O’Grady *et al.*, 2016).

### CLOCK

No evidence was found of a *P. hawaiensis clock* gene (*Ph-clock/Ph-clk*) through BLAST mining of the head transcriptome or genome-inferred peptides. Searching the genome using *Eurydice pulchra* CLK 5 (AGV28720) identified orthologous sequences on scaffold LQNS02278184.1 (supplementary data, section A). When translated and combined, these sequences returned *clock* in the American lobster *Homarus americanus* (AWC08577; 3E^−69^), while SMART domain analysis identified bHLH, PAS and PAC domains. A small number of reads mapped to these fragments, indicating very low levels of expression in the head. Through iterative searches of the *P. hawaiensis* and *H. azteca* genomes using the CLK protein reported for the amphipod *T. saltator* (O’Grady *et al.*, 2016), and informed by read mapping evidence, this was further extended to a putative final sequence encoding a 2,065 aa peptide. A putative Ha-CLK was also identified in this manner (Figure 1B).

Searching the *T. saltator* transcriptome with Ph-CLK and Ha-CLK identified two additional contigs of interest beyond Tal-CLK (Figure 1B): GDUJ01041853 encodes a truncated peptide with a bHLH domain that returns *clock* in *Euphausia superba* (ANW48378; 6E^−24^), while GDUJ01061398 shows homology (BLAST alignment 41% identity, 1E^−40^) to the 3’ end of Ph-CLK and Ha-CLK.

### CRYPTOCHROME 2

*P. hawaiensis cryptochrome 2* (*Ph-cryptochrome 2/Ph-cry2*) was identified in complete form in the head transcriptome, returning cryptochrome 2 in *T. saltator* (AFV96168; e-value 0.0) in a BLAST confirmation query. SMART analysis of the translated 931 aa peptide identified a DNA photolyase and FAD binding domain (Figure 1C), while an extended N-terminus of 357 amino acids had no identifiable functional domain or motif. A neighbour-joining phylogenetic tree confirmed the nature of the peptide as a mammalian-like CRY2 rather than a *Drosophila*-like CRY1 (Figure 2A). This coding sequence was supported by genome search and read mapping evidence: Stringtie assembled an 11-exon transcript (supplementary data, Section A) located on scaffold LQNS02278089.1 that shows 99.5% identity to the *de novo* assembled contig, with the first exon incorporating both the N-terminus and a partial DNA photolyase domain in one contiguous sequence, indicating that the *de novo* assembled contig is unlikely to be a chimeric construct. Further to this the genome-inferred peptides, which were generated with reference to an independently assembled *de novo* transcriptome from embryonic and limb tissue (Kao *et al.*, 2016), include a contig (phaw_30_tra_m.017772) covering 189 amino acids of the N-terminus and the entire DNA photolyase domain, and another (phaw_30_tra_m.017773) covering the first 160 amino acids of the N-terminus. Of the core clock genes identified *Ph-cry2* showed the highest levels of expression (Table S3).

**Figure 2:**
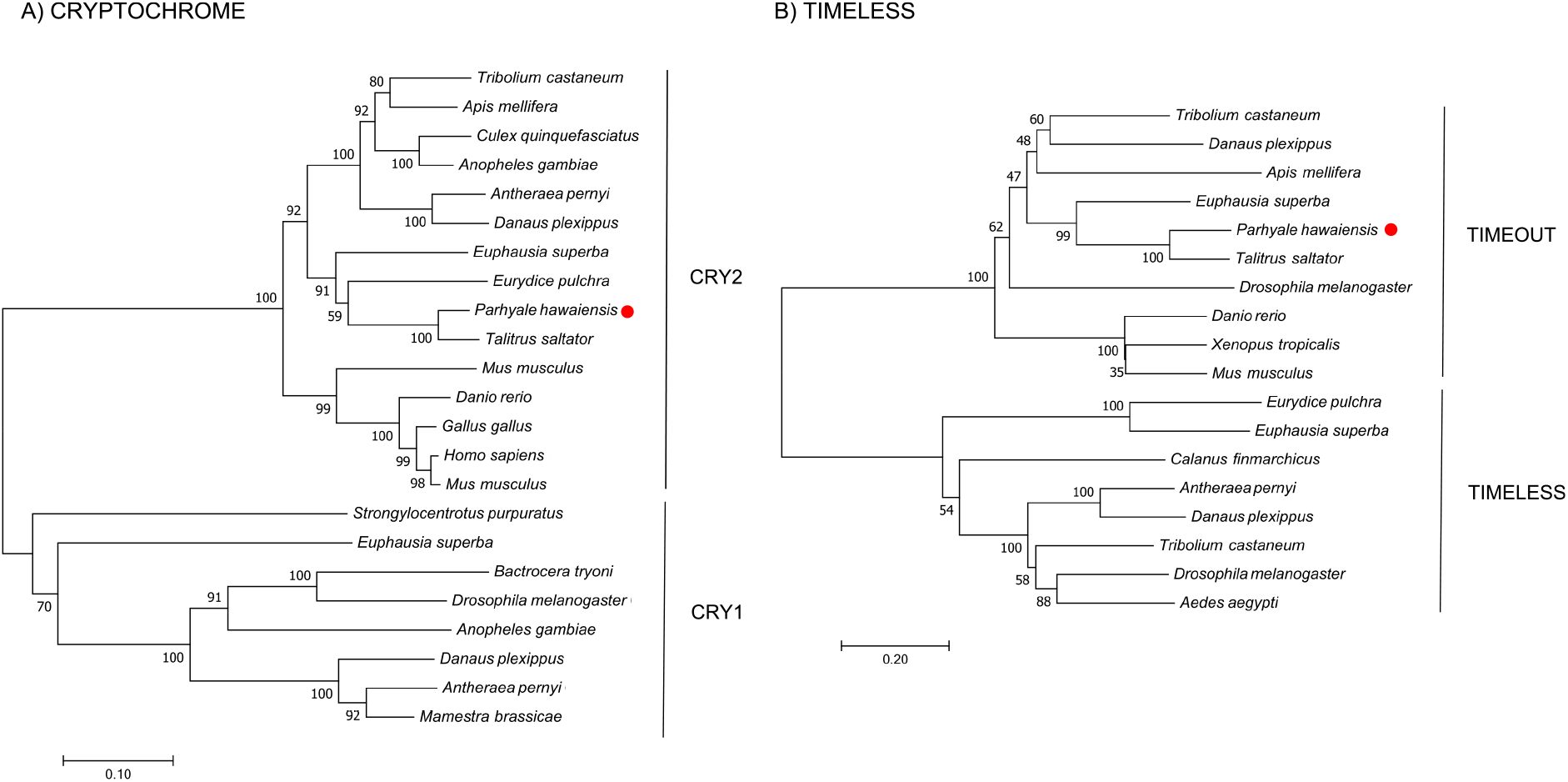
Optimal neighbour-joining trees depicting the evolutionary relationships of (A) CRYP-TOCHROME and (B) TIMELESS/TIMEOUT peptides.

No evidence was found in any *P. hawaiensis, T. saltator* or *H. azteca* resource for a *Drosophila*-like *cryptochrome*.

### PERIOD

A small 3’ fragment of *P. hawaiensis period* (*Ph-period/Ph-per*) was found in the head transcriptome, the peptide of which returned *period* in *Macrobrachium nipponense* from a BLAST confirmation query (ANN13870; e-value 4E^−07^). Searching the genome using this sequence and the predicted PER protein of *Hyalella azteca* (XP_018012603) returned multiple alignments on scaffold LQNS02276498.1 (supplementary data, Section A). A small number of head transcriptome reads mapped to 14 of these these regions. Translated and combined to form a fragmented putative peptide sequence 1,257 amino acids in length, SMART domain analysis revealed two PAS domains, a PAC domain and a PERIOD C domain (Figure 1D). This construct returned *period isoform 1* in *Homarus americanus* (AWC08578; 3E^−99^) from a BLAST confirmation query. *Ph-per* showed very low levels of expression in the head transcriptome (Table S3).

### TIMELESS

A mammalian-like TIMELESS was identified in the genome-inferred peptides resource. This is an ortholog of the *D. melanogaster timeless* paralog *timeout* (Figure 2B), a gene crucial to early development that is seemingly ubiquitous in animals (Sandrelli *et al.*, 2008) and may have only peripheral involvement with the endogenous clock. No true TIMELESS could be identified in any *P. hawaiensis*, *H. azteca* or *T. saltator* resource.

### Regulatory and clock-controlled genes

Putative orthologs were identified for the regulatory genes *casein kinase II α* and *β*, *circadian trip*, *clockwork orange*, *doubletime*, *microtubule star*, *nejire*, *nemo*, *par domain protein E*, *protein phosphatase I*, *shaggy*, *supernumerary limbs*, *twins*, *vrille* and *widerborst*, and for the clock-controlled genes *pigment dispersing hormone* (*pdh*) and *pdh receptor* (see supplementary data, Section B). Each sequence was supported through its presence in an independently assembled leg/embryo transcriptome, the genome, or both.

## Discussion

*In silico* mining of a head transcriptome resulted in the identification and characterisation of four putative core clock proteins in *Parhyale hawaiensis*; Ph-BMAL1, Ph-CLK, Ph-PER and Ph-CRY2. BUSCO analysis of this transcriptome indicated a high level of completeness and despite searching multiple additional resources including the recently published genome, we were unable to identify an ortholog of TIM or a *Drosophila*-like CRY. This suggests that *P. hawaiensis* has evolved what might be termed a mammalian-style molecular clock - a system lacking a TIM or CRY1, as originally reported in *M. musculus* (Reppert and Weaver, 2000). This system is found in vertebrates, hymenopterans and now in crustaceans. The gene reported as *T. saltator timeless* (O’Grady *et al.*, 2016) appears to be a *timeout* (Figure 2B) and no sequence encoding a CRY1 or true TIM could be found in the various transcriptomic and genomic resources available for *T. saltator* and *H. azteca*, indicating that this system may be common in amphipods. This contrasts with other crustaceans studied so far – *Daphnia pulex* possesses the full complement of core clock proteins including a CRY1 and CRY2 (Tilden *et al.*, 2011), as does *Calanus finmarchicus* (Christie *et al.*, 2013), *Trigriopus californicus* (Nesbit and Christie, 2014), *Limulus polyphemus* (Chesmore *et al.*, 2016) and *Euphausia superba* (Hunt *et al.*, 2017). *Eurydice pulchra* lacks only a CRY1 (Zhang *et al.*, 2013), as does *Nephrops norvegicus* (Sbragaglia *et al.*, 2015).

This is not the only interesting aspect of the *P. hawaiensis* circadian system. We were unable to identify a complete coding sequence for *clock*, relying on alignments and read mapping to generate an incomplete construct that is, aside from the other amphipods analysed here, notably different from the orthologs of other species. The putative amphipod CLK peptides appear to lack a second PAS domain and an extensive polyglutamine region that is characteristic of the gene (Saleem *et al.*, 2001), with only very small glutamine-rich regions identified (Figure 1B). They are also extremely long, each over 2,000 aa in partial form, far surpassing the complete *E. superba* CLK (1,344 aa) one of the longest complete CLK peptides identified to date. If the two additional *T. saltator* contigs identified in the Results are part of the full *T. saltator clock* gene this would result in a peptide at least 2,455 aa in length. In *H. azteca* and *T. saltator* there is evidence of fragmentation in this gene, with bHLH-encoding sequences present in separate scaffolds/contigs. This fragmentation was also present in the first draft of the *P. hawaiensis* genome but was resolved with the second draft, and it is likely that the fragmentation is *in silico* rather than *in vivo* for all three species. The default assumption should be that Ph-CLK forms a heterodimer with Ph-BMAL1 to drive clock gene expression as is usual in animals, but the atypical nature of the amphipod CLK peptides - in particular the apparent lack of a second PAS domain which, if accurate, may impact dimerisation - encourages speculation about alternative partners for Ph-BMAL1. In mice BMAL1 is a promiscuous binding protein (Hogenesch *et al.*, 1998) and other bHLH-PAS proteins have substituted for CLK in other species, for example methoprene-tolerent (MET) in the mosquito *Aedes aegypti* (Shin *et al.*, 2012), a putative ortholog of which can be found in fragmentary form in the head transcriptome and complete in the genome-inferred peptides (phaw_30_tra_m.006390).

In this context we consider the cryptochrome Ph-CRY2, which shows an extraordinary extension at the N-terminus, with various lines of evidence suggesting the same may be true for *H. azteca* and *T. saltator*. In *H. azteca* a predicted *cryptochrome* has been reported (accession XM_018166003), and while somewhat extended at the 5’ end it does not include an N-terminus comparable to that of *P. hawaiensis*. However a BLAST search of the genome using the Ph-CRY2 peptide returned alignments on scaffold KV721600.1 suggesting the existence of such a feature upstream of the reported sequence. Read alignment and assembly generated an 11-exon transcript (supplementary results, section A), the second exon coding for the N-terminus and partial DNA photolyase domain and terminating at the same point as the first exon of the Stringtie *Ph-cry2* transcript. When translated, this transcript encodes a peptide 932 amino acids long (Ha-CRY2) incorporating the extended N-terminus with the previously predicted *H. azteca* CRY. In the paper detailing the molecular clock of *T. saltator* (O’Grady *et al.*, 2016) two contigs were identified as representing *cry2*. An assumed complete peptide of 565 amino acids was reported, but one of the *de novo* assembled transcriptome contigs (accession GDUJ01076706.1, reported as comp100937_c0_seq1) includes an N-terminus of 362 amino acids that is not considered part of the CRY2 peptide by the authors, who verified the sequence using RACE PCR and suggest the contig may have been misassembled. However this sequence shows 55.9% identity and 71.3% similarity to the *P. hawaiensis* N-terminus, and considering the lines of evidence from *P. hawaiensis* and *H. azteca* the possibility of incomplete RACE PCR or splice variants should not be discounted. Whether this N-terminus serves a function, and what that might be, is again speculative at this point, but an unusual transcriptional heterodimer may require an unusual repressor. Perhaps then Ph-CRY2 serves in the repressive limb of the *P. hawaiensis* circadian feedback loop as does CRY2 in many other species, and it has evolved this N-terminus as a means to interact with, for example, a MET:BMAL1 heterodimer, or an atypical CLK:BMAL1.

We identified putative orthologs of all regulatory genes searched for, with the exception of *jetlag*, results that can form the basis for future work investigating if, and how, these genes curate the central feedback loops. A candidate for Ph-PDH proved difficult to identify due to its divergent sequence including at the conserved NSELINS motif. *Ph-pdh* was found only through the use of the putative *Hyalella azteca* preprohormone Hyaaz-prepro-PDH (Christie, 2014) as a query term, and while a BLAST confirmation query returned only one unrelated result (2) Ph-PDH shares 47.6% identity and 62.1% similarity with Hyaaz-prepro-PDH. A signal peptide with a predicted cleavage site between residues 36 and 37 was detected using SignalP 5.0 (Armenteros *et al.*, 2019) and through alignment with Hyaaz-prepro-PDH a 23 amino acid mature peptide was identified with 91.3% identity and 100% similarity. *T. saltator* is reported to possess two PDH peptides, the first (Tal-PDH I) unusual with regards to its long mature peptide form but showing the NSELINS motif and the second (Tal-PDH II) identified using relaxed search criteria. This latter PDH sequence and that of *H. azteca* - itself described as atypical (Christie, 2014) - were the only search terms from dozens used that returned a candidate contig. In insects the homologue of PDH, called Pigment dispersing factor (PDF) is a neuropeptide and neurohormone that is crucial for the synchronisation among the different circadian neurons (Renn *et al.*, 1999). In crustaceans PDH has two distinct types, *α*-PDH and *β*-PDH, which are found in multiple isoforms (Strauss and Dircksen, 2010), and the neuroanatomy of PDH-expressing cells has been described in numerous species, with evidence of PDH involvement in circadian rhythmicity in several of them, such as the crab *Cancer productus* in which only one of two *β*-PDH isoforms was found to be expressed in the brain and was able to substitute for *D. melanogaster* PDF in *pdf*^*01*^ flies (Beckwith *et al.*, 2011). While we have confirmation of expression in the head, the novel nature of the putative Ph-PDH requires further work to confirm its identity and to ascertain whether the sequence divergence has any impact on its functioning in the circadian pathways.

Given a lack of timepoint replicates in our head transcriptome data no firm conclusions can be drawn regarding rhythmic gene expression, though the data is suggestive of a bimodal pattern in a number of clock-related genes (Figure S1, Table S3). This transcriptomic data also suggests very low expression levels for the core clock genes in head tissue, low enough that only short fragments were assembled of *Ph-bmal1* and *Ph-period*, and nothing of *Ph-clock*. This result is consistent with the isopod *E. pulchra* in which it has been determined that Ep-PERIOD is present in only 10 cells in the brain (Zhang *et al.*, 2013).

This *in silico* analysis of these amphipod circadian systems has generated plenty of avenues for future enquiry. These sequences need to be confirmed through further sequencing, and their expression patterns more stringently characterised, both temporally and in terms of location using more extensive RNA-seq and/or RT-PCR. If it is found that Ph-CLK has been accurately described here, its ability to bind with Ph-BMAL1 and drive gene expression needs to be confirmed. The possibility of other partners for Ph-BMAL1 should also be investigated, and the functional significance of the N-terminus of Ph-CRY2 determined. Finally, further to the molecular work required it will be vital to develop behavioural and physiological assays that can be used to study the effects of clock gene manipulation. In *Drosophila melanogaster* a number of rhythmic phenomena have been identified including locomotor activity, eclosion, egg deposition, eye sensitivity, and oxygen consumption (Helfrich-Förster, 1996), while as a model organism *P. hawaiensis* has been largely exploited at a molecular level and often at early developmental stages, and behavioural observations are lacking. We have gathered encouraging preliminary locomotor activity results (Hunt, 2016), albeit compromised by issues with temperature control, hunger, age-matching and water evaporation, aeration and fouling. These results align with the observations of Poovachiranon *et al.* (1986) in indicating a nocturnal peak in activity that persists in constant darkness, with some individuals exhibiting 12 hour periodic components, a finding that may be of interest to researchers studying circa-tidal rhythms given that *P. hawaiensis* is found in inter-tidal zones. We urge interested parties to work towards optimising such behavioural assays and also to undertake the identification of other relevant characteristics that may be experimentally tractable. Only with this information in hand can one begin the task of employing *P. hawaiensis* to bring about new insights in crustacean chronobiology, a field with a rich research history that is ripe for significant advancement through the exploitation of a model with a diverse and powerful genetic toolkit.

## Acknowledgements

We are deeply grateful to Ana Patricia Ramos and Michalis Averof for supplying us with animals and husbandry advice and for permitting us to search an unpublished leg and embryo tissue assembly which proved vital in identifying the PER fragment in our own head assembly. Thanks are also due to Damian Kao, Aziz Aboobaker, Anastasios Pavlopoulos and all other authors of Kao et al., (2016) for granting us access to the then unpublished *P. hawaiensis* genome. Parts of this research first appeared in the thesis of BH (Hunt, 2016). This research used the ALICE High Performance Computing Facility at the University of Leicester.

## Author contributions

BH performed RNA extraction, *de novo* assembly, genome and transcriptome analysis, and drafted the manuscript. ER and EM conceived the study. All authors approved the final manuscript.

## Data Accessibility

The *P. hawaiensis* head transcriptome is archived at the NCBI SRA under project PRJNA399131. Newly identified/expanded putative coding and peptide sequences of core clock genes are detailed in supplementary data.

