## supplementary data for "*In silico* identification of a molecular circadian system with novel features in the crustacean model organism *Parhyale hawaiensis*"

**Supplementary methods**

The Trinity utilities ‘align_and_estimate_abundance.pl’ and ‘abundance_estimates_to_matrix.pl’ (--est_method RSEM, --aln_method bowtie2) were used to generate a TMM-normalised (Robinson and Oshlack, 2010) expression matrix by aligning the quality-trimmed reads to the head transcriptome. Values for all circadian-related contigs were extracted, along with the values for a number of putative housekeeping genes identified via BLAST search (Table S2). For each contig these were normalised to the highest value (highest = 1 after normalisation), and then the mean normalised value calculated. Finally for each timepoint the departure from the mean normalised value was calculated and plotted (Figure S1) using ggplot2 (Wickham, 2016) function in R v3.4.0 (R Core Team, 2018).

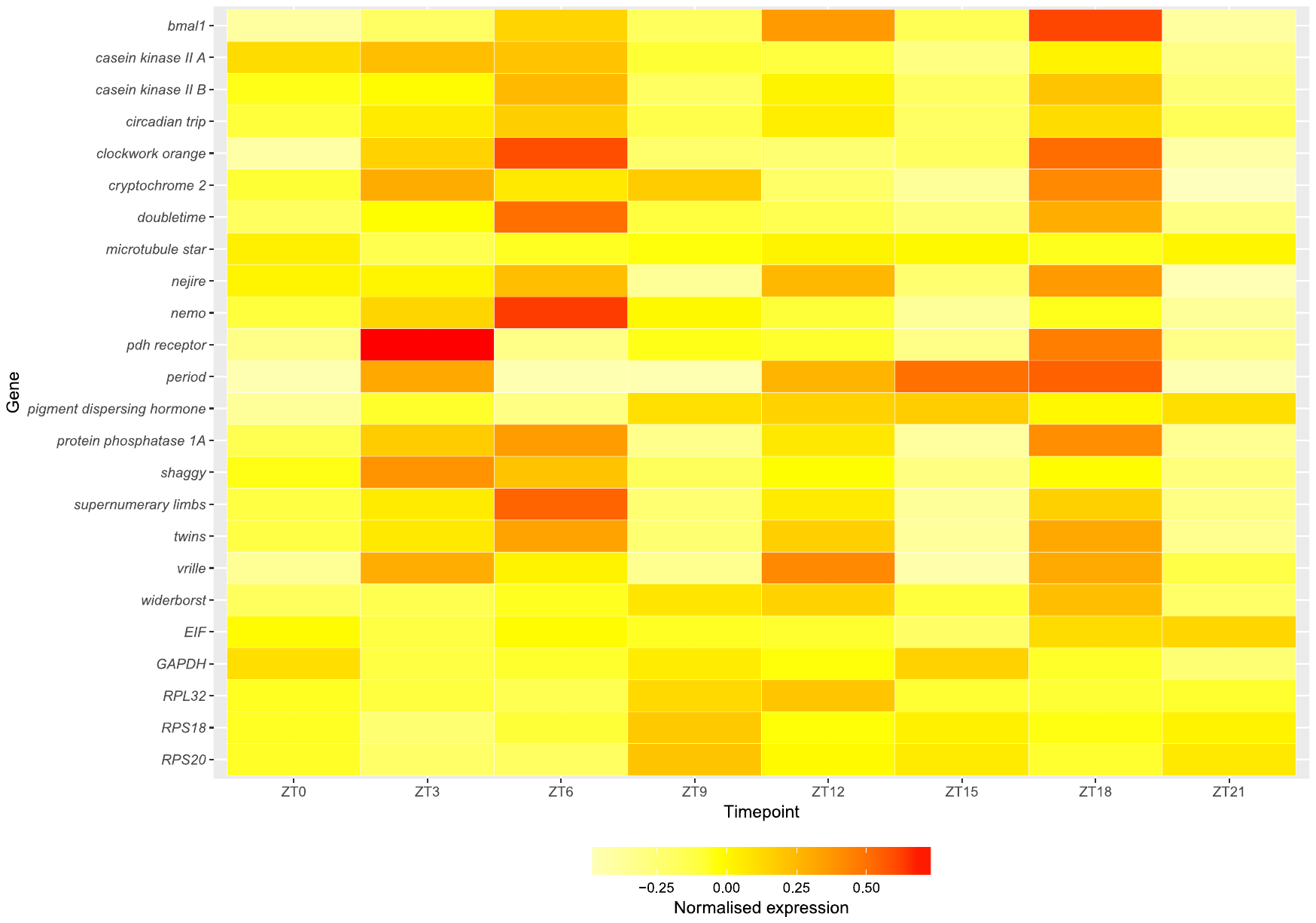

Figure S1. Expression of putative clock genes and housekeeping genes across 8 timepoints in light:dark 12:12 conditions. ZT = *zeitgeber* time, ZT0 = lights on, ZT12 = lights off.

Table S1: Accessions of peptide sequences used for generation of phylogenetic trees.

| Peptide | Accession | Species |
| --- | --- | --- |
| CRYPTOCHROME | EFA04537 | *Tribolium castaneum* |
|  | NP_001077099 | *Apis mellifera* |
|  | EDS29383 | *Culex quinquefasciatus* |
|  | ABB29887 | *Anopheles gambiae* |
|  | ABO38435 | *Antheraea pernyi* |
|  | ABA62409 | *Danaus plexippus* |
|  | CAQ86665 | *Euphausia superba* |
|  | See supplementary data | *Parhyale hawaiensis* |
|  | AFV96168 | *Talitrus saltator* |
|  | AAD39548 | *Mus musculus* |
|  | NP_001070765 | *Danio rerio* |
|  | AAK61385 | *Gallus gallus* |
|  | NP_004066 | *Homo sapiens* |
|  | AAD46561 | *Mus musculus* |
|  | XP_786331 | *Strongylocentrotus purpuratus* |
|  | ANW48376 | *Euphausia superba* |
|  | AAU14170 | *Bactrocera tryoni* |
|  | NP_732407 | *Drosophila melanogaster* |
|  | ABB29886 | *Anopheles gambiae* |
|  | AAX58599 | *Danaus plexippus* |
|  | AAK11644 | *Antheraea pernyi* |
|  | AAY23345 | *Mamestra brassicae* |
| TIMELESS/TIMEOUT | AAR15505 | *Danaus plexippus* |
|  | AAF66996 | *Antheraea pernyi* |
|  | EFA04644 | *Tribolium castaneum* |
|  | AAY40757 | *Aedes aegypti* |
|  | NP_001157553 | *Mus musculus* |
|  | XP_002934575 | *Xenopus laevis* |
|  | XP_006565495 | *Apis mellifera* |
|  | NP_001265529 | *Danio rerio* |
|  | NP_722914 | *Drosophila melanogaster* |
|  | AGV28716 | *Eurydice pulchra* |
|  | EEZ99220 | *Tribolium castaneum* |
|  | EHJ76705 | *Danaus plexippus* |
|  | AAF54908 | *Drosophila melanogaster* |
|  | GAXK01195225 | *Calanus finmarchicus* |
|  | ANW48379 | *Euphausia superba* |
|  | GDUJ01099210 | *Talitrus saltator* |
|  | AUI80375 | *Euphausia superba* |
|  | phaw_30_tra_m.019341 | *Parhyale hawaiensis* |

Table S2: Search query accessions and results for putative *P. hawaiensis* housekeeping genes.

| Name | Full name | Accession | Contig | E-value |
| --- | --- | --- | --- | --- |
| EIF | eukaryotic translation initiation factor 5 alpha | ALD48732 | PH.c108324_g1_i1 | 3.00E-92 |
| GAPDH | glyceraldehyde-3-phosphate dehydrogenase | BAC77082 | PH.c123129_g1_i1 | 0 |
| RPL32 | ribosomal protein L32 | Q94460 | PH.k31.comp224_seq3 | 6.00E-49 |
| RPS18 | ribosomal protein S18 | AEB54647 | PH.k21.comp177_seq4 | 9.00E-76 |
| RPS20 | 40S ribosomal protein S20 | ALE99171 | PH.c89656_g1_i1 | 4.00E-60 |

Table S3: TMM-normalised expression values for putative *P. hawaiensis* circadian and housekeeping genes across 8 timepoints.

| gene_id | 09:00 | 12:00 | 15:00 | 18:00 | 21:00 | 00:00 | 03:00 | 06:00 |
| --- | --- | --- | --- | --- | --- | --- | --- | --- |
| bmal1 | 0.00 | 0.14 | 0.38 | 0.15 | 0.53 | 0.17 | 0.71 | 0.00 |
| casein kinase II A | 17.70 | 19.90 | 19.66 | 13.88 | 13.49 | 9.72 | 15.92 | 9.55 |
| casein kinase II B | 23.19 | 24.11 | 32.31 | 18.46 | 24.99 | 18.40 | 31.01 | 16.56 |
| cryptochrome 2 | 1.97 | 3.45 | 2.54 | 2.97 | 1.49 | 0.85 | 3.93 | 0.34 |
| circadian trip | 10.56 | 12.70 | 14.29 | 10.10 | 12.49 | 9.11 | 13.49 | 9.62 |
| clockwork orange | 0.00 | 0.64 | 1.15 | 0.22 | 0.20 | 0.26 | 1.05 | 0.00 |
| doubletime | 0.35 | 0.52 | 1.10 | 0.44 | 0.39 | 0.27 | 0.87 | 0.23 |
| microtubule star | 131.48 | 108.92 | 120.45 | 122.90 | 129.51 | 127.00 | 121.06 | 128.56 |
| neiire | 1.65 | 1.64 | 2.17 | 0.71 | 2.23 | 1.03 | 2.49 | 0.43 |
| nemo | 0.44 | 0.82 | 1.59 | 0.59 | 0.45 | 0.00 | 0.52 | 0.00 |
| pigment dispersing hormone | 31.34 | 51.73 | 36.24 | 62.45 | 66.28 | 67.45 | 56.12 | 62.93 |
| pdh receptor | 0.00 | 0.91 | 0.00 | 0.24 | 0.22 | 0.00 | 0.69 | 0.00 |
| period | 0.00 | 0.83 | 0.00 | 0.00 | 0.77 | 1.03 | 1.08 | 0.00 |
| protein phosphatase 1A | 9.08 | 15.38 | 18.94 | 5.51 | 13.20 | 4.17 | 19.93 | 5.05 |
| shaggy | 6.80 | 11.79 | 9.69 | 5.26 | 7.00 | 3.87 | 7.04 | 4.11 |
| supernumerary limbs | 1.05 | 1.53 | 2.95 | 0.68 | 1.54 | 0.28 | 1.84 | 0.51 |
| twins | 1.03 | 1.35 | 1.84 | 0.80 | 1.54 | 0.55 | 1.81 | 0.62 |
| vrille | 0.70 | 2.72 | 1.88 | 0.76 | 3.12 | 0.46 | 2.75 | 1.44 |
| widerborst | 5.10 | 5.39 | 6.16 | 7.24 | 7.86 | 5.75 | 8.51 | 4.85 |
| EIF | 550.94 | 487.54 | 551.86 | 523.31 | 513.04 | 430.07 | 630.65 | 644.93 |
| GAPDH | 1508.12 | 1167.41 | 1238.34 | 1419.71 | 1294.95 | 1578.77 | 1248.02 | 965.88 |
| RPL32 | 870.78 | 813.72 | 761.47 | 1073.89 | 1160.78 | 838.97 | 835.78 | 849.03 |
| RPS18 | 1512.55 | 1144.77 | 1444.89 | 1989.97 | 1559.80 | 1675.56 | 1547.46 | 1660.79 |
| RPS20 | 1997.26 | 1604.00 | 1653.89 | 2699.71 | 2139.38 | 2292.26 | 1956.35 | 2322.85 |

**Nucleotide and peptide sequences**

**Section A - core genes derived from genomic/transcriptomic resources**

***bmal1*/BMAL1**

Contigs from head transcriptome

>lcl|PH.k21.comp127566_seq0

CAGTAGCAGCGGCAGCAGTGGCAACTACCCCACCCTGCCCGCCGACCTCACCCGCCTGTGCCCCGGCTCCCGCCGCGCCT

TCTATGCACGCATACGCTGCCCCAGTGTCAATAAGGTGCAGTCTGATGACGGTGGTGGTGGAGGTGACAGTGGGAGCGTG

TGCGACGAGAGCATGACCGGGGACAAGCGCTACCTCAGCATACACTTCACTGGGTACCTCAAGAGCTGGCAGGGGGGCCG

GCGCGCCTCGTGTGGCGGAGGGGACGATGATCACGATTTTGGAGATGCTGCTTGTCTGGTAGCCATCGGACGTCTTCACA

GACCTCACGCTGACTTCCCTCCGCTGCATTTCATTGCCAAACTCTCGGCAGAAGCCAAGTACAGCTATGTTGATCAGAGA

GTGAGTGTAGTGCTGGGCTGGCTGCCACAAGAGATCTTAGGCGCGAGTGTGTTCGAACTGAGCCACCCCAGTGACCACTC

GACTCTCTCCGCTGCCCACCGTGCCTTACTTGGGAAGACGTGCATGGCGCAGTCTCTGCACTACCGTTGCCGGCACAAAA

ATGGCCGCTGGGTGCAGCTGACCTGTAAGTGGACGCTCTTCACTAACCCATGGACCAATGAGCTGGAGTACATCGTCGCC

AGTAATAGCGTATTGCCGTCCTCTCCTGCTCCCGATGACGCGCTGGCAGATGCCTCCTGTCGCAGTGCAGAGCCCGTGAT

ATCGTCCCCTACCGTGGCCAGCGTCTCGCTGGGTGGTGGGTCTTTGGATGTCCGCCCTGCTTCGGTCACCAGCTGCTCCT

ATGGCGGCGATGCCAAGGCCTTCACTGCTGGGAAGGACAACCTTAGCTCTGACGCAATTCGAACGCCCCATGACGGTAAA

GGAAGCG

>lcl|PH.k19.S23178339

TGACGGTAAAGGAAGCGGCCCTACGGCACGTGATGCCGCCGACTGCCGCACTGGAGCGCACCCACAGCTGTCGGCTGACA

CTGGCGAGACTGGGGACAACAGTGCACTGCCCCAGCAACTGGACGCCCCCACTCGCCTGCCCTTCCATCATCACTACAAC

ACCCGCAGTGAGTCGGAAGCAAGTGGCGTTGGCGAGACCACCAGCGACTCCGACGAGGCTGCCATGGCAATTATTATGAG

TCTCCTGGAAGCAGATGCTGGCCTGGGCGGTCCTGTGGACTTCAGCCATCTGCCTTGGCCTCTGCCCTGAGTGTAGCTGA

AGTGAGCGGTCCTATGGGTTATGTGGAGCAGAGACGCTCTGATGCGTCATGGATGGCCTTGTGTGTGCCTGCTACGTCAG

CTACTCGTCCAAAGAAGCAACAGGATAAGTTTTGTTGTTCCAAACTGAAGGCGACGGCTGTTACTGCTCATTCACGCTTT

CTGCTCAGCTAGTAACCAGAGTAGGCTACTGTACATTCAGGACAGAGCCATTTGGAAGTTAACTT

Peptide sequence identified from genome-inferred peptides.

>phaw_30_tra_m.024548

MYSTGGYSNTHAEYISECGSIASVASLSSDGIAMKKKIPGHGECHNEDDLECSKLARSSAEWNKRQNHSEIEKRRRDKMNTYISELSRMIPQCRSRKLDKLSVLRMAVQHIKMLRGSLNSYTEGQYKPGFVSDDEVQLLLKQECCESFLFVVGCDRGKILFVSESVAHILQYTQQELLGSSWFDILHPKDLNKVKEQLSCGDMNRRERLVDAKTLLPVHQSPNSSSSSGSSGNYPTLPADLTRLCPGSRRAFYARIRCPSVNKVQSDDGGGGGDSGSVCDESMTGDKRYLSIHFTGYLKSWQGGRRASCSGGDDDHDSGDAACLVAIGRLHRPHADFPPLHFIAKLSAEAKYSYVDQRVSVVLGWLPQEILGASVFELSHPSDHSTLSAAHRALLGKTCMAQSLHYRCRHKNGRWVQLTCKWTLFTNPWTNELEYIVASNSVLPSSPAPDDALADASCRSAEPVISSPTVASVSLGGGSLDVRPASVTSCSYGGDAKAFTAGKDNLSSDALRTPHDGKGSGPTARDAADCRTGAHPQLSADTGETGDNSALPQQLDAPTRLPFHHHYNTRSESEASGVGETTSDSDEAAMAIIMSLLEADAGLGGPVDFSHLPWPLP*

***clock*/CLK**

***P. hawaiensis*** – identified by alignment and expanded using mapped reads

Aligned sequences

LQNS02278184.1:5360739-5360901

AAGTCCCGGAACCTGTCTGAGAAGAAGAGACGGGACCAATTCAATGTCCTCATAGGAGAGCTCGCCGCCCTCGTCACCCCCGAGGGGGTCCCGAAGAAGATGGACAAGAGCACCGTCCTGAGGGCCACCATCGCCTTCTTCAAACAGCAGAAAGGTACGTTT

LQNS02278184.1:5362063-5362219

GCGGCACCGGTTCCAGTCGCGACAGAAGCAGTGTGGATCTGAACGATAATTCCAAGCCAAACTTCTTGACGAACGAAGAGTACACGCACCTCATGCTGGAGGTAAGTAGCACCGTCCTCATGATGGCAAACACTACCAGTTATGGAGTATTGTAAG

LQNS02278184.1:5403976-5404114

GTATTCGTCGGTATCTGCAGGCTGGTGCGGCCACAGCTGTTGAGGGAGATGCGTCTGCTGGAGCGAACCAACACGGAGTTTGTGTCCCGCCATAGCCTGGAATGGAAGTTCCTATTCCTGGACCAGCGGTCAGTTGCG

LQNS02278184.1:5406430-5406538

AGAGCTTCAGCCATCATCGGGTACCTGCCGTTCGAGGTGCTCGGTACATCGGGCTACGAGTACTACCACGTGGATGACCTGGAGCGCGTGTCCACGTGCCACCAGTTC

LQNS02278184.1:5409584-5409734

GTGATCCGCACCGGCAAGGGATCCTCCTGCTACTACCGGTTCCTCACCAAAGGGCACCAGTGGATATGGCTCCAGAGCCACTACTACATCAGCTACCATCAGTGGAACAGCAAACCGGAGTTTGTGGTGTGTACCAACACCGTAGTCAGG

LQNS02278184.1:5411511-5415480

AGTTACGATGATATCAAAGCCGAGATGCAGTCCAGCTCGTGTCCCACGCCGCAGAGTCCCCCTAATGGAGGAGACGAACCAACATCAGGACCCGGCTCAGTCCCAACCCCATGCGGGGCAAGTGGTCCTGACGGGCTCCAAATCCAAAAACCTTTCTCCAAGGCGGACAAAAAGACCAAGTTCACGTCGTACAAAGATCAGGGatataaacccgacgaaaaatcTCCGGTTGTTGAATGCCAGTTTCAAACGCAGCGTCAAATTCAAAATTGTAGTAGAAATCAGCTTTCTACTATCCATGAAGAGAGGCTCAAGGAATCCTCAACTCCGTTTTCAGTCAAAACAGAAATGGATCAACAGATGCAAAGCAGTAGTTCAACGAAAGAACCTACAATGAATTATGCCCCCATTAGTGACGTATCATTGGAATTTGAGTTTGGGTACAAAGATGCTGAAGACGCCAAGCAAAATATGCAGTTTCAAACTAAGAGTATACTCAAGACCAGTCCTCAGCAAACCCAGATGGTTCGACCAGGGGGAGAATGTTTCCAACCCAGTAGGTCTTCTACAATCGCTCAACAGCAAGGGACAGAAAGCCAAAGATTGATGAGCGAGTCCATGGATTACCAAACTCCATGTCAGAATTACGTGAATGACCAGTCTCCAAGAGGCGTAAGGAAGACCGAAAGCGATCCTCTCCTCAGCGTCAGCATTGAGGAACGGTGTCAGGACTCTAAACTGGCTCACCTTGATATCTCCAACGCAGTTTCAGACAGCCAGGTAAAGGACTTCGATGTACTGAACCAGGGACAGTCGCTTGCTTTTGATATCGACGACGACTCTCTACCCACGAATCTTCTTGATTTGGGCAAGAAAGATGACGGACTGATGACTGGAGCGATCCCGTTAGAAGAAGGCCCGAGCTATGAAACGCTCCAGCCTCTGGACTTCGAAGCCATAAAGTCAGATGTTGATGGTCTTGATCCGACTTTCGGTGATGTTACGGCAGATGATTTCCTGAAGCTTGACGCTGACGCCAACCAATTTTGCCAACCCAGCGTACTCATCGGAAACATTATGGGCAGGGGACCATGCACGGACGAGGGCAACAGTCGAGAAAGAGAAGTCAAAGTACCCTGCAGTACAGCGTCTACAACTGATGCTTGTGAAGCTAAAGAACGTATATCTCCTGTCAAATCTTACAATACTTTGCAGTGCGTGACGTCTAGTAACGATGTTTTTGCTAATCGATCTGAAAAGCCACCAGTCCCAGAAGCTACCGTCTCttcaaatcagttgaaaaactgCGCTAACCAGAAGGGTAAACCCAACACTTTTCAAGCACTAGAAGGGCCTTCTAAGCTAAGAAATCTGCTCGAACGAAATTTATATGAGAACCCTGGAGAAAAAACTGCTAACATTCATATGAAACAAAATCAGAAGCTGAAGAAGAGCAAGACGGATGGTAAAAGTGAAAGACATCCATCGATGAGTCAGCTTGATTCTCCGTCTCTTCGAAAAGACCGACACCAGCAGCAGGTGAAACTTCAGGGCCCTCCTGCTGAGCAGCACCATTCTGTTTCCTCTACACAGATGCAGAACGAAATCCAGATTCACGGGCAAGCTGGTAATCGAGAGAGACAAGCATTTCGAGGAGGAGCTCAGCCTGAACAGATCTACCAGGCCACTGAAGCAAAACCAGCAGTAAACAACAGTTCATCTTATATGGgaggaaatattttattgaatcaGTTGCAAGGATTTAAGGAATCCAAACAAAAAAACCAAGTTTCTTTTCAGGATGAAAGCCACCAGAATAGACACACCAGCACCAACAGATACGACGAACGCCATAAGTTTCGTATGGAACATTCGGGAAGTGATAAGCGAACTTTTGGAAGCCCATCTCAGGCTGGCAGTCGGCCAAGGTTTGTCACTGTCAAAGGGCTTCAAAATCAAGGAGGATACACGAAAATTACTCAGAGTCAGGTGCTTAGTCGAGGAGGTCATTGTGAAGGCCAGTCTTCTGCCAAAGTGGCTCAGAATCAAGGATTTTGTGCTCAATATAGTCATCCAAGCTCAACTAGAACAGATAACATTGATCCAACGCAGCTCCCCGGCTATTCGAGCAATCCGGTGCCTGTCACTCAGAGTTGGCGAACAACAAACATAGGTATCGCCACTGATCCGAACGAAACTTCACGAAAAATTAAACAATGCGAATCCAAAGTCAATGTGAATAAAGAAAAACGACCCCAGCAGATGGCGACTGCACTGAATCCTGACTATCAACAGTTCCCTCATGAAATCaatcagcagcaacagcagcagcagtttaaGGTTCAGCAAGGTTGTGATACTCGGCAATACTCTCAACTAAATGAACGCTACTCCGTCCAGCAGAACCAAGCTTATGGACAAGCTAAACCCCAAGCGAACCAATCTCTTTACCCTGTGAAAACTTTGAGGGACGGCGGCAAGGTTGTTTTGGATCATCAACCCCATCCGCAGACTCTCAATGAGTCTTGTGGGATGGGCAGTAATATCCCTCCACAAAACCGTTTTCAGGGAAATGAAGCAATGGCAAATTACTCTCAGTTTATACCTGGAGTCTATAGCTATCAAGATGTCCAACAGGGAGCTGCCTTATGTTCATCATTCAACGTGATGCACGTGAGACCAACAGTAATGGGAGGAACTCAGTATCTGGCCGGTGAGATGCCCCAGAACCTGGCCCATGAGCTTTATGGATCAGGTCTCCCTCACCATCCGGTTGCAGGGACGGGTATGCAGATCCATAGTCAGAATACTACAATATACACCCAGCCAGTGGCTCCGATGCTGAACGGTGGGGATGTGCTCATACAGGCCGGTACGTCTTATCCTGCAGATAGGGCTGTGGGACAAAGCCCTTCCGTAGTTTGTTCGACTCAATCTATAGTCATGCAGCCCATGGCCAAACCCACAGTAATTTCATCCAACCGCTGTCAGCATCCCAGTATGCATCCTTTCCTTCAGAACAAGTCTGGTGGTCAGAACATGGTTCAGACCCAAGATAGTCAACATGGTACATATACAGACCAAGATTCAAATGCCTGTGTAAACAAACAAGGTGATTCTCACTTGCAGCCTCAAGAAATGACCCAGTCTCAAGTCGCAATGAATGAAACTTGCAAGATGTCTGCAGAAACTGCCCAAACAGGTGCTGTTACTCATCAGGTGACCGATCAGTCTAGGAACTTTTCCCAAATGACAATTCATTCACAGGATGAAACTTCTAAAGGTGCTGATGATTTTGGGTTTCCATCTAGAGAAAATTTTTTCCAAGATCTAGCCGACACATTTCAGTCCATCCAAGGTGGCAGAAATGAAAAACCCATTGTGGATAACCTGGTTGAACAATATGGATCTGCTCAAAGCTCCCGCAGCCTACTTGAGAGCGCGAGGAAATCCCCTTCCAAAAACATGGGTTATAATCCAGCAAAGAAACACTCCAAAAATAAATCAGAGAAACACCCCAGAGGAGGAGACGCTCCATCTCAGTCGCACTCGCATGCGTCGAATTCATCCGTAGCTAATGCAAGTGACCAAACAAAATCTTCTGACAAGTCTAAAAATTCGATGAATGGACTTTCCAAACTTCAGCAGCTAGATTCCATTCTCAGTTCTTCTTCCAGACACGTGACTTCCTCGACATCGGACTCGAAATGTTCGTCGCTTGAGTTCTTCCTCGAAAGTGAGCTTGGAAGTGGCATTTCAAACCAAGGTGGACCTAATTCAGCAATGGACAGCGGCACTTATCCAAAGTCAAATGGGTCCAATGTGGATGATATACTTCCGTATCAGAACACTTTGGACCCAAGCGACGTGATGAAGGCTGATTTATCGAATATTACTGGGAAAGAGTTTCTTCAGCAGTCATCTTTCCATCACCTGCCAAGACTTTGGGTATCGAGCAAGCATCGGCGA

LQNS02278184.1:5417295-5418813

GGTTCCCCTGCCAAACCTTCTTCTTCTGGCATCAGGAAACAGGGACTTGCCCAACCCAATCGGCACGGAGACTCTTTTGGATTCAATCCAAAATCTACTACTGATCAGACTGGAGTACAACCCAtaatagaaataaatgaagaCTGCCTCAAACTAAACGAACGTGGTGGAGGTGACATCAGCACTTCCAGCGAACTTCAACGGAAACAAACTTCCAATAAAGCTGCAAATCAACCAGGGACGAGCCAAGTTGATAACCAGTTTCGTAAGTACCCAATAACTCAACCAGGAATGGAGGTTACCAATGGTTCAAGACCAGCAGAATCGACAAGCAAGAATAAACCATACAACAAAATAGACGAGTTTTGGTTCCGTCAAGAAGGTACCAATAGGACAGCTGTATCAACATTTGAAGGAATGACGAAATGTGCTTCCCAAGAAACGATTCCAGAGAACGGAGGCAACAGTTTGCAGAAACAAAAGTTGAACTCTACAATTCTGTCGGATCGCTTGTCTGGTTCTTTGCCGGAGATTCCCATGTCTAGTTTCAATTCACCATTTTCACACGTCTTCTGCATAGCTGAAGTTGAACCGGCTGTAATTCCAATGTCTACAGATATCGGCTCGTCAGTTTCTTTCAGGGAACCTACCACCACTCAAAGTGGATCTCAATCAATTTCTGAAACGCCTTCTTTCGGAACCTTTCCCTCAGGAGGGAAGCCATTGTGTACCGCAAACATTTTCCATGCGTCTTCTTCAGCTGTAGTCCAGTCTCCAATGTCACCGATGGTTCACGGTGTGCCTTCCCCCATGATACAGTCCCCTATATCTCCTATGGTGTGCCCACTACCCTCCTCGACGTACGTCTACTCGCCGATGTCCCCGAAAACCAGTTCAATGAAATCCCTACCGTCCCCGATGGTACAGTCCCCTCTGCCGTTCGTCCAATTTATCCCGACCGTGAGTGGCGTCAACCAGTCGATGTCTCCAGTATCTCACACATCTGCTCCCCACCCAAGGAACTGTTCCTGTGGACACTGCGGAGTATCCCCGTGCCCCAGTCGGTCATCATCGGTCGGACCCTCTAACAACAGGATCACCGATATCAAGCGAATGAACTCCATCAACATCCCTGCGTCATCCGCAGCTGATTCATCATCGCAGACCTCCTTCCAGAAGCGGATACCTACTAGTTTCTTTCCAGGCATTCCACAGCGACCTAACTTTTCTGGCTCCGATGAAAGTATCTCAGGTAAAGCTTTGACCTTCCAGAAAGGAATCCCATCCGCAGAGGGTACCTGGAACGGATCCCATATCGAAGTGGAAGCCCCTGGAAAGAAAGCGAAGGACTCATCATACGTCACTAGGTCCAGCTCTTTTCCATCCCCCAGAGCGGTGTACACTGTGCCTGCTGTGGTGTCCCGAGGCTCCATCAAGACCTCGACTCCTGTGAGCGCCAATGTGAATATCATCGGCCAGCCGTCGGTGGGTCGAAAATCTGTCGCTCGACAGCTGCGT

Constructed peptide

bHLH

PAS

PAC

PolyQ

KSRNLSEKKRRDQFNVLIGELAALVTPEGVPKKMDKSTVLRATIAFFKQQKGTFGTGSSRDRSSVDLNDNSKPNFLTNEEYTHLMLEVSSTVLMMANTTSYGVLVFVGICRLVRPQLLREMRLLERTNTEFVSRHSLEWKFLFLDQRSVARASAIIGYLPFEVLGTSGYEYYHVDDLERVSTCHQFVIRTGKGSSCYYRFLTKGHQWIWLQSHYYISYHQWNSKPEFVVCTNTVVRSYDDIKAEMQSSSCPTPQSPPNGGDEPTSGPGSVPTPCGASGPDGLQIQKPFSKADKKTKFTSYKDQGYKPDEKSPVVECQFQTQRQIQNCSRNQLSTIHEERLKESSTPFSVKTEMDQQMQSSSSTKEPTMNYAPISDVSLEFEFGYKDAEDAKQNMQFQTKSILKTSPQQTQMVRPGGECFQPSRSSTIAQQQGTESQRLMSESMDYQTPCQNYVNDQSPRGVRKTESDPLLSVSIEERCQDSKLAHLDISNAVSDSQVKDFDVLNQGQSLAFDIDDDSLPTNLLDLGKKDDGLMTGAIPLEEGPSYETLQPLDFEAIKSDVDGLDPTFGDVTADDFLKLDADANQFCQPSVLIGNIMGRGPCTDEGNSREREVKVPCSTASTTDACEAKERISPVKSYNTLQCVTSSNDVFANRSEKPPVPEATVSSNQLKNCANQKGKPNTFQALEGPSKLRNLLERNLYENPGEKTANIHMKQNQKLKKSKTDGKSERHPSMSQLDSPSLRKDRHQQQVKLQGPPAEQHHSVSSTQMQNEIQIHGQAGNRERQAFRGGAQPEQIYQATEAKPAVNNSSSYMGGNILLNQLQGFKESKQKNQVSFQDESHQNRHTSTNRYDERHKFRMEHSGSDKRTFGSPSQAGSRPRFVTVKGLQNQGGYTKITQSQVLSRGGHCEGQSSAKVAQNQGFCAQYSHPSSTRTDNIDPTQLPGYSSNPVPVTQSWRTTNIGIATDPNETSRKIKQCESKVNVNKEKRPQQMATALNPDYQQFPHEINQQQQQQQFKVQQGCDTRQYSQLNERYSVQQNQAYGQAKPQANQSLYPVKTLRDGGKVVLDHQPHPQTLNESCGMGSNIPPQNRFQGNEAMANYSQFIPGVYSYQDVQQGAALCSSFNVMHVRPTVMGGTQYLAGEMPQNLAHELYGSGLPHHPVAGTGMQIHSQNTTIYTQPVAPMLNGGDVLIQAGTSYPADRAVGQSPSVVCSTQSIVMQPMAKPTVISSNRCQHPSMHPFLQNKSGGQNMVQTQDSQHGTYTDQDSNACVNKQGDSHLQPQEMTQSQVAMNETCKMSAETAQTGAVTHQVTDQSRNFSQMTIHSQDETSKGADDFGFPSRENFFQDLADTFQSIQGGRNEKPIVDNLVEQYGSAQSSRSLLESARKSPSKNMGYNPAKKHSKNKSEKHPRGGDAPSQSHSHASNSSVANASDQTKSSDKSKNSMNGLSKLQQLDSILSSSSRHVTSSTSDSKCSSLEFFLESELGSGISNQGGPNSAMDSGTYPKSNGSNVDDILPYQNTLDPSDVMKADLSNITGKEFLQQSSFHHLPRLWVSSKHRRGSPAKPSSSGIRKQGLAQPNRHGDSFGFNPKSTTDQTGVQPIIEINEDCLKLNERGGGDISTSSELQRKQTSNKAANQPGTSQVDNQFRKYPITQPGMEVTNGSRPAESTSKNKPYNKIDEFWFRQEGTNRTAVSTFEGMTKCASQETIPENGGNSLQKQKLNSTILSDRLSGSLPEIPMSSFNSPFSHVFCIAEVEPAVIPMSTDIGSSVSFREPTTTQSGSQSISETPSFGTFPSGGKPLCTANIFHASSSAVVQSPMSPMVHGVPSPMIQSPISPMVCPLPSSTYVYSPMSPKTSSMKSLPSPMVQSPLPFVQFIPTVSGVNQSMSPVSHTSAPHPRNCSCGHCGVSPCPSRSSSVGPSNNRITDIKRMNSINIPASSAADSSSQTSFQKRIPTSFFPGIPQRPNFSGSDESISGKALTFQKGIPSAEGTWNGSHIEVEAPGKKAKDSSYVTRSSSFPSPRAVYTVPAVVSRGSIKTSTPVSANVNIIGQPSVGRKSVARQLR

KSRNLSEKKRRDQFNVLIGELAALVTPEGVPKKMDKSTVLRATIAFFKQQKGTFGTGSSRDRSSVDLNDNSKPNFLTNEEYTHLMLEVSSTVLMMANTTSYGVLVFVGICRLVRPQLLREMRLLERTNTEFVSRHSLEWKFLFLDQRSVARASAIIGYLPFEVLGTSGYEYYHVDDLERVSTCHQFVIRTGKGSSCYYRFLTKGHQWIWLQSHYYISYHQWNSKPEFVVCTNTVVRSYDDIKAEMQSSSCPTPQSPPNGGDEPTSGPGSVPTPCGASGPDGLQIQKPFSKADKKTKFTSYKDQGYKPDEKSPVVECQFQTQRQIQNCSRNQLSTIHEERLKESSTPFSVKTEMDQQMQSSSSTKEPTMNYAPISDVSLEFEFGYKDAEDAKQNMQFQTKSILKTSPQQTQMVRPGGECFQPSRSSTIAQQQGTESQRLMSESMDYQTPCQNYVNDQSPRGVRKTESDPLLSVSIEERCQDSKLAHLDISNAVSDSQVKDFDVLNQGQSLAFDIDDDSLPTNLLDLGKKDDGLMTGAIPLEEGPSYETLQPLDFEAIKSDVDGLDPTFGDVTADDFLKLDADANQFCQPSVLIGNIMGRGPCTDEGNSREREVKVPCSTASTTDACEAKERISPVKSYNTLQCVTSSNDVFANRSEKPPVPEATVSSNQLKNCANQKGKPNTFQALEGPSKLRNLLERNLYENPGEKTANIHMKQNQKLKKSKTDGKSERHPSMSQLDSPSLRKDRHQQQVKLQGPPAEQHHSVSSTQMQNEIQIHGQAGNRERQAFRGGAQPEQIYQATEAKPAVNNSSSYMGGNILLNQLQGFKESKQKNQVSFQDESHQNRHTSTNRYDERHKFRMEHSGSDKRTFGSPSQAGSRPRFVTVKGLQNQGGYTKITQSQVLSRGGHCEGQSSAKVAQNQGFCAQYSHPSSTRTDNIDPTQLPGYSSNPVPVTQSWRTTNIGIATDPNETSRKIKQCESKVNVNKEKRPQQMATALNPDYQQFPHEINQQQQQQQFKVQQGCDTRQYSQLNERYSVQQNQAYGQAKPQANQSLYPVKTLRDGGKVVLDHQPHPQTLNESCGMGSNIPPQNRFQGNEAMANYSQFIPGVYSYQDVQQGAALCSSFNVMHVRPTVMGGTQYLAGEMPQNLAHELYGSGLPHHPVAGTGMQIHSQNTTIYTQPVAPMLNGGDVLIQAGTSYPADRAVGQSPSVVCSTQSIVMQPMAKPTVISSNRCQHPSMHPFLQNKSGGQNMVQTQDSQHGTYTDQDSNACVNKQGDSHLQPQEMTQSQVAMNETCKMSAETAQTGAVTHQVTDQSRNFSQMTIHSQDETSKGADDFGFPSRENFFQDLADTFQSIQGGRNEKPIVDNLVEQYGSAQSSRSLLESARKSPSKNMGYNPAKKHSKNKSEKHPRGGDAPSQSHSHASNSSVANASDQTKSSDKSKNSMNGLSKLQQLDSILSSSSRHVTSSTSDSKCSSLEFFLESELGSGISNQGGPNSAMDSGTYPKSNGSNVDDILPYQNTLDPSDVMKADLSNITGKEFLQQSSFHHLPRLWVSSKHRRGSPAKPSSSGIRKQGLAQPNRHGDSFGFNPKSTTDQTGVQPIIEINEDCLKLNERGGGDISTSSELQRKQTSNKAANQPGTSQVDNQFRKYPITQPGMEVTNGSRPAESTSKNKPYNKIDEFWFRQEGTNRTAVSTFEGMTKCASQETIPENGGNSLQKQKLNSTILSDRLSGSLPEIPMSSFNSPFSHVFCIAEVEPAVIPMSTDIGSSVSFREPTTTQSGSQSISETPSFGTFPSGGKPLCTANIFHASSSAVVQSPMSPMVHGVPSPMIQSPISPMVCPLPSSTYVYSPMSPKTSSMKSLPSPMVQSPLPFVQFIPTVSGVNQSMSPVSHTSAPHPRNCSCGHCGVSPCPSRSSSVGPSNNRITDIKRMNSINIPASSAADSSSQTSFQKRIPTSFFPGIPQRPNFSGSDESISGKALTFQKGIPSAEGTWNGSHIEVEAPGKKAKDSSYVTRSSSFPSPRAVYTVPAVVSRGSIKTSTPVSANVNIIGQPSVGRKSVARQLR

***H. azteca*** – identified by alignment and expanded using mapped reads

Aligned sequences

JQDR02011873.1:37346-37502

CCCAGGAAGTCGCGCAACCTCTCCGAGAAGAAGCGACGCGATCAGTTCAACCTCCTGATAGCCGAGCTGTCCGCCCTTGTCAGTCCGTCCGCCGCCAAGAAGATGGACAAGAGCAGCGTCCTCCGGGCCACCATTGCGTCCTTCAGGGACCAGAAG

JQDR02011873.1:37821-37887

AATGAGCGCTTCAAGCCGAGTTTTCTGACCAATGAGGAGTACACACATCTAATGCTCGAGGTGAGT

JQDR02018666.1:25-133

AGGGCATCCGCCATCATCGGGTACCTGCCGTTCGAGGTGCTGGGCACGTCAGGGTACGACTACTACCACGTCGACGACCTGCACCGCGTGGCTGAGTGCCACCGCTTC

JQDR02018666.1:2140-6303

TTTGCAGTTATGACGACATCAAGGCAGAACTCGAAAAGTCTCCCCAGAATTACAACGAAAACTCCAATCAGAGCTCGATGAGTGACGACAAAACTGCGACAACCTTAAGTACTTCTCAGGTCAACGCCGCGTGTGGCGACTCCGACCTACGGAATCTTGGAAGCAATAGTAAAACCAGTTCTTACAATGTGGATTGTGGTAATAATGGCGAGCATGAAAAGTTAACTTACCAAAAGGAAAGGGTGGAAGATTGGACCCGAGATGATGGGCGTATGGGAGGAACGGAGAATCTCGCATCTGACCCCTTTGGATGCCATTTTGTGACCAATACCTGTTCTTCTTACTTTGAGGAGAGGTCTCTGCTCGCAGCGCAAGACAGGCAAGCTGCAGCTGAAATGGGCATCGTGGTCAAAGCGGAAACTGATATGATTAATCAAACCACCACCCCAAAAACTGAATCAAACGTGAGAGAGCAAATGGGTAACCCTGCTACACCTCATGACGTTACTTTCTTCTCAGAATACTCTGACAACATTCAACACCAAATGAAAAGCCAAAACAAAATCACCGAACGAGACTCGCAACAACTGCAACCACCTTACCAGGATACGACAATGAATAAAAGAACGTCGATGGAGCAATCCCTTTCATCGGCTGATAGAAATAAACCTTCTATCAGAGTGAACCGATGCCACCAACGGTGCTTAGGGGAACCTATGGAGTTTCAAAATAACcgacatcagcagcagcaacaccaacaGCGTAAGCAGCAGCATCCACAACAACACCAGCAGAATGCACAGCAGCAACACCAACAGCAAGCAAGCAACGCTGTTTGTTCAAATCCTTATGGAAATAACTTCGGAAAAACGGAAAGTGATCCCATGCTCTTGAGCGGAAGTGTCGATCCTAATGACTCGAGTAACATATTCTCAGACATAAACTTTGCCAAAACTGATAACAACATCCCACCATTAACTTTTGAAGATCTCAATCCCATTGAATCAATTTCTTTTGACATCGGTGTTACTTCCTTGAACACGAACATCTTGGACcttgaaaacaaagaagaaaacttaatttcagGTGCCTTGGGCCTAGAAGTAAGCAGCGGGTACACAGCTCTTGATTTCGATGCTATCAAAACAGACGGGGAAACATTTCCCGATGCTTTCGGCGATTCTGGCAGCGACGATTTCCTAAAACTAGATATAGATTCCAATCAGTTCTCTCAACCGAGCGTGTTAATCGGTAACATTATGGGCAGAGGACCACGCATTGAAAGTAAGAACATTAAAAGTAACCTCTCTGGTGATGAATATGGAAACCAGCCGATACCAAACAAAAACGTGTCTCAAGACAAAAAGCTGTATGGAACTCTCGAGTCGTTGACGCCGGCCAACAAGACTTATCAAGAACAAGTACTCAACGTTTTCGCTGAAGATGAAACGTCGTCGAGTCAAAACGCCTCTAATGAAGGTGGCAGGGACATTCCCAGAAAATCCTGCAATCAGGgtgaaaattcaaaactaaGGAACCTACTTGAGAAAGACTTGAAAGAGCCAACAAAGGAAAACCTCAAAGACTCTTCAAACAAAAAAGATCTTTGTAATAAGAAAAGTAGGATCGATCATAGGGAGAACCATGCAGCGCAGCATTCAGAGTCTGTCAAACATCATCAGGCAGAGTCATCGTCGTGCAGGCAACAACTGTCAGATGTGCAAGAAACGATCAATTGGTTTCCAAACAAGCAGAATCAACAAATGCAGCCGCAGCAGaaccaacagcagcaccagcaagaTCATCAGCCAACGCATCAGAACCtccatcaacagcaacaacgcCACCATCAGCAGCAGAACCCACAGCATCAACAACAGACCAACCAACAGCTAAAGAcccaacaacagcagcagtcaCAAATCGTGCAGTTTCAAGACCAGCATCACGACTCTGGACTTGACAATCCTGGTCAGTACACGGTGAAGTCCAACTCAAGCAGGCATGAACGACATGAGAAACTACTTCTAGAAGATGAATACCCGATCGTCAAAGAAGAAAATGATAGCAGCCAAGTAACTTCGGATAATAATCAGCGGGAATGCACACGCCACAACAACCCTGTTACAGGATTCCACCTTACTTCCTCGCAGCAACAACACCAGGAAGTAATTCTAAATTCAAAACAGCTAACAAAATTTGACTCCAAGCGAGGAGACAAACAGACAACTATACACGAAGGCAATGAAATGCCTGATCTTCGTTCCCACGAGCTACAAGAGTGGAGTAATAAACGCACTGGAGCTGGAAACTTTGTTCAGTGCAATCCACAAACCCTCGTTCATGAGGAACAATGTTGGTCAGATGCTTACCAGGGCAAGAAACCAACTAAAACAGAACTGGGATGGACACCTAAGCTTGCGGAAACAACAGCACCCTACACCAATAAGTCTGGGAAGCAAAGCCAACTGCAGTCTTTCCGTGAACACATTACTGTAGCTCGAAACAGCCGTGAAAATTCATACCAGCAACGACAGCAGGAACACATTCAAGCTCGATTCACGTATCAAGTACATGAAGATAGGAAATTTGAACCTCAAATAACGCTCCAAGAAAGCCAAAGGCGTGCCGAACAAAAGGGCATTGGAGATCTGATTGTTGAAAATCATCTCGACAGTAACACCTCGATTCATAGGCAAGTTGGAAGTGATCAAAGATTGAACTACACCTTATCCCACTATATTCCAGGTGTGTACAGCTTCCCTGAAACATGTGCTGTACCGCAAGAGGCTACACTGTGTTCTCCTTACAACGTGATGCATATGAGACCTACAGTTCTTGGAGGAGCTCCTTTCATTTCAAACGAAGTTCCCCAAACCCTTTCACGCGAGTACTACACGGCTTGTATTCCAGACCGTGCGATCGGCACTGGATTGCAGATACATAGTCAAAAAAGAACGATCTATTCCCAGCCCATAGCCCCTCTGCAAGGTTCCAACAGTGAGAGTACCATGATTGCCAATTATGGTGGCGATGAACAAGATTCAAGACAAAACGGAATGGACAAATGTACTACCAACCCTGTCGACCAACCTGATCATTTTCAGGTGGATGGTCGTAGCGTGACTCAACTCCCCAACACCTACCTTGACAAGAATATGAACCAATTCTCTCTCAACAAAGCAGTCGCAACAAATCAAGATAAAAAAGGTTACTACGCTAGTCTTACTTGGCAGTCATCCAATCAAAACGTAGCTCAGCAACTGGGTAAAAACTCAAATGTTGTGGATCTCTACACTCAGGCTCAGTTCCTAGTCAACCAGCGTCCATCCCAGAAAATGAACGCTACAATGAGTCGTAAAAACAACGTACAGGCTGACGCCAACCAAACATACCCTTCCGAGGCACAAGGTGTCGGACACCAGATGAACGACCAGTCACATAACTTCTCTCAGATGTCCATACATTCCCGGAATGATTCTAAGCATCATAAACGCTCGAAAGAGAAAGGTATAAGTTCAACTAGTGACTTCTTCCAGGGGTTAACCAATCCCATTGAGAACGTCCAacaggaaaaattttcgaacaatCTACGCAACTTTAACGAGTTAACGCAGAGGCAAAATCAAGATGGATATCGACAGTCTAACACTGACAACTACTATCACTCCAAGGGCTCAAAACGGGAGCACAGagaacaggaaaaaaatatgtcTGAAAGGCTCCACAGTGGAACAGAAATGACTTTGCAATCGCATTCACATACTTCCAACTCTTCAGTAGCAAACGCAAGTGACAAGACAAAGTCGTCGGACAAATCAAAACATTCCCAGCAAGGATTTTCCAAACTCCAACAACTTGACTCTATTCTAAGTTCTTCCTCGCGTCATGTAACTTCTTCCACTTCGGATTCCAAGTGCTCATCCTTGGAGTTTTCCCTCGAAACTGAATTCGTGGGTGGGTCTAATCAACAGACGGATAGCCAAGCAGACACGCAATCAAGACCTAATCATCGAGATTACGGAGACCGACCAAAGACGTCAACCAACCTCGATACGAAAACGGACCAGAAACATGATTTGTCTGAACTTTTTGGCGAAGAAGAGTACGCATTGCCAAAGCATCTCCCTAGACTCTGGGTGTCCAACCGACAACGGCATCAA

JQDR02018666.1:6572-8806

ATGCACACCGAAAGCACCGCACCAACAAGCAGAAGAAACGTACCAAGGATGACAAAGAAAGCTTCTCCGGTCATCACTCTAATAACCTGCAGCAGGAAAATTCGCCAAACAAAGAGCGACCGCCCTGTGAAGATGGACAGGAACGTCAAAACAGGACAGAGGGTAACGAGTTCTGCAAAAGGTTAGAGACATCCAAATCTTTCCGGTTCAAAAATTCGAGAGAGAACAACATGTCAGCCGGCGTCTCTTGTTCTGATTATTCAAAAAAGGATGAGTGTGAAAAAGCAGTTCAAGGCAGTTTTTCGCTGTCAACTTCAGTAAAGTTTCCAGACGCTCTTTCGTCGGACGACCAAATTAACAATGGATCCCAATGTGGGCAAGCTGGACGATCTACCGAAACAGGGGGAACCCAGTGTCATCTAGGTGGAGGTAACGTTGAGAGTTTGAGACCGAAGCAACAAATCTGTGAAAAGAAGATGTCGACAGAATCTAAAAACCAGTCTCTCAATGGTAACGAATTGCATCCACCGAAAAGCAGCGACATTACCGTTCAACAAGATGGAGTTGGGCGAATCACCTATTCTCAAGATGAACCCGAGATGAAAGACGTGACAATATGCGCCGGGTCTCAAAATTTTTCTACCATAAATTTTGAGGAGTAcgaaatgatgcaaaaatgcAACACTTTTTCACCTAGCAAAGAGACACCAGGTCGAACAGAACTGCAACGAAAGTCTCCAACTGCATTCAGTACCGCCACTTTTGAAATCCAACCATCATTCACAAGGAAGGCGGGTGGAGTTGTCCAAAATTCAGAAACATCACCATGTGAATTAGAAACTTTTGTCAAGTGCGGTGATACACTCGATAGTTTTTTGAGGCATCGAAATAGAGTTGAAGAGAACCAGCAAAACTCATATCAATGCAATTACCAACCAGGAACGGATTCCACACTCAACGACATGCAGAAAGCCCGAAATCATCAGGTCCCAAACGTGGACCAAAAACCTGTGCTATGTTTGGCACAGGGGCAAATGGGTGGAGCTATTAACGCTTCTTGTTTTCGAAGAGCGACCATCGGCAACAGTTCCCTCAGGTTCGATTGTGTGGTCAAAGCATCGTCACAAGAAAGTATTCCTGAAAATGTTAATGCAAAAGTTTCACAGAAATATCGAATGCCGTTGGGTGGTCTCCAAGATCGTTTGTCAGCATCGCTCCCAGAAATACCGATTTCCACACAAACGGGGCTGCCAATTGCACGAATCTTCTGTGTCGCAGAGCGGGCTAATGAAAACTTGAATCCCGTATCGAAGAATTTAACTCAGCAAACTTTCTACACAACCCACGAGCAAGCGACTGCAACGCAGAACTCAAACAAACTTCCTGAAAATTCATATGGATGTTTCATCTCTCCAACTGAAAGAAAGCCTATTGGTACAGCTTTTTTTCATATGCCTTCTTCGCCGGCTATAACCCCTTCTCCAATGTCCCCGATGATTTATGGGGCACCGTCACCGATGGTCCAGTCGCCAACATCACCCATGGTTTACCCTCTGGCATCATCAGCTTACGCATATTCTCCAGTTTCTCCTAAAACTAGCTCAATGAGGTCTCTGCCGTCCCCTATGGTTCAGTCCCCGTTGCCTTACGTTCAGATAATTACCGCGGTGAGTGGGACTACACAGCCAATGTCTCCAATGGCACACAACTCGGCATCACACCCAAAAGCCTGCGCTTGTGGATACTGTAAAGCTACATGTTCTCCATCCCGATCTTCATCCGTGGGACAATCAATTAAGTCGTCTTATACCCGAAGGTTAAATTCAAACCAAGCCGTTGGGGTATCGACTCCAGTAACTGATTATGAGCACCAGAACAACTTTCAGAGACGCGTGGGTAGTTATTTCCACACTGCTAAGCTGAATCGAGGAGACGACCCTACTCTGCAGGTTGACacgggtggtggtggtgctttTGGGATTCAGCGGAATTCAATCACAAGTAATGACCAGAATAGAAGCCCATCGTTTGGACCCGGGATTAACATGCCTAGCCGTGAAATAAGAGAAGTGGTCATGTCACGCTCGAGTTCGTTTCCTTCGCCAAAACGGCTTCAACACATATCAGCTATGGCTTCCGAAGTGGCTTTGGATTCCACTCCTTCTGTTAACGCCAATTTGAATCCCGTAGGTCAGAATACCATTATAAAATCAGCAGTAGTTGGAAAACATCGTTGAAGAAGATTGATTTCAAGAAGACAACtccattatcaaatattttattgcgtcctaacttcaataataatttaagtcaTTTAATGTAACGTAAACTCAGAAAGATGAATAggtcatactgtccaaataatgactcaTTTTCTTACCCACGGTGCGAGCATCGGGCAACACTGTTACTCCATTCTATGAGCCAAAATGTTATCAGTTACTCTATCTTATTTTTTGTGGGTCTAGATGACCCAGACAGGCGCAAAAGAGTttatcaaaatttggacaaataCTCACTAATTTCTTTCCCTGAAACATCCACAGAGACTTTGTGTAGTGGTAGCTTCACCGAAATGAAGTGAGATTTTCGTGGGTTCGAGCTTCGGGAAGCTCAGTAGTGCATTTCACGATGTCACTGTTGGGCGGTGCTTGGTAGCCGTCAATGAGTGGTGCATCAGCATGGTCTTGTGTTGGTACTCTCTAGGTACTTGTTACCTAGGGAgatattaaattctttaaatccACCCCTGTTTATTAGTAAAACTTCTTCACTTCAGTTTCCACCGTTTTTATTCCTCTTAACTAGTGGAAAATGacgaaaatgttttgaataacttatttgtgtttgaacttgaatgaacaattgaaaaaaactttttattgtagtttatttgtttactgGGAGCCACTTCGTTGgatttaaatcatattatacgatgaaataaatttatattgttcacGATGATATAAACGAGGTTTAGAGGCGTTGCGAAAAGAATAATAtaccaatatatattatttaatctacTATAGATACATTTAAAACATAGATATAAATAAGTAGATaagaaagtatttttattagtagatattagatataaattttatcgtaCGACCAATTATGATGAATTACAATGTGATGTTGTCAGAATTTGGTATCCTGAAAGTGCTGTAGGAATTTGGTGTCTTGAATGTCCTGtaggaatttggtgtcctgaaagtcctctaggaatttggtgtcctgaaagtcctgtaggaatttggtgtcctgaaagtcctgtaggaatttggtgtcctgTTTATTGCCCGGCCAGTTTAACCCTCTTAACCTGAAGATATAACCTGAAACTAACCACCCGCAACCCTCACGGCAACGTCCAGCTAACCTTGCAGGTTAGAATTCTGACTTCAAGTTAACAAAAAGTATAATTATAGGGCATTCGTCAATATTAACTACTAAGATTTGCTAGTagtaattataatgatgatgaatattaatacttttaattataattattaatgtgctGAATGTGTAATGTGATGTACTGTGTGTTAACGTATCTTAATGAGGTAGCTGTTCGGGTACCAACAATCATGTACACTAGCTCCATGTTCACCCactatatcaaaatattatatttaatccaaaattttacctctAATCCAAAACTTTGCTCTTAAATCccaaattttttcctcaatacCGAATTTTCCCCCTCTAATccgaaatttttacttacaatccaaaattttaccctctaatccggaattttcattcaaa

Constructed peptide

bHLH

PAS – PAC

PolyQ

PRKSRNLSEKKRRDQFNLLIAELSALVSPSAAKKMDKSSVLRATIASFRDQKNERFKPSFLTNEEYTHLMLEVSRASAIIGYLPFEVLGTSGYDYYHVDDLHRVAECHRFCSYDDIKAELEKSPQNYNENSNQSSMSDDKTATTLSTSQVNAACGDSDLRNLGSNSKTSSYNVDCGNNGEHEKLTYQKERVEDWTRDDGRMGGTENLASDPFGCHFVTNTCSSYFEERSLLAAQDRQAAAEMGIVVKAETDMINQTTTPKTESNVREQMGNPATPHDVTFFSEYSDNIQHQMKSQNKITERDSQQLQPPYQDTTMNKRTSMEQSLSSADRNKPSIRVNRCHQRCLGEPMEFQNNRHQQQQHQQRKQQHPQQHQQNAQQQHQQQASNAVCSNPYGNNFGKTESDPMLLSGSVDPNDSSNIFSDINFAKTDNNIPPLTFEDLNPIESISFDIGVTSLNTNILDLENKEENLISGALGLEVSSGYTALDFDAIKTDGETFPDAFGDSGSDDFLKLDIDSNQFSQPSVLIGNIMGRGPRIESKNIKSNLSGDEYGNQPIPNKNVSQDKKLYGTLESLTPANKTYQEQVLNVFAEDETSSSQNASNEGGRDIPRKSCNQGENSKLRNLLEKDLKEPTKENLKDSSNKKDLCNKKSRIDHRENHAAQHSESVKHHQAESSSCRQQLSDVQETINWFPNKQNQQMQPQQNQQQHQQDHQPTHQNLHQQQQRHHQQQNPQHQQQTNQQLKTQQQQQSQIVQFQDQHHDSGLDNPGQYTVKSNSSRHERHEKLLLEDEYPIVKEENDSSQVTSDNNQRECTRHNNPVTGFHLTSSQQQHQEVILNSKQLTKFDSKRGDKQTTIHEGNEMPDLRSHELQEWSNKRTGAGNFVQCNPQTLVHEEQCWSDAYQGKKPTKTELGWTPKLAETTAPYTNKSGKQSQLQSFREHITVARNSRENSYQQRQQEHIQARFTYQVHEDRKFEPQITLQESQRRAEQKGIGDLIVENHLDSNTSIHRQVGSDQRLNYTLSHYIPGVYSFPETCAVPQEATLCSPYNVMHMRPTVLGGAPFISNEVPQTLSREYYTACIPDRAIGTGLQIHSQKRTIYSQPIAPLQGSNSESTMIANYGGDEQDSRQNGMDKCTTNPVDQPDHFQVDGRSVTQLPNTYLDKNMNQFSLNKAVATNQDKKGYYASLTWQSSNQNVAQQLGKNSNVVDLYTQAQFLVNQRPSQKMNATMSRKNNVQADANQTYPSEAQGVGHQMNDQSHNFSQMSIHSRNDSKHHKRSKEKGISSTSDFFQGLTNPIENVQQEKFSNNLRNFNELTQRQNQDGYRQSNTDNYYHSKGSKREHREQEKNMSERLHSGTEMTLQSHSHTSNSSVANASDKTKSSDKSKHSQQGFSKLQQLDSILSSSSRHVTSSTSDSKCSSLEFSLETEFVGGSNQQTDSQADTQSRPNHRDYGDRPKTSTNLDTKTDQKHDLSELFGEEEYALPKHLPRLWVSNRQRHQAHRKHRTNKQKKRTKDDKESFSGHHSNNLQQENSPNKERPPCEDGQERQNRTEGNEFCKRLETSKSFRFKNSRENNMSAGVSCSDYSKKDECEKAVQGSFSLSTSVKFPDALSSDDQINNGSQCGQAGRSTETGGTQCHLGGGNVESLRPKQQICEKKMSTESKNQSLNGNELHPPKSSDITVQQDGVGRITYSQDEPEMKDVTICAGSQNFSTINFEEYEMMQKCNTFSPSKETPGRTELQRKSPTAFSTATFEIQPSFTRKAGGVVQNSETSPCELETFVKCGDTLDSFLRHRNRVEENQQNSYQCNYQPGTDSTLNDMQKARNHQVPNVDQKPVLCLAQGQMGGAINASCFRRATIGNSSLRFDCVVKASSQESIPENVNAKVSQKYRMPLGGLQDRLSASLPEIPISTQTGLPIARIFCVAERANENLNPVSKNLTQQTFYTTHEQATATQNSNKLPENSYGCFISPTERKPIGTAFFHMPSSPAITPSPMSPMIYGAPSPMVQSPTSPMVYPLASSAYAYSPVSPKTSSMRSLPSPMVQSPLPYVQIITAVSGTTQPMSPMAHNSASHPKACACGYCKATCSPSRSSSVGQSIKSSYTRRLNSNQAVGVSTPVTDYEHQNNFQRRVGSYFHTAKLNRGDDPTLQVDTGGGGAFGIQRNSITSNDQNRSPSFGPGINMPSREIREVVMSRSSSFPSPKRLQHISAMASEVALDSTPSVNANLNPVGQNTIIKSAVVGKHR

```
cryptochrome 2/CRY2
P. hawaiensis –identified by de novo assembly of head transcriptome reads. Sequence below generated by alignment of reads to genome and assembly of transcript with Stringtie (below).
Genome coordinates

LQNS02278089.1	StringTie	transcript	4842778	4859580	1000	+	.	gene_id "HEAD.92040"; transcript_id "HEAD.92040.1"; cov "5.055981"; FPKM "0.248153"; TPM "0.338697";
LQNS02278089.1	StringTie	exon	4842778	4844273	1000	+	.	gene_id "HEAD.92040"; transcript_id "HEAD.92040.1"; exon_number "1"; cov "3.626181";
LQNS02278089.1	StringTie	exon	4844509	4844617	1000	+	.	gene_id "HEAD.92040"; transcript_id "HEAD.92040.1"; exon_number "2"; cov "6.593780";
LQNS02278089.1	StringTie	exon	4846689	4846831	1000	+	.	gene_id "HEAD.92040"; transcript_id "HEAD.92040.1"; exon_number "3"; cov "4.827119";
LQNS02278089.1	StringTie	exon	4847533	4847717	1000	+	.	gene_id "HEAD.92040"; transcript_id "HEAD.92040.1"; exon_number "4"; cov "4.462806";
LQNS02278089.1	StringTie	exon	4848885	4849114	1000	+	.	gene_id "HEAD.92040"; transcript_id "HEAD.92040.1"; exon_number "5"; cov "6.340151";
LQNS02278089.1	StringTie	exon	4849556	4849723	1000	+	.	gene_id "HEAD.92040"; transcript_id "HEAD.92040.1"; exon_number "6"; cov "6.672047";
LQNS02278089.1	StringTie	exon	4850439	4850579	1000	+	.	gene_id "HEAD.92040"; transcript_id "HEAD.92040.1"; exon_number "7"; cov "6.084380";
LQNS02278089.1	StringTie	exon	4852901	4853052	1000	+	.	gene_id "HEAD.92040"; transcript_id "HEAD.92040.1"; exon_number "8"; cov "4.383372";
LQNS02278089.1	StringTie	exon	4853978	4854180	1000	+	.	gene_id "HEAD.92040"; transcript_id "HEAD.92040.1"; exon_number "9"; cov "5.939760";
LQNS02278089.1	StringTie	exon	4855505	4859229	1000	+	.	gene_id "HEAD.92040"; transcript_id "HEAD.92040.1"; exon_number "10"; cov "5.112722";
LQNS02278089.1	StringTie	exon	4859428	4859580	1000	+	.	gene_id "HEAD.92040"; transcript_id "HEAD.92040.1"; exon_number "11"; cov "12.333298";

Assembled nucleotide sequence

>HEAD.92040.1 gene=HEAD.92040
gATAATACATGCGTTGCTTCAGTCGTAAGACATAGAACTACTGCGTAACTTGCAGGTAACCCATGCAGTA
AATCATGACTTGAGAAGTgctatcatttatattttttcttggttGAGCGTTGGAATATTCTTGCAATGGG
CGAGCGCAATGACCCTGAAGTTTCGATCCCAGAAAGTTCTGTGCCCGCTCCAGTGACCGGGGGTCAGGTT
GTGATGGTCCATCCATCGCGGCAGTCATCGGAGGACATGAATGTGGTGGCCAGCAGGTCTTTCATCTCCG
AACGATTTGTCCCCAATCGACTGACTTCAGACTACCCCTCGATGTCATCGCTACCGATCATGACGAACAA
GGTGCTGACCGGGACCGTGTTCATGACTGATCCCAGTACCATGACTCAGATGACGATGGTTGGGGACTAC
AAGCTGGAAAGTAAGGCGCTGATCCCTCAGAGTATGCTGGACGTTCATCCAATGTATGGGGACATCAAAA
GCATCCAAGGAAGGGCATTCATCGGTGAGCAGATGTGCTCTGTGCAGCCCATGGTGACAGACGCCTTGGT
GCTGTCGGGCAGAACAATGGCACCTCGCTTGATGCCCCCAAAAGCATCTCCGATCAAGGACGATATGATG
ACGGGCATTACGATGATAAGTGGCAAGCCGCTCTCAGGTCGAGTTATGGTTGGCGATGCCAGGGCAGTTA
CCACACGGATCGTTACATCCGATAAAGCAACTCCAGTAAAGACTACCACTTGTGAGAAAGCGTCTACCAA
TCGACCTACTGATGAAAGAACAACGCCAAAGACGTCTTGTGCCACAACCGTTCCGAAGGTCCATGCTGCG
ATTGAAAAGCATCCTCCTTCCAAATTGAGCCAAATTGACAGGATGACTGCTGCTCGCATGGTTGGGGACC
AACCCCAACAAGTTAGACAAGAAAAAAAGCAGACGATGGCTTACAAGACCGGAATAAAAAATTATCCCAT
TGAGAAAGGTAACCGAACTGTAACTCATGAGAAGCAACAGCTCTCAAAGAATGATTCTGGTGAAAAGCCG
ATGGTGAATGAGCGATCATCAACTACCCAGCCGGAAGTCAGCCGTGGAGAGCACTTCAACTTTGGATGCG
ATAGCAACGCTGGGCCTAAGTTAAACTCGATGACCCTCGAAAGAACCAAGTCTCCGTACCAGCTTGCGGA
ACCAAAGATGTCTCCTAGGAAGATCGCATCGCCGGGAGAACGCTCGGCGCAACGGACTATTCTAGGGGAC
AATAGCGTCGTCCAGGGAGACTGCACAGGAGTTTATGGATCTTCCCTGCCGGCAGAAAAGAAACGCTGTC
GTGTTACTCCTGGAAAACACGTGGTACACTGGTTCAGGAGAGGACTCCGACTCCATGACAACCCTGCTCT
GAGGGACTCCATCATTAACTGCGAGACATTTCGCTGCATATACATCCTGGACCCGTGGTTCGCTGGATCT
TCCAATGTCGGAGTCAACAAGTGGAGGTTTTTACTGCAATGCCTGGAAGATCTCGACAATTCTTTGCGCA
AGCTCAACTCCCGCCTGTTTGTTGTACGTGGTCAACCAGCCAACGCTCTGCCGCAGCTCTTCAAAGAGTG
GAACACAACAATCCTGAGTTTCGAGGAAGACCCGGAGCCGTTTGGGCGCGCCAGGGACACCAGCATCATC
GCCATAGCGCAGGAACTTGGCATAGAGGTCATTGTCAGGACCTCCCACACGCTCTACAAGCTCGACAAGA
TAATCGAGAAGAAAGGAGGCAAGCCGCCCCTCACCTACAAGACCTTCCAGAACATTCTGGCGATGATGGA
CCCGCCCCCAGCGCCCGTCCGTCCCGTAGTGGTGGACGACCTCAAGTTCGCCTCTACCCCCCTGCAACCC
GACCACGATGACAAATACGGGGTGCCAAACCTCGAGCATTTGGGTTTCGAGACGGACAACCTGCCCCCGG
CGGTGTGGAAGGGCGGGGAGACGGAGGCCCTATCTCGGCTCAAGCACCACCTGGAACGTAAGGCGTGGGT
GGCCTCCTTTGGCCGGCCCAAGATGACCCCGCAGTCGCTCTTCGCCTGTCCCACGGGCCTGTCTCCGTAC
CTGCGCTTTGGGTGCCTGTCTGCCCGCAAGTTCTACACTGAGCTCAACGAGCTCTACATCAAGATCAAGA
AGGTACCGGCCCCGGTATCGCTTCACGGCCATCTACTGTGGAGGGAGTTCTTCTACACCGCCGCCACCAA
CAACCCTAAGTTCGACCATATGAAGGGCAACCCCATCTGCGTGCAGATACCTTGGGACAAGAACCCTGAA
GCTCTCGCCAAGTGGGCCCATGGACAGACAGGGTTCCCGTGGATAGATGCCATCATGATGCAGTTGAGGA
AGGAGGGATGGATCCACAACGTGGCCAGGCACGCCGTCGCCTGCTTCCTGACCAGGGGGGACTTGTGGGT
GTCTTGGGAGGAAGGCATGAAGGTGTTCGATGAGCTGCTGCTGGACGCCGACTGGTCGGTCAACGCTGGC
TCCTGGATGTGGCTGTCCTGCTCCTCCTTCTTCCAGCAGTTCTTCCACTGCTACTGCCCCGTGCGGTACG
GGAGGAAGGCAGACCCCAACGGGGACTTTATACGGGCGTACCTGCCTGTACTAAAGAACTTCCCCACGAA
GTACATCCACGAGCCCTGGAAGGCCCCGGAGGCGGTCCAGCGCACGGCTCGCTGCCTCATCGGCCAGCAC
TACCCGCTGCCTATCGTGGACCACGCGACCCAGAGCCAGTGCAACATCGAGCGTATGAAGCAAGTGTACC
AGCAGCTAGCCCACTACAGGGCCAATGCGACGTCTCGCTCCTGCAGTGACACGAAAGGCTGCTTCAAGTC
CTCCGGTTCGGGTCGCCCCTTGACTGGAGGTCGAATGGTGACGACTGTGTGAAGATTACGCGCTTGACAT
ATTAGGGTGCGGTGTAGTCGCTCTGTGTGATGCGCaatcttatatttttatatttgctgtTGAAGTATCT
ATTAATGTCTTAGTGTTATTGAATATGATTAATACCTTTTCCGCGTCCGAACGATGAAATTTTCTCACTC
AACTGCTGCCATTGCCTCCTAATTCCTTGGTGAATTACTGTTAAATTTATTCATGATGTAATTTGAGTAG
TTTCTAATTAATTTGTGAAGTGTTGTAATTTTCGTTGATGGATTTTTGATGAATAGTGGTGAATTACAGT
AAATGTGCTACTAACATTGTGACAAGCTAATTCTCTTGCTGTCAAATTTAGTAATAAATTGTACTCGAAT
TTAGTGATTCAATTTTGTCAGATTCATTAAATAGTTATAATATACTTTAAGACACTGTTATATTACCGCT
GAATTTACTGATAACTCGTGCCATATTTCACTATTAAATTTACTGATAATCTCCGCTCAAATTCAAATTC
ATTATTCCTGCATTTAGTGACAAGCTGAACTCGATTTCATTGATCTATTACAGCTGAATTCAGAAATGAA
TTGTACTGGAATTCATTGTTATTTTGTAACCAGCTTAACTGCTTCAGACCCCTCCTGATCAAACCTGCAA
TTTGTGCACTTTTGGGGTAGCGACTGTCAATTTACTGTTTCAATATAAACAACTTTGAGGCTTGTTGCCT
ATTTCTATTTCCAACTTTAATGAATAGCACTGACACCGAAATCATTAAAAAAGTATCTcatgttttattt
aatttcaaatgtacCAATCACCACATTGCTGCCTGTAATGGAGGAAGCCTCATCAAAAGCCTTGACTGAA
GCCTGGAATATCTTTATCCGTCCCTAAAATTGCTTTTAAACGTATTACCCTAATTCCGAATCAAATACTC
TTTTCTTTTGAGTAGCTAATTTGTACGTATATAATAAACATCCCAATCATTTCTAATTCCAATCCAATCG
GAGCTTCTCCATTTTGATCAGCAATATTGACCTGATATACAGCCTTGGACAAAATTATGAATCGccctaa
aaacagtaaaaaaaatctgcttAATAATATGTACTTGATTAATTATTCCCCAAAATACTAAGTTATAGTA
TTAAGTATGGTAGCATCTTATACTTAGGATGGCCTCCTTTAGACTGTAACAGCTTTTTCGCGTTTTGGCA
TACTAGAGTTTAACTAGCACAAGTGGCTGCAACTAGCCCTCCTTCATCAACTTTGGTATCTTATTGCTGT
TTTGAGAGACATTAAAGCATCCTGATTCGTTTCCCTATGTGTGACCACACATGCTTGTTGGCTCCAGGTC
GGGCGACTGAGAGGGCCTCTGAAACCAGCCCCTCGAAACTCTCGATGTGCGACATGAGGCATTCCTTGTC
CTATTGGAAGGCATCTTCGGGAATTTTATTAtcgaatattttcttcagtaactggctgtacactctgtcc
aacatgtcggtgtacgttgctgaattcattctaCCCTCGCATGACTGTGTacagcattccagtccctgaT
GACGCCATGCAGCCCCAAATCATAACGCTGCTTTCCCCAAACTTCACCGAAGGCACAACGCATTGTGGGT
AATAAACTTCTCTAACTCTTCAACGCTCATAAACAGCGCCAGGTGTTTCAAAAATTCTGTAAATAGATTT
TTTAAAGACATGTTAAAATCACCAAAGCggtaatttacaaaaataaaatgttgaaaGTGACCGTGCACGT
TTGGTTTGATAAAACATCAAAGAAACTCAACTCAAAGGTGTTTTCACCAAACGTAAGCTCATCATCACCA
GTCTAATGTTCATGAGCTTTTGCAAACAACTGTCGTATCTTCTTGTTGGCATCAGGCAGCAATGGTTTTT
CCTAGGTTTGCATGATTTGAGACCAAATTCTACACGTCTTTATCTCACCGTACGCGATGAGATTTCTTTG
CGAGTTGCCTCCTTAAACTAGGCTTTCAGATTTTCAGCAAGACTCTCTTTATCAGTGTGTCCTCTCTTGA
CGTCGTATACCTTTTTTTGGCCTAAAACTGCACGGTTTTTGAATGTGTTTGTCTTCCTGTACCATTTGAT
AGCAAGTTGGATACAATTCCGTGAACAtttcattttttagcaatttcattttgagaatgaCCAATATCGA
ACAATGTAACAATTTGAgccatttttttcacaaaatttctttTGTGTGACAGTTCTTTTTAAACCAAtgt
actaaataataaaaaataaaaaaattatgttatgcAAATGAAGAAAAGTATTAAATGTGGTGAATTTGCT
CAGACAACTTACACGCGATTTGCTGCAGCCACTAAATTAACTGGTTTAAACAGTGCTTTCGTAAGAAACT
ATCTAACTTCCCACGACGAATAAAACTGTTGAcgtaaaaaatgtaaacgtttcgggattaatttcccatc
ttcagaactaaaaatcagagatgtaaacgtttcgggattaatttcccatcttcagaactaaaaatcagag
atgtaaacgtttcgggattaattttccatcttcagaactaaaaatcagagatgtaaacgtttcgggatta
actTCCCATTTTAAGAACTAAAAATCAGAGATGTAAACGTTTAAAATGAGTAATTTATTTCTATTGTACT
CTCTAGAGATTAGGGACTGAGATGTTAACGGAGGAGGTGTAAGAGGCGAATGACCGGAGAAAGCTTAACC
GATAACATCTAGGTGCCTAGATAgcatcatataaataaattattagttATAGGCGTTTACATCCTTTATT
TTGAgttctgaggatgggatattcatcccgaaacgttaacatgtTTTTTATCGACACTTCTATTCATTAT
ATGAAAttgttcttagtttaatatttaagtTCTCTAACTTTGTCAATTAGATTGTAATTATCAGACAATA
CAATCATCGGACATTACCTACACAGGTCAAATCTAGTCAGAtatttctaaaaacaaaattttcctcGGCG
CTTCATGGTTTTGTCCTTGGCTGTATAAACGTTTTGAACTTTAGGTCTCAAATTTCTCAGATCTACTGCA
AAAGCTCTTCCCTATGAATTTTTGGCAGCGTCAAAACGCCTTTTCACGATTATAACGTCTGGAGCGGCCA
ATAATTTTGTCGGCAAGATCGtattattagtatatatattacattaattATTCATTCATGAACTGTAATG
AGAAAAATGTTGTCCTAGAACCAGGACACTTTTCCGCCCGCAAACAGTTGCGCCAGTTACCTTGGCGTCG
TGGTGCAGCACGCACTCTATAAATTTTCGCACTGTACCAACGCGTCAGGGCAAGAACAGTAACTTCTGCC
TCTGAATAGGTAAGCGAAACAACATTTTTCTCAATCATGTCACTGTAGCCTATCTTGCGACTGGCAGCAG
CTTTGATATTGCGAGGGGTGACGACAATGGCAGTCTAATTTTCCGATATGGTACCGACAGCAGTAGCTTG
ATATTCCGACATGGTGGAGGTAATGGCAGTTTGATATTCCGACATGGTGGACGCAATGGCAGTTTATTAT
TTCGACGGGGTGACAGTAATGGCAGTTTGATATTCCGACATGGTGGAGGCAATGACAGTTTGATATTCCG
ACATGGTGGAGGCAATGACAATTTGATATTCCGACATGGTTGACGCAATGACAGTTTATTATTTCGAAGG
AGTGACAGTAATGGCAGTTTGATATTCCGACATGGTGGAGGCAATGACAGTTTGA

Peptide
DNA photolyase
FAD binding
MGERNDPEVSIPESSVPAPVTGGQVVMVHPSRQSSEDMNVVASRSFISERFVPNRLTSDYPSMSSLPIMTNKVLTGTVFMTDPSTMTQMTMVGDYKLESKALIPQSMLDVHPMYGDIKSIQGRAFIGEQMCSVQPMVTDALVLSGRTMAPRLMPPKASPIKDDMMTGITMISGKPLSGRVMVGDARAVTTRIVTSDKATPVKTTTCEKASTNRPTDERTTPKTSCATTVPKVHAAIEKHPPSKLSQIDRMTAARMVGDQPQQVRQEKKQTMAYKTGIKNYPIEKGNRTVTHEKQQLSKNDSGEKPMVNERSSTTQPEVSRGEHFNFGCDSNAGPKLNSMTLERTKSPYQLAEPKMSPRKIASPGERSAQRTILGDNSVVQGDCTGVYGSSLPAEKKRCRVTPGKHVVHWFRRGLRLHDNPALRDSIINCETFRCIYILDPWFAGSSNVGVNKWRFLLQCLEDLDNSLRKLNSRLFVVRGQPANALPQLFKEWNTTILSFEEDPEPFGRARDTSIIAIAQELGIEVIVRTSHTLYKLDKIIEKKGGKPPLTYKTFQNILAMMDPPPAPVRPVVVDDLKFASTPLQPDHDDKYGVPNLEHLGFETDNLPPAVWKGGETEALSRLKHHLERKAWVASFGRPKMTPQSLFACPTGLSPYLRFGCLSARKFYTELNELYIKIKKVPAPVSLHGHLLWREFFYTAATNNPKFDHMKGNPICVQIPWDKNPEALAKWAHGQTGFPWIDAIMMQLRKEGWIHNVARHAVACFLTRGDLWVSWEEGMKVFDELLLDADWSVNAGSWMWLSCSSFFQQFFHCYCPVRYGRKADPNGDFIRAYLPVLKNFPTKYIHEPWKAPEAVQRTARCLIGQHYPLPIVDHATQSQCNIERMKQVYQQLAHYRANATSRSCSDTKGCFKSSGSGRPLTGGRMVTTV
H. azteca – identified by alignment of reads to genome and assembly of transcript with Stringtie, and with reference to previously reported H. azteca CRY2 (accession XM_018166003).
Genome coordinates:
KV721600.1	StringTie	transcript	195083	208564	1000	+	.	gene_id "HYA.34936"; transcript_id "HYA.34936.1"; cov "2.705847"; FPKM "0.149709"; TPM "0.284359";
KV721600.1	StringTie	exon	195083	195104	1000	+	.	gene_id "HYA.34936"; transcript_id "HYA.34936.1"; exon_number "1"; cov "0.370370";
KV721600.1	StringTie	exon	195138	196627	1000	+	.	gene_id "HYA.34936"; transcript_id "HYA.34936.1"; exon_number "2"; cov "1.442456";
KV721600.1	StringTie	exon	197820	197928	1000	+	.	gene_id "HYA.34936"; transcript_id "HYA.34936.1"; exon_number "3"; cov "2.417941";
KV721600.1	StringTie	exon	198099	198241	1000	+	.	gene_id "HYA.34936"; transcript_id "HYA.34936.1"; exon_number "4"; cov "1.221445";
KV721600.1	StringTie	exon	198739	198923	1000	+	.	gene_id "HYA.34936"; transcript_id "HYA.34936.1"; exon_number "5"; cov "3.305706";
KV721600.1	StringTie	exon	199562	199791	1000	+	.	gene_id "HYA.34936"; transcript_id "HYA.34936.1"; exon_number "6"; cov "2.889855";
KV721600.1	StringTie	exon	200228	200395	1000	+	.	gene_id "HYA.34936"; transcript_id "HYA.34936.1"; exon_number "7"; cov "3.038691";
KV721600.1	StringTie	exon	200651	200791	1000	+	.	gene_id "HYA.34936"; transcript_id "HYA.34936.1"; exon_number "8"; cov "3.297872";
KV721600.1	StringTie	exon	202412	202563	1000	+	.	gene_id "HYA.34936"; transcript_id "HYA.34936.1"; exon_number "9"; cov "2.299342";
KV721600.1	StringTie	exon	203176	203378	1000	+	.	gene_id "HYA.34936"; transcript_id "HYA.34936.1"; exon_number "10"; cov "3.211823";
KV721600.1	StringTie	exon	203908	208564	1000	+	.	gene_id "HYA.34936"; transcript_id "HYA.34936.1"; exon_number "11"; cov "3.101782";
Assembled nucleotide sequence
>HYA.34936.1 gene=HYA.34936
GGCCAATGTCATACAATGAAATCTTTTTTTCACAGATTTGAGTGCTTTATATATCCTGTATTATTATTGT
CTTCATTACGTCTTGAAAATCGTGCCCCAATTTTACGTCTTATCTAACTTGCGTTTTGAGGTACAATATT
TTGTCTCAATGAGTGAGAGGAACGAGCTCCCGCCTATGGCAACTTCAAGTGAAGAGCCTCAGGCGCCGCC
AACGACACAGGCAGCAGTTTCCCGCGGGCCGCGCCAGCGACAGGACAGTTTGGACGTCGTCTTGAGCAGA
TCCATGGTAGCTGAGCGGGTAATGCCGGCTAGGTTTTCGTCAAATTACCACGCCATGCAGTCCATGCCGA
TGGTAGCAAACAAAATTCTTACCGGAACAATGATAATGACTGACCCAAATGCAATGACCCACATGACGAT
GTTTGGGGACTACAAACTGCCAGGGAAATCTCTCATCCCGCCGAGCATGATGCAGGTTAGATCGATGTAT
GCAGACATCAAGAGTATACAAGGACGAGGCAGTCTTGACGATCAGATGTACACAATGCAACCCATGGTAA
CCGATGGCTTAGTTTTACCTGGAGGAATGCTTACACACCCGTTAATGACTTCTAAATCTTCTATGGTCAA
GGATGACATGATGACGGGTATAACTATGATCAGTGGTAAGCCGCTCTCTGGCAGAGTGATGGTGGGGAAT
GCACGCGCGGTCACCACCCGCGCGGCCACAACAGACAAGACCAGCCCTACCAAAACACCCAACTGCGACC
GTGTCCTACAAAACCGTCCGCCCGAGGATCGAACGATTCCGTGTCCGAAAATAATCGGACCCTCCGTTGC
ATCAAAGCCCCATGGCATGGGAGATAAGCATCCACTTTCAAAACTTCGTCAAATTGATAGAATGACCGCT
TCCAGATATGCCAGCGAGCAGTCCCAAGGTCCTTCATCGGATAAAAAGATTCCTGACAATCACAAGACTC
AATTTAAAAGCTTCCCTGAAAACAGCTCTCGAAATCTCAAAGAAGAATTTGTAGTAAACACTAATGATTC
ATTGGGAAAAAATATCCAGTCTGCCGAACGCGAATCAGAATTCGTCTTCGGTAGAAATCCGAAGTTTGGC
AGCGAAGATTTCGGAACTTCTACAGTCGACTCAATGGCACTAGATCGGACCAAATCCCCAGGCCAAAACT
CTTCTTTTGGCCTCAAGATGTCGCCGAATAAAACAGTATCCCCAGTGATGAGATCCACCCATCGTACCAT
ATTGGGAGACAATAACTTGGCACAACCTGAGTGCAGTGGGTCTAGCGGTCCAAGCATGTCTTCAGAGAAA
AATAGTTCGACCATCAAAGTTCGTTCGGGGAAGCACATTGTTCATTGGTTCCGTCGAGGCCTTAGGCTAC
ACGACAATCCCGCTCTCAGAGACTCTATATTCAATTGTGAAACATTTCGCTGTATATATATTCTCGACCC
GTGGTTTGCTGGCTCCTCTAATGTGGGAGTCAATAAGTGGAGGTTCCTCCTTCAGTGTTTGGAAGATGTC
GACAACTCCTTGCGTAATCTCAACTCCCGTTTGTTTGTGGTACGAGGGCAGCCTGCCAACGTACTGCCTC
AACTTTTTAAGGAATGGAACACGACTGTGCTGAGCTTCGAGGAAGATCCTGAGCCCTTCGGCCGTGCGCG
GGATGCCAGCATCATCGGCATCGCCAGGGAGATGGGAATCGAGGTTGTCGTACGCACATCCCATACGCTC
TACAAGCTGGACGAAATAATCGAGAAGAAGGGAGGGAAGCCGCCTCTGACGTACAAGACGTTCCAAAACA
TCCTGGCTATGATGGACCCTCCGCCTGCTCCTGTGCCACCTATCGTGGCGGGTGATCTGCAGCACGCGTT
CACACCCATCCAGCCCGACCACGATGACAAGTATGGCGTGCCCAACCTCGAACATCTTGGCTTTGAAACG
GAACAGCTCCCGCCGACTGCGTGGAAGGGCGGTGAGACGGAGGCACTTCAGCGTCTCAAACACCACCTGG
AGCGGAAGGCTTGGGTGGCCTCCTTCGGCCGTCCCAAGATGACGCCGCAGTCGCTCTTCGCCTGCCCTAC
AGGACTCTCCCCTTACCTTCGCTTTGGCTGCCTCTCAGCTCGAAAATTTTACACTGAACTCAATGAACTT
TACACTAAGATCAAAAAAGTACCTGCACCAGTTTCGCTTCACGGCCACCTACTTTGGCGGGAGTTTTTCT
ACACGGCTGCCACGAACAACCCAAAATTCGATCACATGAAAGGCAATCCAATCTGCGTCCAGATACCGTG
GGACAAGAATCCTGAGGCCCTGGCGAAATGGGCTCATGGCCAAACTGGCTTTCCGTGGATCGACGCGATC
ATGACTCAGCTGCGCGCGGAGGGTTGGATCCACAACGTGGCACGGCACGCGGTCGCCTGCTTCCTGACGA
GGGGCGACCTCTGGGTGTCGTGGGAGGAGGGCATGAAGGTATTTGACGAGTTGCTTCTGGACGCCGATTG
GTCAGTGAATGCGGGATCGTGGATGTGGCTGTCGTGCTCCTCCTTCTTCCAGCAGTTCTTCCACTGCTAC
TGCCCTGTGCGCTACGGCAGGAAGGCTGATCCCAACGGCGACTTTATACGAACCTACTTGCCGGTTCTTA
AGAACTTCCCGACGAAGTACATCCACGAGCCATGGATGGCGCCGGAGAGCGTGCAGAGCAGCGCGCGCTG
CATCATCGGACAGCACTACCCGCTGCCGATGGTCGACCACGCCACGCAGTCCCAGACAAACATCGAGCGC
ATGAAGCAAGTCTACCGCCAGCTCGCCCACTACCGAGCCACCATGTCATCAAGGTCCGGTGGAGATTCGA
AATCACGCTGCAAGCATCAACAGACTTCAGCTCCCAACAGGAGCCTCTGCAGCAATCGCGTTCTTACAAC
TGTTTAAATAGTAACAGGACTTTAATTAAGTTAGGATTGGGTTAACAACAATCAGCGGTTCCAGAGCATC
GGTAACATGGAAAGATTTCATATGCTTGTATTAAGCTACTCTGGAACACTCATCATTCAATTCTgaaaca
agaattttttcacacTATCTGATTACTGGAAACAAAAATGGGAGATGTGGTTAAATAAAACTAGATTTTT
TTCGATTGGATTATCCATTAAAAGTGGGTATGGCGTTTTTTGATGTACGGAAATTTGTCCTGTTAGTCCA
CAGTCAAGAATAACTCTGAAATGTGGtgattaattactttaattatgaGTTGAGTCGCTCGGGGGGCTGT
TGTATTGCTTTCTCCCTCTGTTGTATTTCACCTCAATTTGTGGGCGATTTATTTTCCCGTACAGCAAGAA
GGATGAAGATTCCTGTAGGAATGAAACCATCTAATGTGACCTTAACTATGAGATAGAATGttttaggaaa
ttaaaatttgtgcaaaTACCAAATTTACCGGATTaacttttgtaaaataataatgccAGAATGTTTGTCG
CAGTATCAACGGCTTACCTGAAATTAATTGGACGTTTAAACATCAGGCTTCTTAGAATATAAAGAATGTT
ATTTAAGCGCTCCAGTGGAGCTGAAATGTTagatgataaattcaataaaacccTTTACTTCTGCCGTGTC
TGTGTCGTTACCTGCTGGAGGTTGTAGCGAGGGCTTCTTGGCAGTCATCGTGGGTCTGGTAGAGGCCAAC
ATACCGCCTGCCCCAGAGAGCGACTTGATATTATAAGATAACTTGTCAAACATCGGAAAACCTTTGATTA
TATTAGGACGGAAATGAAATAACTATACTAAGCAACCATATTGTATTAGCAGAGAggtttttatattgaa
aacagtgttatcaataaaattcaaagaagaaaaattcaagtaaatagcGTGAAGAAGAGACGTTTATTGc
ggaatatattatatatagtaatgAATGTGAAGCACTTAACTTCCATACGACATGATGCATGGTAGCTCTC
AACTTGAGGACTTGCCCGGTCAGTGGTCAGTGGCTGAAAATTTTGTAGCGCAGCTTGTTAATTTTCAATC
GCATCGTTAAGTGTCCATTTTTTTAGAACAAACACCATGCGGACTAAGTGTGTGTTATCGCTTGATGCTT
CCTGGTATGTATATGCCAGATATATCCTTAAAGTTGTCGTATATGTTAAACCCCGATGAGGCATTACATG
GGGGTATTTCTTCTGTGTGTTATTTCAGTTCAAGTTGCAATTATTGTACCGCAACTCTCGAACTGTGTTC
ATACAGAACATTTTTTCCAAGTCTGTGGAACGTGttagatatattttttggtgCTACAACCGTTCGTGAT
TGGGTAAAATCGACACACGTTTTTACGTGTGAAATTATCTGTTATATAACtacaaataacaacaatataa
tattataatattatattgttgtaaaatattgttgCCATGAGCcgatatacaatatatttggtatttataa
ttaaacGAAAAGGAATTTTGCAAGTGAcctaaaaagaaaataattgaccaTAATAAATTAGACAAAGAGA
TGTTACTTAGGCAGTGGACGGGTTTCGTCAATTGTCGACTTGCCCGGGAACGCGATTCGAAGTGTCCTGC
AGCGCTGCACTTGAGAGACAAGTTTACTATATAATCTAGGATACACTAACTGTCGTTCATGTTATTCATG
TCTTAATGTCTTATAATAACACCTTTAGTCAGAGctattatagatatattattatattatttaagagCTA
TTACCACaagaatcaattaaatatttctatcttTGAATGGGTTTCAAAAACAGCCTAATCATAGctttaa
acattatttctacGATAACTGGAAATGAATGTGTAAACAAAGAATTTGCAAGTTAAAACATTTGACAATC
ATGACGAGACGGGACAATCAAGATCATGTGCTATTTGTAGATATAAAATAAGAGATCGATAACCTATCCT
CGATCGATTTCTATCTTGTTTCCAATATAAACGCCATGACAGAATATTTGTTGTTAGATTTTAGAagcaa
atttcaaatcattactTCGATTTACAGCTGAAGTTTATGTCTAGAATTAAGCTATAGTATATATCATAGT
AATTGTAATGTCTATAGTTCGAATAATCTTAACCATGTACGAGTAAAGATtttattagttgaaaaataaa
cttcaaagttattttcatgGCATTTCGAAGGTGATCAAAATGTCTACCTTCAAACGCATGGCTTTAGTCT
AAGCAGAGTTCAAATTCATCCCATCTTCAGAAGGGACTGGACCTATTCAACTGTTGAAGTTATGCGATCT
ATCAGATCATCCAATATTGTTGGTACTGTGAAAGCACATCAACGAAAACGTTGATAAATCACAAGAAAAA
GGTTACCTGCAGTACATTCTGGTGATCGCTGATGACAGAAACGTAtagatgcataattttttcggtACGA
AAGATCTACCGATCACGTATtcatagattaaaaatatttttacagaccGCCCGAACTACAATTAAGGTTA
CACGATTCCTTAGCGAAGAACGTATAAATTTTGAGAGTTTCTTCTTCCCAAAAGTACGTTTTTGATGCAT
TCAGCTCTTATAACATTACAGTAATAGTTTTATGATCAGattgtaatagtaatagttTTATGAAGTCAGA
TTATATTATCAGCTTATACCTGTTACGTATAATAAATTACGATCAtaacttcatatttattcaatacaat
gGCTTAGCCGTATGAAACAAATGGTCTCAACGCTCGAATGGAGTGCATATCGTAACTCTCTTCATAGAAT
TTCCAATAtgcaaatataattcattactCTGATTGTATACCATCCAATGAATAGgtaaaatataacatta
ttttctaaAGCACTTGGATATCGGGGACAACATAAACACCGAGGATACTTGTTGCTTAAGTATGGCCATA
AAGATACTTTATATCGTTTGAATCTgctaaaatttagaattttgggCCTCTCTTCATCTGAATCGTGaac
aattttgtataaaatgagTGCCAGTTTTCTTTCATATAGTCAATAACAATTAGTGACATTGAATAGCTAA
GGCGAGAAATTATAATTGCAACCAACAAGGCAATTGCATGTTATTTTCAGTGCAAAGTCGGATCGCTTGA
ACATGTTTCGTCCCATATCTGTGCTAGTATAATACTCGTATCAAGAGGGGTGTTCTCCCTGTGgaggtaa
aaaaaacttagttAACAAACGATAATCTGTTGCATACAATACAGAGCATGGATTACTCATCAGGTAATCA
ATTAAGACTTGCTTGCATTCCActttatttccaaaaaataatatcgtctttatttcatcaaatcGGAATA
TATGAAGATATAATTTCTATCTATTTAATGAGACGACATTGCCATTTTGTACAGTCCAGGTGCATAAAGT
TAAGAAGCTATCATTTTTGTATTatctaaaactaaattttggtctaaaataagtgaaataattgGCTTTT
AAATAAGGTTTAAACACGAATATTTTAGAACAGTGTGAATATATGCCTACGTACCGTACCGTATTTTCCT
TCCTTTAACGGGTTCTCAAATGGACGattcttaaaatgaaattttttgtgttatgCTATTTTTAGCCATT
TATATTGAATAGTAATGATTGTTTTACTAACTAATGACATTCAATAAACTACGTGTTTATCAACTTTACA
TGTTCACTGTCAGCCGCTCGCTCCACCCTTGCTAAGATTTAGATTTATATTGTACTTATTAAGCGTATAT
CTTGCTATCAGTAAATTTGCATGTTGCGGTAGATCATCATCTGTTGTACAATTATATTCGTTTCAATATG
TCTTGCTATATATCAATAACCTTGCATGTTGCGATAGAAATAGATCATTATTTGTTGGATAATTATATTC
GTTTCAATATATCTTGCTATCAATAATCTTGCAGGTTGCGATAGATCATTAtcttttgtataattatatt
cgTTACTATAAACATAATACATGCAATCAGCAAATTCTTCTGAACTAACAGCGATggctttttaacacgt
tttcttagcttagcttcgcgtttgtttgtttgtttgtttgtttgttgttaacattgacacgacacacgtt
ttttggggcgagccacaatttattttctactttttagtccacaagaaaaacgtgtttttttaaaaagttt
ttttaagtattttttcattttgcttatgAGTCACTGACAGCCTTTGACTTAAATTTAAACCGCCGTCTGG
ACTAAAGTAATAccatttaaagtttttctttgtatttctATCCacagataatttaatattcatttaaata
ctcTGAATATctcacatatgaatttttcctttGCAACCAttgacataatttaaaataatataaagacaTA
AAGAACTTActccaattttaatttaaatgttgccCAAATGTGTGTTAGAATAATGAATATCCCGCTTCAA
TAAAGCTTCT

Peptide
DNA photolyase
FAD binding
MSERNELPPMATSSEEPQAPPTTQAAVSRGPRQRQDSLDVVLSRSMVAERVMPARFSSNYHAMQSMPMVANKILTGTMIMTDPNAMTHMTMFGDYKLPGKSLIPPSMMQVRSMYADIKSIQGRGSLDDQMYTMQPMVTDGLVLPGGMLTHPLMTSKSSMVKDDMMTGITMISGKPLSGRVMVGNARAVTTRAATTDKTSPTKTPNCDRVLQNRPPEDRTIPCPKIIGPSVASKPHGMGDKHPLSKLRQIDRMTASRYASEQSQGPSSDKKIPDNHKTQFKSFPENSSRNLKEEFVVNTNDSLGKNIQSAERESEFVFGRNPKFGSEDFGTSTVDSMALDRTKSPGQNSSFGLKMSPNKTVSPVMRSTHRTILGDNNLAQPECSGSSGPSMSSEKNSSTIKVRSGKHIVHWFRRGLRLHDNPALRDSIFNCETFRCIYILDPWFAGSSNVGVNKWRFLLQCLEDVDNSLRNLNSRLFVVRGQPANVLPQLFKEWNTTVLSFEEDPEPFGRARDASIIGIAREMGIEVVVRTSHTLYKLDEIIEKKGGKPPLTYKTFQNILAMMDPPPAPVPPIVAGDLQHAFTPIQPDHDDKYGVPNLEHLGFETEQLPPTAWKGGETEALQRLKHHLERKAWVASFGRPKMTPQSLFACPTGLSPYLRFGCLSARKFYTELNELYTKIKKVPAPVSLHGHLLWREFFYTAATNNPKFDHMKGNPICVQIPWDKNPEALAKWAHGQTGFPWIDAIMTQLRAEGWIHNVARHAVACFLTRGDLWVSWEEGMKVFDELLLDADWSVNAGSWMWLSCSSFFQQFFHCYCPVRYGRKADPNGDFIRTYLPVLKNFPTKYIHEPWMAPESVQSSARCIIGQHYPLPMVDHATQSQTNIERMKQVYRQLAHYRATMSSRSGGDSKSRCKHQQTSAPNRSLCSNRVLTTV

T. saltator – constructed from NCBI accessions GDUJ01076706 and JQ413343
Peptide
DNA photolyase
FAD binding
MEERNVTESSVVATSSQDIQGPSGTKSSVVLQSRSTPDDLNVVLSRALVAERVVPSRFSSNYQTMSSMPIMTNKILTGTMIMTDPNAMTHMTMFGDYKLESKSLIPPSMMHVRPMYGDIKSIQGRAVLGDQICTVQPMVTEGLVSSGRIMTPQLMAPKTSLIKDDMMTGITMISGKPLSGRVMVGDARAVTTRVVTSDKTSPAKTANSDRISHSRPPEERTTPGSKAVGSPAIPKSNILNDKHPPSKLSQIDRMTASRITGEPPLTPPVKVEKKNHATHKSSSKNLLHEKACKIQITEKHNFQKNDSLDKNNLSEHVSSTHSEFMFGENLSFGCENNGTSKLNSMTLDRTKSPGQSSSVDVKMSPNKIASPGERSIHRTILGDNNLTFSESGSSSGLAFTAEKKNTSKVVSGKHVVHWFRRGLRLHDNPALRDAIVNCETFRCIYILDPWFAGSSNVGVNKWRFLLQCLEDVDNSLRNLNSRLFVVRGQPANALPQLFKEWNTTVLSFEEDPEPFGRARDASIIGIAQEMGIEVIVRTSHTLYELDKIIKKKGGKPPLTYKTFQNILAMMDPPPPPVAPIEASDLKHAYTPLQHDHDDKYGVPNLEHLGFETEHLPPAVWKGGETEALSRLKHHLERKAWVASFGRPKMTPQSLFACPTGLSPYLRFGCLSARKFYTELNVLYTKIKKVPAPVSLHGHLLWREFFYTAATNNPKFDHMKGNPICVQIPWDKNPEALAKWAHGQTGFPWIDAIMTQLRTEGWIHNVARHAVACFLTRGNLWVSWEEGMKVFDELLLDADWSVNAGSWMWLSCSSFFQQFFHCYCPVRYGRKADPNGDFIRTYLPVLKNFPTKYIHEPWMAPESVQRNARCIIGQHYPLPMVDHGTQSQNNIERMKQVYQQLAHYRANISTRPCGDSKLRCKYPLHSTA
```

***period/*PER**

***P. hawaiensis*** – identified by alignment and expanded using mapped reads.

Aligned sequences

LQNS02276498.1:4910093-4910393 ATGGAGTTACTGCCGCCTCCTCGAGAGAGCCTAGGGCCTCAGGGACCACCCAATGAAGGTCCTATTGACCCCGTGGGAGACCCGAGATCATCTCCCCACCCGAAGGTAGACTCCACCACCGCAGACACCCGTGAGCAGGAGCGTGGCCCccagccacagcagcagcaacacgatGCCAATCAGGACTCCACTAACCCCAGCAACCAAACCGACCAGGGTTATGGGTCCACTGAGTCGTCTTTCAATGGCCAGAGTCACAAGAGGTATGAGCTACATGGTGTAGAAAGCGTTACCAAGTGT

LQNS02276498.1:4911572-4911839

CTCACCAAGAGCCGCAACAGCGGCTCCAGCCAGAGCAGTGGGTTCGGTGAGCAGCACAAGAAGGCTGCCCACACGGTCAGCTTGTCGTCCACAGCGCAGGTTACCACAGCCAGGGTGTCGCCCATAGCTGAACCCCATGGCCCCGGCAACAACACTGATAACCTCGCTGACAACCTTGGGGCGATCTCCTGCAAAGCCAAACCTTCTTCTACTTGTACTGTCGAGAAACTGCTCGACAATGCCCACTATGCCTCCACTGTACCAAGG

LQNS02276498.1:4914493-4914871

AGGGAGAAAAGATCCAAGGAGCAGAAGTTGCGGGACGTCAAAAGGAAGAAAGTTGAGGATGGCCACTACCAGTCTTATGACGGCTTCCTTGAGCCGTCAGAGGCGTACACTTTCGGCCCTGGACCACCGAGGCCGTCCGAAATATCGGACGCCAGGGTCGACGACTTTGACGCTCCTATGCTGTCGCTACAAGCTCAAAAGGATGAATTCCGACCACCGTCAGGAAACACAGTGCATCTGATGCCCCAATGCTCTAAGACGGACCACGTCATCACCTCTAGTGCCATGTGCCTGCAGCCCATGGCACTGCCATCAGCGGTCATTGCTAGCAGGTCTTCCGAGTCGTCGGTGGTACCGGCTGTTCAGCAGAGCTTCTTC

LQNS02276498.1:4916999-4917110

ACTTCGTATGCGGTAGCGAGCACCGGTGCCGTTAAGCGTGAGCCACCGGATCACCCCACACTCATCTACACTCAAGCCCTTAACTATATCCTGCGCATCAAGGAAAGCTTC

LQNS02276498.1:4919611-4919815

CAACGAGGATTCACCATGGTACTGAGCATTAGGGATGGCACCGTCATCAGGGTATCGACCAACATGTCACAAATTCTTGGATTCCCTGAAGACAAAATTGTGGGGCACTCCTTTATCGACTTCGTCTACCCCAGAGACTCAGTCCACTTTTCTTCCAAAATTGTTAGTGGAATGAGCCTTCTCATGAAAAACCCGAGTACCAAA

LQNS02276498.1:4920508-4920742

GCAGAGACACTGATGTCACCTTTCTACTGCCGAGTGCGGGAAAATAAATGCTACCAGGCCTCTACTTTCGACATGAAGACCAGAGACTCGTACAAGCCCTTTAAAATTACCCTTAAGTTCAATGAGTCTCTTCCGTCTCTGGAGCACGAACTTATGGCTCCAATATCGCCTGTACACTTTCCTGCACCTGACGTGTTGCTTGCCGAAGTCATACCTGTACCATCCTTTTACCAA

LQNS02276498.1:4921827-4922007

GTTCCTGATGAGGTCATCAGCGGCGGGAATTTCATCATCCGCCACTCGGCTTCTTGCAACTTCTCCGAGTACGACCCAGACGCCATCCCATTCCTTGGCCACCTGCCGCAGGACCTCACCGGTAACTCGATCTTCGACTGCTACCATCCTGAAGATCTGCCACTGCTACTTACCATCTAC

LQNS02276498.1:4923328-4923484

GTAATTCGAGAGGAAGGGAAGCCCTTCAGAAGTGAGTCGTACCGATTCCGAACCTTCAACGGCAGCTGGGTTGTACTGGAGACAGAATGGCTCTGCTTCGTCAACCCTTGGACTCGTAAAATTGACTCTATCATTGGTCATCATAAAGTCATCAAG

LQNS02276498.1:4923570-4923780

AAGTTTAGGTGCAGACATAGCTTGAACGAAGGTCACCATTGCAAATTTTCCCAGTTCTTTACATTATTTCATGTTTGTCTTTATGAACAGGGACCGCGTGACATCTCGATCTACATGGAGAAGTGCGATGGACCTTTGACCAGCTTCTCAGAAGAAGTTTGTCTCATTGCACGAAAAGCTCAGAGGGAGATCATCGACCTGCTCTCAAGA

LQNS02276498.1:4925098-4925158

CCTGTGGCAGCTGGTCTAGCTGAGGCTCTGCGGCAGGAGACGCGATACCCTACTGTTTCG

LQNS02276498.1:4926501-4926582

GCCGATGCAGCAGGACCAGAGCTCCTCCAACATGAAGATTCCCGTAGCTCGTCGGAAGCGCACCCTTGTGGAGATGATGGG

LQNS02276498.1:4933506-4933782

AGGAAATATGGCAGCAAGGACGTTACGGACAGCGGCAGTGGGGAGAGCGCCAGTGGACCCAGCTCCACCAGCGGGGGTGCGGGGTCTGCCATGTACCGGCACGTCCCCCTCACTGAGGAGGTCCTGTGCCGCCACAACCAAGAGATGCAAATGCTCTTTATGGAGCGACAGAAGAAGAATACATTAGCCATTACAACTCCTTCCCACCAGCGGGAGCGACTCTCTagacagaaaatcaaacaaatcaagaAACCTGCCAGGAAACCATCGTCTAAA

LQNS02276498.1:4936246-4936420

CAGTACGCAATGAAGAGACCAAACTCTGGGAGCCAGAGGGGTGACGATCGGGCCCACAAGTTTCCCTTCATCGAGAAGAATTCAGGGAAGGTATCGGATGGAGCGGCGCGCCCCTCGCATGTGAACGGCTCCACGTCTCAGCGCAACCATATGGCCAAGCCCACCATTGACTCT

LQNS02276498.1:4937554-4937809

AAGGTGGCAGGCCGCCCCAACCTGACGGTGCCAGTCAACCAGCCTGGATGTTCATCAGATCGTCCTCAGCCTACGTTCTTCTTCCCCGGCAATGCCATACCGAACCCCCCCAATGGCTTCCATACCAACTCTTTACCCTCCAACCCGTCCACCAGTCTCTCTCAGGAACCCCAAATCAATGCAAATGCCCAACCCTATCCAATGACCCCAATGATGGTCAATCCCCATCAGTCCCACTACATAACTCCGCAAGGT

LQNS02276498.1:4944064-4944292

ccccaCCAACAACCTCAACAACACCATTTACACCCCAAATCTCCCCATGGGATTGTCCACTTGAGGAAGAGGTCGGACTCTCGGGCCACCTCCGTCAAGGTAGAGCCTGGTTCGGTCCGGGGCAGCGTGGCGTCTGCTTCGGGACAGCTTAGGTGCTCCATGTCCCACCAGCCCGAGAGCCTCAGGTCGGAGCTCGACGATGCTGCAGCTGCTGAACAAGGGGTTAGT

LQNS02276498.1:4945420-4945555

CTGGTGTACGTACCGAGGGTTGTGTCCCACGCGTCCCACTTCTCCCGTTCCACGAGTGTGTTGGGGGAGGCTGAGAGCATCGCGTCGCCAGAGAAGAGGCAGCCGTGTATGGAAGACCAACATGCCGAGAATAGC

LQNS02276498.1:4946568-4946598

GATATGGTGATCTCGGAAAGTTCGCCCGCT

LQNS02276498.1:4948381-4948558

GCACAGAGCGTGAGCGCGCTGCTGCCATCCCTCGACAGCAGGCCTGTGCTGCACGACCCCTCCTGGTTGGATCACGTAGAGGTCACACCGCAGCTGCTTTACAGATATCAACTGCGCACCAAAGAGATCGTTGATGTTCTAAAGAACGATATGGATGCCCTGAGGGAGCTAAGTCAG

LQNS02276498.1:4951013-4951163

CAGCCTGCCCTTGTGGAGGACCAGTTGTCTTCGCTCTACCAGGAGCTGGAGATAGACGGCGAGCAGTTACAGCTGGACGAGGGCATAACTTCCTCGTCTGGGGAAGAAATGGTCGACGCTTCCACGAAGGTAACATGGCACGCCAGCAGT

LQNS02276498.1:4953323-4953493

GCGTCCAGTGATAACCGAAGAATGGAGAAAATTCGTTCAACAAGATATTTCAACAAACAAGCCATTATCCACGAAGTGGAAGCTGCCATACCTCCGCCTGAACTGCGTGTCAACCATCGCTATTCAGTTGCATCTGCGAGGCCGGTCTCTATACGGGAGATCCAATGATG

Constructed peptide

PAS

PAC

PERIOD C

MELLPPPRESLGPQGPPNEGPIDPVGDPRSSPHPKVDSTTADTREQERGPQPQQQQHDANQDSTNPSNQTDQGYGSTESSFNGQSHKRYELHGVESVTKCLTKSRNSGSSQSSGFGEQHKKAAHTVSLSSTAQVTTARVSPIAEPHGPGNNTDNLADNLGAISCKAKPSSTCTVEKLLDNAHYASTVPRREKRSKEQKLRDVKRKKVEDGHYQSYDGFLEPSEAYTFGPGPPRPSEISDARVDDFDAPMLSLQAQKDEFRPPSGNTVHLMPQCSKTDHVITSSAMCLQPMALPSAVIASRSSESSVVPAVQQSFFTSYAVASTGAVKREPPDHPTLIYTQALNYILRIKESFQRGFTMVLSIRDGTVIRVSTNMSQILGFPEDKIVGHSFIDFVYPRDSVHFSSKIVSGMSLLMKNPSTKAETLMSPFYCRVRENKCYQASTFDMKTRDSYKPFKITLKFNESLPSLEHELMAPISPVHFPAPDVLLAEVIPVPSFYQVPDEVISGGNFIIRHSASCNFSEYDPDAIPFLGHLPQDLTGNSIFDCYHPEDLPLLLTIYVIREEGKPFRSESYRFRTFNGSWVVLETEWLCFVNPWTRKIDSIIGHHKVIKKFRCRHSLNEGHHCKFSQFFTLFHVCLYEQGPRDISIYMEKCDGPLTSFSEEVCLIARKAQREIIDLLSRPVAAGLAEALRQETRYPTVSADAAGPELLQHEDSRSSSEAHPCGDDGRKYGSKDVTDSGSGESASGPSSTSGGAGSAMYRHVPLTEEVLCRHNQEMQMLFMERQKKNTLAITTPSHQRERLSRQKIKQIKKPARKPSSKQYAMKRPNSGSQRGDDRAHKFPFIEKNSGKVSDGAARPSHVNGSTSQRNHMAKPTIDSKVAGRPNLTVPVNQPGCSSDRPQPTFFFPGNAIPNPPNGFHTNSLPSNPSTSLSQEPQINANAQPYPMTPMMVNPHQSHYITPQGPHQQPQQHHLHPKSPHGIVHLRKRSDSRATSVKVEPGSVRGSVASASGQLRCSMSHQPESLRSELDDAAAAEQGVSLVYVPRVVSHASHFSRSTSVLGEAESIASPEKRQPCMEDQHAENSDMVISESSPAAQSVSALLPSLDSRPVLHDPSWLDHVEVTPQLLYRYQLRTKEIVDVLKNDMDALRELSQQPALVEDQLSSLYQELEIDGEQLQLDEGITSSSGEEMVDASTKVTWHASSASSDNRRMEKIRSTRYFNKQAIIHEVEAAIPPPELRVNHRYSVASARPVSIREIQ

**Section B – Regulatory genes identified in head transcriptome**

Casein kinase IIα

>lcl|PH.k29.comp2312_seq0

ACAAGCCACAATAAATAGTAGAGCCAAGAGGCGGGTCTTCTGTGTACTCCCTTTTGCCTCTTCGCCATTTTGCGCTTTCA

TGTGCACACTGGTGCTACTCATTGTTAACGGATAGTAATTTTCTTCTAGTAATTACAAAGTATATTTTTGAAGAGCGAAC

AGTCGAAGAATTTCTTTTCACCGTAATGTTTTAGATCATTCTGGCAGCAATAGTGTTGTCTGCACTTCGGAATCTCCCTG

TGCCTGTAGCACTGCTACGGTCGGCTGCTGCTGCTGCTGCTACCCCCGTACTTCCCTTGGCCACTGGTCCTGTCATTGGC

AAGGTCAGCCTTGCTGCAGTTGTTTACGTGAGCACGGCAGCCTCTACAACTGTCACCACGACGACCCACGTGGCTAACCC

GGCATCCTCTGCCACCACCATGATGCCTTTTCGCAGTCGTGCGAGGGTCTACGCGGAGGTCAACACTCTCAGGCCCCAGG

ACTACTGGGACTATGAGTCTCATCTCATTGAGTGGGGCCAACAAGATGACTACCAACTTGTACGTAAGTTGGGCCGTGGA

AAATACTCGGAAGTATTCGAAGCAATCAACATCAATAACAATGAAAAGTGCGTGGTTAAAATATTAAAGCCCGTCAAGAA

AAAGAAGATCAAACGTGAGATTAAAATATTGGAGAATCTCCGGGGCGGGACGAATATCATCACCCTGCAGGCTGTAGTGA

AGGATCCCGTGTCCCGCACCCCTGCGCTCGTCTTTGAGCACGTCAACAACACAGACTTCAAGCAGCTTTATCAGACACTT

AATGATTATGATATTAGATATTATCTCTATGAACTGCTCAAGGCTCTAGATTATTGCCATAGTATGGGCATCATGCACAG

GGACGTGAAGCCCCACAATGTAATGATCGACCACGAGAATAAGAGACTGCGCCTCATTGATTGGGGCCTCGCCGAGTTCT

ATCATCCTGGCCAGGAGTACAATGTTAGAGTAGCGTCCAGATATTTCAAGGGTCCTGAACTTCTCATTGACTATCAGATG

TATGACTACTCCCTTGATATGTGGTCATTAGGCTGCATGCTGGCAAGCATGATATTCAGGAAGGAGCCTTTCTTCCACGG

GCACGACAATTATGACCAGCTTGTCCGCATTGCCAAGGTTCTTGGTACAGAAGAACTTTTCGAGTATGTAGAGAAGTACC

AAGTTGAACTGGATCCCCGCTTCAATGACATTCTTGGCCGGCATTCCCGTAAGAGGTGGGAGCGCTTCGTCCACCACGAG

AACCAGCACCTCGTGTCGCCAGAAGCTTTAGATTTTCTAGATAAGCTCCTTCGCTACGACCACCAAGAACGGCTCACAGC

CCGTGAGGCAATGGAACATCCATACTTCGGGCCGATTGTTAAGGACCAGGGCATCATGATTGGGTCACCCACGCCGCAGC

AGGCACCACCTCCAGGCATTCCTGGAATCCAGGAGTAGTCCAGTGAAACCGTAGGTGATCGTGATAGCCCTGCCTTGTCC

TTCAGTTCCCCGTATTGGCCTCTCCCTGACTGCGAGCTGTGACGCCTTTTGGTGCCAGACTTCATTTTTTTTACCTCGAA

ACTTATGCTCTGCATTATTGCTTTGCATTGTCTTGTAATGTAACGCTGTGAGGAGGTTCTGCCGTGCCGTAATTATCTTT

ATATTAATACTGTACCAGCGGTGCTGCTAAACGTGGCATCAACCTATTGCTTTGAAAGTAATGGTTCCAAATGGTAGCTA

ACACCCAGTGTTGTCCTTCGTCGGTGTCGTCTTGGCTAGCCGTCGCGCTAATTTTTTTTTTTTTTTTTTTAAACATTGGT

AAAACTTTGGGGTGTGTGCGAAAAGTGCTAATTTATAGAAATTAATATATTTTTTAGATTTCAGACGTGGTGTTACCCTT

TTTTTCCTTTTTTTGCGATTTGTACCTTTTTTGTGAATTTCATACATTCGCAACGTCCTAGTTGAACGAGTCTGCCTGGC

TGCTGCATGCTCCACTACCCATACTTACGACATTGCTAATGTTATTACAAAATTCTATTGCTTCGCTGCCCTTACTCGCA

TTGAGTCGTTTCATGTTCTTTAATTCAGAGTATTTCCGCTGCAAAGCAATCACAAGTTAACAGTGTCATATCTCCAGCTC

TTTTCATTATATTTATGTCAACACGTCGACGCTACATTCTCTGGCCAACACGCTCAAGTGAATTGGCCTCATGCGGTATC

ATCGTTACTAGTAGAGTAAGCTTGTGGCAGTTTCCTTTGCTACAGACAGTGCAAATACTGTAGTCGTTGCAATCTGACCG

TTATTTTTTTTTACTCAACTTAAAATTCCTTTGTCTAGGTTGTAACTAGGTAACAAGAAATTATGAACTTTTCCGTTTGT

ATAGGCAATTTCTGCGTAACGTCAAGCTAGTCACGCACTTTTTTTCACCTATAGCTGTGTGTTGCCATTTTTGTTATTTT

TTTTCCACAGTTTTCAAGTGCAGCTTGGTTGCTCCGGTTGAGTTATCTACGGTTGTCACGTCTATACACGTTTTGCCATA

GTTCTAAAGGTTTTCACTGCAGAAACTCTTCCTGATTTTTGCAGAAAACTCCCTCTTCAATTGCACACATCTGCAAATTT

GCCATTCATTTTGCAACGTAGGGAAATGAATTGTTTCATGAACTAACTCTACCGTATGCCCTTAATGAAGTAATTAACTA

AGCGAAAACTTCTGTGCACCACATACTGTTACGTGTACAACATTGGGGTAACTCGACTGTCTTTTTTGTTTATCTGCAGC

GAGATCAGTACATTATTGCGACAGTAATCCTCGGCACGGATTTTCATTAGTTGTGTGTGCAAACTGTTTGAACTAGACTA

GGACTGTACGTGCTCAGTGTTTTGTATTATTTGTATGGAATTTATGTTCATGCTTACTTTTCGCGTTTGTATAGGTGAAA

CGAGCACTGAGTGGTAATAGTTTGTGTCCTCATCGACTTCAGCTGACAAGTGGGGTATGCCTGGAACCATTCTCTGCTTG

AGTCTACTTAATAGTCCTCTTCGCCCCTTTTTCTATCAGTTTGTCTAATAATATGTTGACCGCAACTGCCAGTATAGGTA

GTAGCTACTACATATTTGTATGACTGGGTGCCATGTATGGGTGGCACGCACCGCGACCAGCGTAAGGTGTTTTCTTTTTT

TTTTTTCAACTTCTCAGCCCTGCTAAGATCGTGTTTGAAATAGTATTTTTATCAGTGCAGCTCACCCTACAGTTGTGTAC

TTACTACGGAGAATATCCATTGTTTCGCTATGTTCACATTATGTAACCGTTGTCCTATACGTACGTAGTCTAGAGTGACT

GTGGGCGTTTGCTTAGCTGTCCCCCGACCCTCACACGCCGACTGTAGCCTGTAGCTGCATTAAAATCTGAGCTTTATTGC

TGCTAAATTAATGGTCGCTATTTTTATCTACTGCATTTAAGGCTTTTTTAAAGCTACACGGCGGTATAAGCTTCGACGGG

AAAAACATATAAAATTAGTCGGAATTTAATGTCGCCATCGGGCTACACTTGGCGTGGCGGGATTAATGCCTTGGGTGCTC

AGTTAAAGAAGATTTTTTTTTTTTCTACGAAGAGCTAGAGTCTAGCTTTGATTTGAACGGGTTTACGGACGTTTGCTTTG

CTCTTGTAGGCTATTGACTGTCTCGGCTGTGGTATAGGTAGCGAGCAGTAGGGCGCTGCAGCATTGGTGTTCAGGATACA

TTTCAGTTAATTTATCAGTTATAGAATAGCTTTTAAAAGTTGACTATTATCTAGGTGTCCGTAGTAGAGTTTTTTCTATA

TATAATAGAGAGAGAGAGGCGTAATTCTGTACCTTCACCAAAGTTATTTCGTATGTTGCTTGGGACACCGAGGGAGCTCT

GCCTCTCGCTTTCGACGAAGAAGATTGGCTGTGCCTAGTCAACGTGCAGATAAAAAATTATTATTTTTTTACACATACTC

GTTTTTGTTGCTAGAGATCATTATATGCGCTGCATCTCGCCCGATCGGAGAAGCAGGCTATATTGGGCCTGGCGTGATTA

TTTCGTTCGTGAGGTATATTCGGCGCAATGTTAGCCTTGTAACAGAACGCACGTGTTTCTGGTGAAGTGAAGCATCAGCG

GTCTAAATTATTTCTCTTCTACCTATGTGTGCATTTTACTTGAGAGCTTCTAGTACCTGGTTGCTTCCGCATTTTGTTAA

AGCCATGCCCGACGTTGCTTGGTAGCTAAATGTCTGCTGCTAGACTGATTGTAGTAGAGGTGATGCACGCATACTTTTAT

TCATTCAGGTCTTGCAGTGTATTTTTGTTTAGAAACGAGAAATCGCTACGTAGCTACTTCACCCAGGCGTAAGTGCTTTT

TTTTCTTCAAAGCCATGTTTCATATATTCCCCGATTGTTCTCTCGCGGCTGCGGATTGTGCCGCCGTCAAACTCGTGCTG

CTAGCATAGGAATTATTCCAAGTTCAGGCATATTTGGATAAAGTTATATAACGCCTTCGCGAATGTTCTTTGCCAGAGTG

ACTTGGCGTATTGTTGTAAGTTTAGTACGTGATGAGGACGTGTTGTAAGAGATATTTTAGTGATAAGGAATGTTGACGGC

AATGAGCGAACGTAACTCTTGTAGCTAATACAAAATAAATGAGCAGAAATAGTTGATTATACTGATAGTCTTTGACTATA

TAAAAGATTGATGCTGTGTGGAAATAATGTGCATTGTATTGTAAATCAGTTAACGATGTTCTCGTTAGATTATGCCCAGC

TCATGGTAATAAACGTTCTATTGATCATTCGTATTACGATGCATTTTTAATTAATTACATCCTGTATCGAACAAAAAAAA

AAAAA

Casein kinase II β

>lcl|PH.k21.comp3428_seq3

TTTATTGATATCCTTATAAAAGTCTTTCGTTTTGATACAATAACATAATGAATCTGTATCCGTAAATAACAGTCTTGCTT

TTTCTTTGTATCTGTTTTTAATGAAATTGTAATGAAAATCATACATTAACGTTTTACTGATATCTAGTTGTTAGCTTGGA

CTGTACAAAATGAGCAGCTCAGAGGAAGTGTCATGGATTGCTTGGTTTTGTGGCCTCAGAGGAAATGAGTTCTTCTGTGA

GGTTGATGAAGATTATATCCAAGACAAATTCAACTTGACGGGATTGAATGAACAGGTTCCTCATTACAGACAAGCTCTTG

ACATGATTCTCGATTTAGAGCCAGATGATGAAGAAGACTTGCCCCACCAGAGTGACCTTGTGGAACAAGCAGCAGAAATG

CTTTATGGGCTTATTCACGCAAGGTACATTCTTACCAACAGAGGCATTGCACAAATGATCGAAAAATATCAAGCCGGGGA

CTTTGGCCACTGCCCCAGAGTTTATTGTGAAAATCAGCCTATGCTGCCAATCGGCCTGAGTGACGTGCCCGGGGAAGCGA

TGGTGAAGCTCTACTGCCCAAATTGCTGCGACGTATACAACCCCAAATCGTCTCGCTACAATCACATCGATGGCTCATAT

TATGGTACCGGCTTCCCCCACATGCTCTTTATGGTCCATCCAGAATACCGTCCCAAGAGGTCCACCACCCAATTTGTTCC

TAGACTTTATGGCTTCAAGATTCACCCAATGGCGTACCAACTACAGCAGCAGGCAGCTGCCAATTTTAACGTACCAACAG

CTATACGCCCAGTCAATTACAACAACGGTAAAAGGTAGTGAGCTTTTGGCTACCGAGAGGCGATGACTCAACATCCCATG

GCTGTTATCTTTCAACTTGAGAATCTTTTAATGCCTAATGTCGAGCATCTTTTGGTGCATGTAAGGCTTATGCTTGCAAT

TTTTGGTATATCCATGTTTTGATGTTATAGCTGTCAAGTTCCTTCCCGGCGCCCATCATTATGAAAGGTGCTAATGTTCT

TCTAAATATTGGTGTTTTTCTTCCTATTCAATTTCAACATTGCTGCAGTAAGTAACTTTTTCTCGACAACAACAGCTCCG

ATTTCGTTTGTTTTGCATTGTTATATAAAAGGACTGATGAACAATATTGATTTTGTTCTTTATCTATTCACTTATGAAAT

TACTGTTTTTTGTGAATATTCACCTGTATTTTCTTTATTCTATATAACTTATCGACTTGTAAACAGAAATTTTTAGTCAC

TGCTGTTAATGAAAGGGTTATTGCATTAGGAACCTGCACGTATTTAGGAATAAATTCTTTAGTGTATCGTACTGTAATTT

TATTTTCTATTTTGTAAATAGATACTACCTTTTAAACTGATTTAGCATATTTATAAGATGAGTGTTCTGACCGGACTTTT

CTTCAACCTTTGTTACGACTACGTAGTATACCTCGTGTTGCACCAGGACCCTATAGGTTGCCTTAACCTTGGGTCTTCGG

GACGTTTTTTCTCTTCGGGCCCACTTTCATACTCCTTCTGTTGATGCTTGTTGTCATCTTCTAGACTTGCAGTTCCCATA

AACGTTCATCCTGTGTGGGAGCACCCTTGTGCATTCTATCCGCGTTTCTGTCGACGTATTGTATGTCCCTTATTTCTCCC

TCTTTTTCATCGCCATTGTTGGGAAAAGAGATGCCTAAATTTTGCAGTGTCATAATGAAAAGTATTTGCTGCTTTGAGTT

TGCACACTTCTATTTGTGATGTTTCCTGCTCTTCCTCTCCTTGCTCGCTATGCTGTTTCCATACTGTTGGTAGCTACAGT

GCAGTGTACTTCTCATTGTTCGGTGCACTATTTTGTATCTGGTCTGCTGGGCTCCCTGTTATACGGTGGATAAGTGACAT

GCACCCTCTGTGTAATAATAACAACTAAAGCACTCTGAGAGCGCTTACCTCCGCCAAGCAAGGCATCGGCAACGAAAAAT

TCAATTGGTTCCGAAACCATGATTTTCTTAGTATCAACACCAAAATCTAATCGTTTGTGCCCTTCAACAGTGTTAACCCA

TTCAGTAATTTTTTTGGAGATCCGTTCATGACTTATTAAGTTAGGGCACTAACAAAATATAGGCCGACAGACGAATGGGC

GGATCTTCACCAAAGTTAGTCATCTGTTCCTCCTGACATACCTAACAATTCCTGTCAATTTCATCCAAATCGGTTCATAA

CTTTTTGAGTTATCCTGGTGACAGACTAACAGACAAACAGACAGTCAGACAGACAGACAGACAGACAGACAGACAAACAC

GACGAAAACATAACCTCCTTGGCGGAGGTAATTACCC

Clockwork orange

>lcl|PH.k21.comp80764_seq0

CTAGCTTACTCCGCGCTTCCTATCATCATGTTCGTACAACTCCTTACTTCTGCTCTCCGACCCGCTGTCTCACCGTGTCA

TCGAGAAGCGCCGGAGGGACCGCATGAACAACTGCCTCGCCGACCTCAACAGGCTCATCCCTCCCTTCTACCTCAAGAAA

GGCCGGGGCAGGGTCGAGAAGACTGAAATAATTGAAATGGCTATCAAGTACTTAAATCATTTACAGCAGCAGCAGCACGA

TTTTCGGGAGATCTCACACAATTCTGTATCGGGATCAACCGCAATGTCAGGAGACCAAAAAGACCAGTGGGTGGCAGGCT

ACCAGGAGGCCATGGCCAAAACTCTGCAGTTCCTCGTAGAGGTCGAAGGACTTTTTTCTGGCGACTCTCTGTGTGTGCGG

CTCATGAACTACCTCAGCGTGCACTGCAAGACGACACTAGCTCAAGAGTGCTACTCCAGCCGCAAATCAGCAGCCAGTCC

TGCCTCGAGCAGCGGGTACCACGCCAATGGCTCTTCGAGCGACAACGGCAACTACAGCAACTGTGGGAGCAATGAAAGCA

CCAGCTGTTCGCCCGACAACCCGCCCTCTTATGGCCCAGCTGCTGAGGTGAACAAGTCCAACACCTGTGAAGGAGATGCG

GT

Circadian trip

>lcl|PH.k31.comp1615_seq0

GTCAGGGTAAAGTGAACACAGTCTGTCGAACCAACTTCATGTGCTGTGTGAAGTTAAAAGCTCTTCGTTACCTAGTCACC

ATTTCCGGAGAATTTTATGCGTCATCAGTAGTGTTGTTGGCATGCCTGTAGTGTGGTGTGGCAGTGCCTATGGCCGAGCT

GCGGAGCTGTGAGGCTGGCAGCACACCGCTAGGGGGCTCTCAGTTGCCTGCTCCAGCCTCATCCCCATCAGGTGGTAGTT

CTCACACCAATACCTCACGTAGCAACTCGTCCTCTTCACGCAGCAGTCTTGTTCGCACTCCCTCCCACTGCCAAGCAAAG

GACGGAGCTGCTACTCCCGTCAGCAGAGGCAGTCGCAGGAACAGTGTTAACAAGCTCCCTTCATCTTTCCCTTCCATTGC

TACTCGAGCTAGAAGCTCAAGTGGCTCTCATCAGACTTCTGAGACCGTAGGTGGTAAGACTGGTAAAGCTGCTGGTCACC

AGTCTCCTTCACTGCCGTCCTTTGGTGAGACAACCTCCACAGTGGCGAGTGTCTCATCCCTGCCCCTGCGCAAGCGCAAG

TTCTCAGCTGTTGAGTGTTGTGCCATTGATCTTAGTGTTAAAGGCTGCAACAAAGATAGTAGCTCTAGCAGACCTCAATC

ATATCTCACTTCTGCCCATTCTTCTAAAGAGGAGCAATTATCCTCTCCTCTGACTCGCAAGCAGGCAGCCAAAGTTGCGG

CAGAAAGTTCAGAGACTCTGATCAGTGGAGGCCTGTGCACTCCTGGCAGATCACGAAAGTCTGCTCGTTTGAGTCTACCC

ACTTGTGTTGCTCAGCAGCAGGGCTCCACACAATCACCTGTTGAAAGGCGAGTCACCAGACGAAGCTTAGCCATTCAGTC

CTCTTCTCCTGCTTCTGGGGCTGTGAGCGCCCGCAGACAGAGCCTCAGAACACAAAGAGTCAGATCCTCTACAAGCTCTT

CCCCTTCTACTTCACCCGTTACTCCCAGACGACTCACAAGACTTAGTGCTCAGTTTGGTCTGTGCAGTGATACACCAGCT

TCTGACTTCCCTTCTCTTCCTCCAAGTGCTACTAAAAGGAGGAGAATTGCTAGCGGCAGGAGCACCAGCTCGCTAGACTT

GGTGGACGGCCCAAGCCCGGCGAAACGGGGCCGCCGAGTGTCCAGCAACACATCCACCTCGTCTGGCGGCGGTGGTGTGT

CACAGTCACACGATTCCTATTGTGTTGCTGGGGCTGAGCTACCCAGTTCATCCTCGTACTCCTCCCAAGGTATTAGGTCC

CGCTCTCTCAAGGTCAACTCTTCCGAAGGCTTGAACGTCGCGCTGGCTTCTTCACCTGGTAATTCGAGCTCCTCTCCACC

ATCCTCTTCATCCTCCTCATCCACTTCAGGCGTTTTCAGCATCACCAACCTTGCCGAGTTCACTCACACTGTCGCACCTG

CATCTGCTGCATCCAACTCAACAGGTGCCTCTAGTAACCCTGCAGGCAGTTTGCTGGACCTGCCTCCTTCTGCTCTGCCT

CTGCCACAAGAGCCAGAGCCACGTCGCAGCTCTTCTGGCATAGTCAAGTCATATCCCTTACGCAACAGGCTACGCAGTGA

AGCGCACAGCAGTTACTGTGATCCTACGAGTGTATCTGCTGCTGAAGCTTCCCCAGCGTGTGGGGCCTCCACTGCCGCCC

CCGTGTGGTATCACCAGAGCAGCCTCAATACCGTGTCATCAGCACTAGGTGCTACTGGCCTCGCCACTGTTGGTAGCGGT

CTCTCCAGTGGTAGTAGTGACTTATCACACTTTGCGCCTCCTGTACCTCAGCAGTCTCACTCCACCACAGTTGGGAGCAT

TGGCACTCCTGCTGGTGTTGCGGCTGCTGCAGGCTCTTCCAGTGGGGCCATTGATTTGTCTGTTCACTCCAGTCGTACCG

CGGGAGTTCTTGCAACCTCAGCCACTCCCGCCGCTTCTGGAGCCCCTAGGTACTCTCACCAAGTGACCCCATCCCTCCCT

TCCTACCATTACACCTCCCACCCACCCCTTCCTCGGCACCAACCCCCTCCTCCTCCCCCTCCATCCCACTCGCAAGCTGG

TGGGTACTCTCTGTTCCCTCCAGCATACCCTGCAGCCCAGTACCAATATCCCTACGGTAGCGGAGCACAACATCAGCCCT

ACCCCCCACCACTGCCACTCAACCCCGGGGCGCCAGGTCATCTCCTACAGCCGGGTTATCCTCCCATTAGTGGTCCCACA

GCTGGACTGTACCGCGCTGCAGCAGTGCCCTCTGCTACTGTCACGGCCCCTCCCCCACCTCCTACGGGAGCCACCAGCAG

CAGCAGTGGAGGTGGCAGTTTCCTCACAGACACTTTTAGAAGATTGAGGGACTCCCGTCGCTCTCAGCACCACCACCAGC

AGCAAGCGGGTGATCTTTTATCCTCCACCACCATAGCTGGTGGCCTCATAGCGACAAGCACTCCAGGACGTGGTTCGTCG

CCATCATCTCGCAGGGGCTCGGGCAAAGAGTCCAAGAGTTACTGGCGCAGCAGATCCAACACCGTCAGTGGCGGTACCCC

ATCTAGCGGCGCCTCCTCGAGCGGTGTAGCGGTCAGCGGCAGCAACTCGGGCGGCGGCAGGAGATCGTCTGGCGGCAGCG

GTGGTAAGAGATCCAGCGGTCGCCACAGCAACTCCAACAACAAAAACACCACCAACACGGGGACCTGTGCCAGCACCAGC

CAGCACTGGTCAGCATGTGCTGGTAAAGGCGGCGGCAGCAGCAGTAGCAGCACCAGTAGCAGTAGCAGTGGGGCAGCCGG

TGCTGCTGCATCCGCTCTTGGACTGGAACCTCCTCCTCCAGGCAGTGGACCTTCAGGTACAATGTCTGGAGCGCCCAACA

GCAGCAGCGGCGGTGCTGCTCTGGGTGCTGCTGCTAGTGCTGCTTCTAGTTCTGCCTCTGCACCGAGCGCTTTGGGTGCT

GGTGCTGTGGGGATCTTGCTGGATCCTGGGGAAGATGACACTGAAATGGGGCGACTTGAGTCTCTACTGGCAGCCAGAGG

CTTGCCTCCCTCTCTTTTTGGAAGTCTCGCGCCGCGCATGCATAACTTGCTAAACAGATCCTCTACCTCCACTACCATAG

GTAGCAAAGCCAACCAGCTGCTTGCGGGTCTACAAGCTACTGGCGACGAGGGGCAACAGTTGCAGGCCCTCATCGAAATG

TGTCAACTGCTTGTAATGGGCAATGAGGATACGCTTGCAGGCTTTCCTATCAAGCAGGTCACTCCAGCCCTCATTACCTT

ACTCAACATGGAGCATAACTTTGACATGATGAACCATGCCTGTCGTGCTCTCACATACATGTTGGAGGCGTTGCCTCGCT

CTGCCAGTGTCATCGTAGACGCTGTGCCAGTGTTTCTGCAGAAGCTCCAAGTCATCCAGTGTATGGATGTGGCAGAGCAG

AGCCTTACTGCCCTGGAGGCCCTCAGTCGCAAGCACTCCAAAGCCATTTTGCAGGCGGGTGGCATTGCCAGTTGCCTGAT

GTATTTAGACTTCTTCTCGCTGCCTGCGGCTCGTGCGGCGCTCACCATCACCGCCAACTGCTGTCAGAACATCACTCCTT

CCACCTACCACTTCGTTGAGGACAGTCTACCCATCCTCGCCAATAGGATCTCCCCAACTGGTGACAAGAAGTGTGTCGAG

TCAGCGTGTCTTGCGTACTCTCGGCTCATCGACAACTGCCACCACAACCCTGAGAAGTTACGCCTTATTGCCAAGCATCA

GCTGCTGAGTAACATGCAGACTTTGGTTGTGGCAGTACCCAGTCTGCTGAGCAGTGCTAGCCTGATTCTCGTACTGCGTC

AGATGGCAGTGCTTTGCAGCAACTGCCCTACCCTCGCTGTAGAACTACTCACCAATAATGTTGCCGAGACGCTCGTTCAG

CTCATCACAGTTAATGGCAGCGCCGGAGGAATGGACGGTGAAGAGTTGGAGATCAGCGCGAGGACCCCTCAAGAGCAGTA

CGAGATCGCCAACCTGGCGTGTGAGCTGCTACCAAACCTGCCCAACACGGGCATGTTTGCTGTCGACTACAGCCTCACTG

CCACCTTCTCTGCCTCCTCTGTCATATCTTCTACCACCTCGGGTACCAAATACACTGCTCTTCCTCCCTTGCCCGTGTCT

TCTTTGCCTCCTTCTCTCCCTCCTATTGTTACTCATCCTCCTCCGCCCCTGCCTAACATGACTGCCAGTGCCCCTCCTTC

TATTGCATCTACTGCCAGCCTGCCTCCACCTCCTCCACTTGCTGGCAACGGAGGAACTGCTGACGTTCTGCCGCCGCCAC

CACCTCCACCTCTTCAGTTACCTCCTGCTGATGGCATCACTCAGCAAACCCCATCAGTCGTCATGCCTCCATTGCCTGGT

GACGACAACTGTCCCTTGCCTGCTCACCTTCCTCCCCCACCTCTCACCAGCTTGCCTCCACCACCGCCTCCACCTCCTCC

CCTCGCTACAGTACAATCATCTGATCCTACAGGTGCCGAGACTGGCAATGGTAATATTTGCGTGACGTCGTCAATAGCGT

CTGGCAACTCCATCTCACCTGCCGTGTCCAGTTCTGCCGCTGCGTCCTCGGGTGCCGTGTGGCAGTGGCAGGATGGCAGA

CAGGGCTGGCTGGCCCAGCAGCCTCCACCAGCTCCTGAGCCTCCTAAAACTCAGGAGGAAGATCCTCGCAAAGTATGTTT

GGAGACCAACCCCGATCTAGCTGCTAACTTGTTGCGTCGGGTCTTCAATTTACTTTACGAAGTCTATTACAAGAGCGCGG

GGCCTGCCGTCAAGCACCGAACGCTCAAAGCTTTGCTACGCATGGTGCATTTTGCTGACCCTGAATTGCTCCAGGAGGTG

ATAAGACCGCAGCACTTCTCGTCTCAGCTGGCGACCATGCTGTCCTCCAGTGATCTCAAGATTGTAGTGAGCGCTCTGCA

GCTCGCTGACCTGCTGATGAAGAAGCTGCAGGAGGTGTTCTCAGTTTACTTCAGAAGAGAAGGTGTATTGCATCAGATCA

AGAAGCTGTCAGAGCCACCGGTAACTCCTGTGGTGCGTACTCTGAGCAGCGGTATGAACACCCCCACAGCTACACCTCCA

GCGTCTCAAAGGAACAGCCTCATGGCCGGTGGAGCCACTCCTCTGCTTGGTTCCACCAGTGGCAGCAATTTGTTAGGGCC

ACCCATGCCTGGCTCATTCGGTACAGCCGGGTCGCTGTACGCCGGCGCTGATTGGCTCACTGTTGACCAGAGCGTGGGCG

GGTCGGTGGCTCTCTCAGCCAGCCCATTGCTGGCTGCTGCGCGCTCCATAAATGACAATCTCGGCGGTGGCGAATCCAGC

CTCAGCTTGTCCTTACATAATGCTTCCATGCCTCACTTGGCCTCGGCCTCTCCCTTGACTGCCGGCCATCTGCCTCCACC

GGTAATAGGATCGAGTTCTGCTAGTTCTTCATTCCACTCGTCCCTCACTCCCTCTCCGTCTCCCAACCCATCCCTCGTCC

ACAGCTACCCACCCCATGCTTCCGACGGTCTCGCCCCCATTTCGCAGCTGCTGGAGCCTAACATATCTGCCAGCAGCCAC

ATGTCTCCTCTTGTTGCTGCAGGTGGCGTGACGCCTCCAGCCCACAGCACCAGCAACTCTGGCAAGCTCACAAGCTCCCC

GCCCATGCGGTTGTCTGAGGTGCTCAAGAAGAAGCGCTCCAGTCGGTTGAGTAGCGGTGGCAGCGGTGGTGGAGGTGGAG

GAAGGTGGAGGAGTAGGAGGCAGGAAGAGTCTCACCAACAGCAACAACAACACGCTACTTCATTCGGCGAGTTCTTCTTG

AAATCCCTGTCATCTACAGCCAGTAGCCCCGACACTCCCGTCAATACAAGCAACAGTCGTTCCTCAGCATGGCACATGAG

CGGCTCCAGCTCTACCCCTCACCATGCAGGGAGCCGGTCTGCTTCTAAGTCACACGGTGGAGGGTCCAGTAGTGGCAGAG

GAGCTGGCAGCAGCAGGAGCGGTTCCAGTTTCCTTGCGAACCTCAACCCTGTGCGGTGGGGCCGCTGTAGCTCCAGCACG

GCTCCTCCTCAAGAAGTCAAGGAGAAGACGGTGAGCAGCAGCAATAACAGCAACAACTATTGCTCTCCTGCTCCACTTCT

GCCCCACCACGCTCAGGCGAGTTGTACTAGCAGCAGCAGCCGGCACAGAGAGAGTGTGCGGGCGTGGATCAGAGAACAGG

CCATGAGACTCGATAGGGAACACTTCGGCCTAGAGCTGCAAGGTCTGACGCATCCTGCCCTCACTGTCCTCAACAGACTC

ATTGGTGCCATACAGCAGCTGCACACCCAGCCTGGCAATTGCATTCCTGCTCTGACCGAGATGATGCACATCGTCACCGG

CTCTGACATCTCCTCGTTCGAGCTCATCCACTCGGGTTTGGTGAACACCTTACTGCTCTACCTCACGGCTACCGACAAAC

GAACAGATAAGGAGCTCACAAAAGGCAGCAAAGATTCTTCTGATTCTGCTGTGGCTGCGACTTCCAGTACCACCGATGCT

GAGGACGGCCCTGGCGTCTCCAGGAGTGGCAGCACTAAGGGCGGCAGCAGCAGTGGTGGTGGCGCGAGTCCCATCAACAG

TGGTGCATTTGAAGCAAGCGGTGGTGCCTCACTCGACCCTGCAGTTGTTGCGAATATCATGAGAACGCCCAGAGATGAGC

GCCTCAGGAGCTTCTTGCATGTCTTCCTAGGCTGCTCGCTGGACCCCCTAGAGCCTGGTGGCTGTTGTGACCCAAGTCAA

GTCTCTCGATTTACGGGGCTCCTGTCCAAGTTGATGTCTTGCGTCAACCAGCTGGAGCAGTTCCCTGTCAAAGTCCACGA

CCTACCGTCTGGTGCGGGCGCTGGCTCTCGTGGTGGTGCTACTTCTGCCATTAAGTTCCTCAACACACACCAGCTCAAGT

GCAACCTACAGCGGCACCCTTCGTGCACGAGGTTACGTGAGTGGCGTGGAGGTCCCGTGAAGGTGGATCCGCTGGCGCTG

GTACAGGCGATAGAGCGCTACCTCGTGGTCAGGGGCTACTCCAGAGTACGCGACCATCAGCACGACGACAACAGCGAGGA

TGACAACAGTGACGATGATGATATCGATGATAATTTGGCGTCCATCTCGTCAAGCCAAGGTAACCAGCAGCACAAGTTGG

AGTTCCTCATAGGCGACCAAGTGCTGCCCTACAACATGACGGTGTACCAGGCGCTGAGACACTACTCTCAGGAGCTGTCC

CCTAGCGATGCCGAGAATGAGGGCGAAGTGCCGGTGTCGCAGAGCGCTGTATGGCTACACACCCACACCATTTATTACCG

ACCTGTGTCAGAAGAGCCGCCCGTCAAGTCATCCCGCAAGGGCAAGGGAGGCTCCAAGTCCAGCCCCAAGAAGAAAATGA

TCACGGAATCTCTCGTTACAGATGCGGCACTTATTGGCGGCAGCTCAGCGGTGTTGGACTCGCTATTGTCTGCTTCACTG

CCGTCCTCCGTAACTGTCAAGGACCCTTGCCTGGAGGTGCTGGCGTTAGTACGAGTACTGTCAGTTCTCAACAGACACTA

CCACAGCCTCTACCCTCCTGCACCTACCAAGCTGCCTGTGCCTATAGCTGATTTCCAAAATGTTAAGCTTACTGCCAAAG

CCAACCGTCAGTTGCAGGATCCTCTTGTCATCATGACAGGCAATCTACCTCCATGGCTACCTCAGATCGCTTATGCTTGC

CCGTTCCTATTTCCTTTTGAGACGCGCCAGCTGCTGTTCTATGCGGTCTCATTTGACCGCGACCGCGCCATACAGCGGCT

ACAGGAGACCAATCCCGAACTTAGTCACGGCGACTCTACGGAGAGAGTCACCCCGCGATTGGATAGAAGAAAGAAAACAG

TCAGTCGAGAAGATATATTAAAGCAGGCGGAGAGCGTCATCAATGACTTGGTGCCTTCTAAAGCATTGCTGGAAATCCAG

TACAGTGGAGAGGTTGGCTCCGGCTTAGGACCAACCCTAGAGTTTTATGCCATTGTATCTAGAGAAATGAAAAAAGCGGA

GCTAGAGTTATGGAGAGGTCAGTCTGTTGATGTAGAGGAAGAGACTTTGGATGGTACGTCCAAGACGGTGAAGTATGTCA

AAGCAGACACGGGTCTCTATCCTATGCCTGTGCCCAAGAACATGAAGTCTGCCGCCATGAACAAAATTAAGCAGAAGTTC

AAGTTTCTTGGCAAGTTTATGGCGAAGGCAGTCATGGACTCGAGAATGGTTGATCTGTCGCTCCACGAGTGTATGCTCAA

GTGGATGCTGAGCGAGGAGCGCTCACTAGGACTGGCCGACGTGCTGTCACTGGACCCTGAGTTCGGTGGTACGCTGCGCC

AGCTGCACGCCCTCGTCCTGCAGAAGAGAAGAGTCCAGGACCAGGCCCGGGTGCAGGGTTACTCCACACAGCAGCTCGAG

AGCGACGTGAGGAACATAACGCTGGATGGGTGCAGTGTGGAGGACTTATCCCTGAACTTCACCCTACCGGGTTACCCGAG

CATCGAGTTGAGGAAAGGAGGGGGAGACACCTGCGTCACTATTGATAATCTCGATGAGTACTTACAGCTTGTGGTGGAGT

GGCTGTTACGCGATGGAGTGTGGCGGCAAATGGATGCCTTCAAGGAAGGATTCGACAGCGTCTTCCCAATTAGTCAACTT

CAGGTGTTCTATCCAGAGGAATTGGAACAAGTGTTTTGCGGCAGCGCTGCTACGTCTCACTGGGACATCAAAGTCTTGGC

CGAGTGCTGCAAACCTGACCACGGCTACACGCACGACTCCAGAGCTGTCAAGTTCCTTCTTCAGATTTTGTCGGAATATT

TACCGAAGGATCAAAGAAATTTCCTGCAGTTCACCACTGGCTCGCCGAGATTGCCAGTTGGAGGGTTCCGCAGTCTGACT

CCGCCTCTCACTATTGTCCGCAAGACGTTTGAAGCCAACGAGAACCCCGACCACTTCCTGCCGTCTGTGATGACGTGTGT

CAATTACCTCAAGCTCCCGGACTATTCTACCTATGAAATAATGAAAGAGAAACTCTGTTTTGCTATTAGTGAAGGACAGC

ATAGCTTCCATCTCTCCTAAAGATCTGCATGACCCATGTTCCTGTGGCGTCTCTGGCTCAGATTGACTTCTCCTGCGGTC

ATCTCACTGCCCTCTGACAGGTGCTGCGTTAGTCGAGAAGCTCCCTTATCCTTTCCCGTGGTGCATTTTTCGCTTGTGTT

CATTACAGCATGAGGGAGCTCATGCTTAGGTACCTCGCCCCAGTAGGAGTGCTGAGTTCTGTATGCTGCAGATTCCCTTT

TTTTTATTTGTTGTAACTTAATGGACATTTAATGTGAAGTATGACAGAATGGATCTATGTTTTCTTAAGAAATTAATAAA

TGAAACGATCTGTTGGTCAGTTCTTGTTGACTCAACTCATTTAACTGGCTGTGTGGCCAGCTAATTTAATAATTTACGAA

GCATCGATGGGCTGATTTTGCCGCAGAATGACTGAGTTTTTCTAATTTTCAGTATCAATAAAAAGGATTCGTGCATTACG

ACCCGCTGACATTGGAAGAGCTTGATCATTCCTTATGTAGCTACTTTTCTAGCGCCTTGATTTGCAGCCCAGTAGCAGTC

AGCTGTACCGCCGCAGTGAGATGATACGCTGGACTGTCCAGCAGGCAACGCGGCATGTACTTGCTACTGTGTAGCTACGA

CCGACGGTAACAGAGCTGGTTTGGCGTGACCGCACCACCACCACCACGTGATCACCTCACTCTGGTGTCGGTCTACACGC

GCGCTTGACGTGCATATGGGGGTGTGGTGACATGTCCTATGTCATGTAGCTAATTGCTGAATGTGATGCGCTAAAGCATT

TTGTATAGTTTCTTGCATTATAGTGCTGCCGTAGTCTTATGTTCAATTATTGTAGAGCCCTTGTACATCTTGTAAATAGT

GGTAGTCTGTAAGTATACTCTTTCCCTACTGTCAATATCTTTGTAGAGCTCCGGCCTGGGGACGGGCAAAGCCTTCTCTT

GTTCTATCTTACTGTCTCTTTTGTGATCTATCAGTTGTGTAGTTCGAACTGAGGCTCTACTGATACTCTCATTTCTTACC

TGTGTTGCTGTTATGCTGCGAGTGGGCGGCGTTTATCCCCCCAAATGTACGCAGTTATTCTGTGAATCGGCAACGTGATG

CCAAGCTGATTCCGGCATTAGCACTGACTGTCCTGACGATATGCGTACGTTGCTGTGCTCGTATTACCGGGAACTATACA

TAGAGCATCGTCTGCTACCTTCTTTCCTAAACCATCAACACAAATGTACTTGTTTTGCTTTGTATGTGGGCGTTTGTGCT

GCTTTGTTTTGTCTTTATTAGAACTTTCAAGATGGTGAAACTTCTTATTGCTGCGATTTGTGAGATCTTGCGAACAGTTG

ATGCCGCCGCGTTACTTGATCTGATTGCCATGGCAAGCGTTATTCCCGACGGTGGTGTGGCCCGCAGTTCCCTGGGAAGT

TTGGAAAGCATACTTTTTATTTCATTTTCACAGTTAACATGAAGTGCCCTTTTTAGCTTATTGTCGAGTACTAAATTCAG

CTCTACCACCATCGCAAAACTTGGAGACCTTGTATATTAGTGAAATTTGCACTGAACCGAAGCTTCACTATCACACGTAG

TGCTGTGTGCGCTTGTGTACGAGTTATTCTTTTGCGTCTTTTTATCATATTGCTTGAAGTAATTGATTTAATTTTGAGTG

TTCATAGTCTGATCATATTGCTATGGTGGAAGAGGAGCACAGTTGCCGTAACGCTCGCGTGCTGCCCAACCATCGTATTG

GCACTTTCATTATTAGCTGTACTTTTCAATGCTTGTTCAGTGTTGTAGCATTATCATATTTCACGAGTGTGTTTGTACAG

TAGCGCCGTAACGGTTTTTTATTGTCTTTGAAGTAGAAAGTAGCTGAAGGATATCTACACATTAATTTCAGAATTGGAAC

ATAAGCATTGTTTGAGATAGAGGTACAATTTTAGTTGGCTACGGTGTTTTTTGTATGCTGCTCGGATTTCACTTGATTTT

TGTCCCAACTCTTGTGCGACCAATAGGTGCTTCTAGGCAGATAGGTACACCTAGTGCAGCGCGGCGCCGGCTGGGGCTGT

GATTGAGCCGCTGTTCCCGTGATCTCCCTGGCGTATGCCAGGCTCAGAGCTGGGCGGTAGCGCGCTCTGCTTCTTTCCTT

TCTTTCATTCGCTGCTCTTCCTGTTTGCCATCCAGTCCCTAGCTAACGATTTCCTATTGCTGTAGCTACGTTTCTATTTC

GGCCCAACTCAGAGCTTAAGAATTCCGTAGTTCGAACACGTAGATCAAGTGATGTCGTTCGCATGCATCTCCCTTCTGAG

TTACAAGCGCTGGTGCCACAATGTCGTGCTGCTTCCGCTTTCCGGCAGACAATGTATTTTTTCTATTCTGCTTTATCCAT

CTTTACCTCTCTACTACTACTACTACTACTACTACTCCTCCTCCTCCTCCTCCTCAGCTCTGCAGCAATGTTTGCATTAA

CATAGTATGTCCAAATGCTGGTATCAGTTTTTTCGCAATGCCCCCTTGTTACACTGCTGAAATTTACTTGAATATCTACC

CAGTTGCCATGCTGCTTTTTTCTCTTGAGTTACAGTTAATCCTTTTGCTCAACCGTATAGCTCGTTGCAGTACGCAGCGC

CGTTATTTATTCATTATCCGTTGTTAACCAAATCCCACCCGTCTGGAGTTGTTATCAAGCGCTTTGTTCCATTGTGAAAG

TTGTCGTTTGTTTGGAAGGCGTCCCGTGGGCGATGAGCGCCGCTCCCTGAAGTCCGAACGCTCCTGCCGCATCTCCACTG

GCGGCTCTTGTTGGTCGTAGCCACGGCGCCCCTTGTTCGTAGTCGCTCGTTGTTGTGTTGCTGAATTTCTGTTTGACTCC

AAGTGTATTTTTCTTGCCTCTTTTATTGTGTTAGAGAATAAAGTTCTTTTATTGTGTATCAGGATAGCTTCGGTTTGTAT

CTTTGGCATTACATAAATATATCGAAAACTAGCAAAAAAAAAAAA

Doubletime

>lcl|PH.k27.comp42178_seq0

CGTTTAGTGTGAGCGGCTGTGTGCATTCAGTGGTCCGTGCAGCTAACTCCTTCACCTTCTGGTGTCTATAGATGCATTAT

CCCCCTACGATATGAGATAGGAATACGAAAACGCTAGAATTATACAACCCCGCGTCATTTGCCAAAATAGCTTAGGATGG

AGTTAAGAGTTGGAAATAAATACCGCCTTGGACGGAAGATTGGTAGCGGATCTTTCGGCGATATTTATCTCGGCACAAAC

ATATCCACAGCAGAGGAGGTGGCCATCAAGCTAGAATGCATCAAAACAAAGCACCCACAACTCCACATAGAGTCCAAGTT

TTACAAGATGATGGCCGGAGGTGTAGGCATACCCGCCATCAAGTGGTGCGGCAGCGAGGGCGACTACAACGTTATGGTAA

TGGAGCTCCTCGGTCCGTCTCTCGAAGATCTCTTCAATTTCTGCTCCAGAAAATTCTCCCTTAAAACCGTACTGCTTCTC

GCTGACCAACTTATCACAAGGATAGAGTACATCCACAGCAAGAACTTCATCCACAGAGATATCAAGCCTGACAATTTTCT

CATGGGCCTGGGCAAGAAGGGCAACCTCGTTTATATTATTGACTTCGGCCTCGCCAAAAAATACAGGGATCCTCGGTCGC

ACCAACACATTCCTTACCGGGAGAATAAAAACCTCACCGGCACCGCTAGATACGCCTCCGTCAACACTCATCTGGGCATT

GAGCAAAGCAGACGAGACGATCTGGAGAGCTTAGGTTATATTTTAATGTATTTCAACCGGGGGTCGCTGCCCTGGCAGGG

GCTCAAGGCCGCCACCAAGCGCCAGAAGTACGAGCGCATCAGTGAGAAGAAAATGCAGACGCCCATTGACGAGTTGTGCA

AGGGCTTCCCTGTGGAGTTCGCTACGTACCTGAATGTGTGCCGTACGCTGAGGTTCGAGGAGAAGCCCGACTACAGCTAC

CTTCGGCAGCTGTTCCGCCAGCTATTCCACCGCCAGGGCTTTACATACGACTATGTCTTCGACTGGAACTTGCTCAAATT

TGGTAAGGGCTCCAGAGGGGGAGACAGCGACCACAGCGGCAGGGGGACGCACTCCACCTCCCACCAGAAGCAGCTACAGG

GGGGTACGCTGCCCTCCGTCAGAAGCGCCCTGCTCAGCCACATGGTCAACAGCAGGGCCCGCCACGACACTGGGGCCGTG

CCTCCGCACGGTAATGCTAACAGTCCGCCCATCTACCCCGGCACGGACCAGCCCACCCCAATGAGCTCTCTCCTGGACAC

CTCCTCCCCGGTCCTCTCCGCCTGCCCTGACCTTCCCCGAGACAGCCTGAGCCGGTCAGAGCTGAGGTCACTGAGACACC

CAAACCCCCAATGGCGATCTGGCGCCCACTTCCGTCTGCGGCCTCCAGCCTAGTGCCCACCTGCCCACCGCTGCAGGCAG

CCTAGCGCCCACCTGACAACCACTGCAGGCAGCGTAGCGCCCACTCCTTCCCTTTGCCCACCAGAACTGTTCCCTCTATT

CCATCAGTATACTCCTGCCTCAGATGAAATGCGTATCTCACTTTCTATCCCTAGGACGGTCTGTACGTTTGCATGGACG

Nejire

>lcl|PH.k51.J3947083

CGGCCCCTTGGGCCTATTATGTTGAAGCCCATGGCTCCTGGAGCTGGAGCTAATGGCGGCCTAATCAGAGGGGCTGCACC

TCCCAACATTATGGTGTCGAGCAGCCCTCTGCAGCAACAGCAGCAGGTCTTCACCAGCTCAACTGTAACGTCTCCTGCCA

ATTTACCAGCGTCCGTGCATTGCTCCACAGGCATGTCGCCTAACTTGCCGTCATCTGTTCATGCTGGCGCCTCTCCTGCC

AATATCCCTAATTCTGTGCTTGGCTCAGCTGGGACGTCTCTCGCATCGAGCGGTTTCCCTGGTGGTGTCAGCACCATGGC

ACCCTCGGGTGTTGCGCCCAACTCTGCCGTTCGTCCCCCACTGGCTCCTCCGCCTTACACCTCAACTGCCATTACTGCCT

CGAGTGCCACAGCTTCTATCAAAGTGCCTGTTACTGGCGAAATGAACTGCAGTGCGCCTGCTGCTGTAGCACCTTCTTCT

GTTGCATCTTCACTGCAAGTGTCGCAACAGCAACAACAAAACCTCGTGTGCTCCGCCAGCCCTTCTGTTACTGACGCCAA

TAGTTTCTCGGGCACCCAGTCTTCCCCCAATACTGCAACGAGCGCTGCGTCTAATGTTCTTCCTGTGTCAAGTGCGCCGC

TTCCTGAAGTTAAGCACGAGCCCACTCCCTTCATAAAGACAGAACCAATGGATGTTGATGTGAAAGATGAACCTGCCTCA

ACTGACGGCAACACTACTGCTCCACCAGCAGACGTTAAGGTAAAGATGGAAATGAAGACAGAAGTAAAAGACGAACCTCA

GTCTCCCTCAGCTGGTGGCGGTCTTAGTACTGACGTGCCTGTCAAAGAAGAGGTGAATGTGAAAATGGAGACGAGCAGTC

CGTCTCGGTCTGATGCTGGTTCTCTGACGCCTGCCATTTCTACCGCCACGACCACACCCGCGTCATCGAGTGCCTCCTCG

ACCGCCAGCACTAGTGCCCCTCCTACCAGCTCAACCACAACCACAGCTTCCACCCTCGCCATGGAGAAGCCCTGTGATGC

CAGTTCGAGCAAACCGAGTGGTGCTGCCTCAAAATCCTCGGGAGGTTCAGCCACTGGCACCCCCACTCCCCCCACCAGCA

CTGCCCCCAGCGTAGGCAAAGCTCCCACTAAGGATTTCCCCCCTCTTTTCACACCAGACGAGTTACGGCAGCATTTGGCT

CCTACCATGGAGATGCTGTACCGCCAGGAGCCTGACAGCATCCCCTTCAGAATGCCCGTGGACCCGTCCCAGCTCGGCAT

ACCCGACTATTTCGACATTATTAAGAAGCCTATGGATTTGTCCACCATCAAGAGAAAGTTGGACACGGGGCAGTACGCCG

ACCCTTGGGACTATGTCGATGATGTTTGGCTCATGTTTGATAACGCCTGGATATACAACAGGAAAACTTCCCGCGTCTAC

AGATACTGCACAAAGCTCGCCGAGGTGTTCGAGCAAGAAATTGACCCCGTGATGCAGCAGTTGGGCTACTGCTGTGGTCG

CAAGCACACCTTCAACCCTCAAGTTCTTTGCTGCTATGGCAAGCAGCTCTGTACCATACCTAGGGACGCCAAGTATTTTA

TGTACCAAAATAGATATACCTACTGTTACAAATGTTTCAATGACATACCCGGCGACGTGGTGGTCCTAGGAGACGATGCG

AGCATTTCTGGCATGAGCATCAAGAAGTCCCAGTTCCAGGAGCTGAAGAACGACGCCCTGGAGCTGGAGCCATTGGTACA

GTGTGTGGAGTGCGGCCGTAAGCAACACCAGATCTGTGTCCTCCACATGGAGGCCATCTGGACCACCTTCACTTGTGATC

TTTGTCTCAAGAAGAAGAGCCAGACGAGAAAGGAGAACAAATTCACTGCCAAGAGACTGCAAACCACCAAGTTGTCCAAC

TACCTCGAAACCAGAGTTAATAACTTCTTGAAGAAGAAAGAAGCTGGAGCTGGGGAGGTGGTTATTAGAGTGGTGTCTTC

TACAGAGAAGACGGTAGAAGTCAAGCCCGGCATGAAAGCGAGGTTTGTGGACACCGGTCAACTGCCTGCTGAGTTCCCCT

ATCGAGCTAAGGCTCTCTTTGCTTTCGAAGAGATCGATGGAGTGGATGTGTGCTTCTTTGGCATGCATGTGCAGGAGTAC

GGCTCTGACAGTCCCGCTCCCAACTGCAGACGAGTGTACCTTGCGTACCTTGACTCGGTACACTTCTTCAAACCCCGCCA

GTACAGAACAGCAGTGTACCACGAGATACTCTTAGGGTACCTCGATTATGTTAGGCAACTAGGCTACACAATGGCCCACA

TTTGGGCATGTCCTCCTTCAGAAGGGGACGATTATATCTTCCACTGCCACCCGCCCGAGCAGAAAATACCTAAGCCCAAG

CGGCTGCAGGACTGGTACAAGAAGATGCTCGACAAAGGCATCATCGAGCGTGTGGTACTAGATTACAAGGACATCCACAA

GCAGGCGTTTGAGGACCACGTTAAGTCTCCTGCAGAGTTACCGTACTTTGAGGGCGATTTTTGGCCTAACGTGATGGAGG

AGAGCATAAAAGAACTAGATCAGGAAGAGGAGGAGAAGAGAAAGCAGGAGGAAGCGGCTCTTGCTGCCGAAGCCGCACAG

GCTGCCATGGGCAATGAAGAAGAAGAGGAAGTTTGTCTTGATGGCAAGAAGAAAGGCCAAAAGAAGCACAAAACCAAAAA

TAAAAACAAGAGTCAAGTTAAGAATAAGTCCAAGAGTAAAAGTAGTGCTCAAAGTTGCAATGATTTGGCCCAAAAAATAT

TTGTTACAATGGAGAAGCATAAAGACGTGTTCTTCGTTATCAGGTTACACAGTGCCCAGAGCGCCGTTAGTTTACCGCCC

ATCCAAGATCCCGATCCTACAATTGTATGCGATCTCATGGACGGTCGCGACGCCTTCCTTACTTTAGCTAGCGAGAAACA

TCTCGAGTTTTCTTCCTTACGACGTGCTAAGTTTTCTACCATGACAATGCTCTACGAACTGCACAACCAGGGTCAGGATA

GGTTCGTCTACACGTGCAACAACTGTAAGCAACATGTCGAGTCTCCCTACCACTGCCAGGAGTGCGACGACTTTGACTTG

TGTACTGCTTGCTTCAACAAGGATGGTCATCAGCACAAGATGATCAAACTTGGCCTGGACATGGACGATGGTTCAGGATC

TGCCGACAGCAAGAACATGAACCCACAAGAAGCACGGCGGAAGTCCATCCAGCGGTGCATTCAGAGCGTGGTGCACTCGG

CGCAATGTCGCGACGCCAACTGTCGTTTGCCGTCTTGCCAGAAGATGCGAAGAGTTTTGCAGCACACCAAAGTGTGCAAG

AAGAAGAGCAATGGTGGCTGTTCCATATGCAAGCAACTCATTGCTCTCTGCTGCTATCACGCACGTATCTGCAAAGACCA

GCGTTGTGTTGTGCCTTACTGCTGCAATATCAAACAGAAGGTGAGACAACAGGAAATGCAGCAGCGATTGCAGCAGCAAC

AGCTCATGAAGCGAAGAATGCAGCAGATGAACGCTGGTATGCCTAGCGTGGGCAGCAGTTCCAGCTCTCCTCCCAGCACC

AGCAAGCCTGTTGTTCAAGCCTCACCTACCTACAACCAACAGGAGGGCTCGACTGCATCGTCTGCCGCTGTTCCAGCTGC

TGGCCAGAATGTTGCCACACAGGCGCCTCCTCCTAATGTCCTCGAAGCTGTAAAGGAGGTGCAGGCGGCGGCAGCACGAC

AGGCCGTCCCGAGCAACGCTGTACCCAGCTACGGTAAGGGCAACCCTATCCCTGGGCAGCCCGGTGGTCATGGGGTGATG

GTGAGTGTGTATCAGGGGCAGCAAGTTATGCAGCAGCAGCAGCAGCCACAAAGGCCGCAGCACGCGATGCAGTCACCACA

GCAGCAGCAGCAGCAACAACAACAGCAACAACAACAGCAGCAGCAGCAGCAACAACAGCAACAACAACATGGCGGCATGA

TGAGCAGCAACAGCATGGGCAAGCCGGTGAACCAAATAAGTGGTCCCATGGGTAAACCTAACGCCATGGTGCAGCAGCAT

CTCATGCAGCAACAGCAGCCACAGCCCAACAGAACCTTGCCTGCCATGGACCAAGTGTGGAACAGGTACCCCAACGCCTC

CTCCAATCAGACGAACGGTGCAGGTGGCATGAGACCAACCCTGATGCACAACCCTGCAGCTAACGGCTCAGTACCCGCAT

CGGGAGCTCCCAATATCCCTGGCTCCAATCCTGCGGGTGTAGTCACGAGCTCTGCTGCTGGTGGTGCTGCTGGGGGAGGT

GCCAACCACCCTGTAATGGGTCCTCCTGGACAACAGCCACAGCGAAATTCCACTGTAGTTTTACAGCAGCTACTGCAGAA

GTTCAGGAGTCCCTCAACTCCTAACCAGCAAAGTGAGGTGCTCGCAATACTCAAAAGTCATCCGCAGCTCATGGCCGCCT

TCATAAGGCAGCGCAATACGCATCATCAACAGCAGCAGCAACAGCAACAACAGCAGCAACAGCAGCAGGGCCAGCAACAG

CCACCGCAGGGACAACAGCAGCAGCAGCCCCAACAACAGCAGCAAAATCAACAGCCAACGCCACCGCCGCAGCAGCAGCC

TGGTGGCGGGATGTACCAGCCACAGCAGGGCGTTCCCATGCAGCAACAGCCGCAGCAGCCAGGTATGCAGATGCCCATGC

AGCAGCAACAGCAGCAAGGTATGCAGATGTCGATGCAGCAACAGCAGCAGCAACAAGGAATGTCCATGGTTGCCATGAGT

GGTAACCAGCCAGGACAGCAACAGGGGCCCCCGCACATGCCAGGAGGGATGCCACACGGCCAGATGCACAGCGCCCAGTG

GTACCAGCAGCAGAAACAAATGCTCGCCTTAAGACAGCAGCAACAACAGCAACAGCAGCAGGTCGGTTTCCAGCAGCCCC

AACCTCCTCTCAATGCCGCAGCCAATCAGCGGCGGCACTTTGGGCCGCAGTCGCACCCCAATTTCGCCAACCAGCCAATG

GAAGGCCAGCAATTTTCACCGATGGGCTTCAATTCTCAGCAGCAGGTGCAAACTCCACCGCAGCAATCAACTCAGCCTCA

GCAGAAGATGATGTTACAGCAGCAGCAAATGAAAGGCGGTAATTCTATATCACCCGGTCCATGTGTTACGGGCATGTCTT

CTCTACCTTCTCAATCACCGCAACAGCTTATGCAATCCGTTACGTCTCCACCTCCGGGGGGAGGTGGCAGCCTACAGCAA

GCTGTTAGATCCCCCCAGCCTTCCCCTAGACCCTCGCAAGCGTTGCCATCTCCGAGATCTGTCGCCGTCCCCTCCCCCCA

CCAGGGCAACCAAGTCCAATCCCC

Nemo

>lcl|PH.c118708_g1_i2

AACAACAACAACAACAACAAAAGGCCAAAGAGAACAATCACCCATCCAATAAAAATTCAGGAGCTCCTGCCGAGAAGTCC

GAAGATCGCCCTATTGGCTATGGGGCGTTTGGGGTTGTTTGGTCCGTGACGGATCCGAGGTCGTCCAACAAAGTCGCTCT

GAAGAAGATGCCCAACGTGTTCCAGACGCTCGTGTCCAGCAAACGAGTGTTTCGTGAACTGCGCATGCTCTGTTTCTTCT

CCCATGAAAATGTACTGAGTGCTCTGGATATCTTACAACCTCCTCCTCTGGAGACTTTTCAAGAAATCTATGTTATGACG

GAGCTCATGGAGTCTGACCTACACAAAATCATCGTATCTCCCCAGACTTTATCTTCCGACCACATCAAAATATTCCTGTA

CCAAATTCTTAGAGGTTTAAAATACTTGCACTCCGCGAGAGTAATTCACAGAGATATCAAGCCTGGTAATTTATTAGTCA

ACTCCAACTGTATTTTGAAGATCTGCGACTTTGGTCTGGCCCGTGTAATGGAAGATGACACGTCCCGGGACATGACCCAG

GAAGTGGTGACCCAGTACTACAGGGCCCCCGAGCTGCTCATGGGAGCCAAATATTACACGCAGGCCATTGATATCTGGTC

CGTGGGGTGCATCTTTGGCGAGCTGCTGGGCAGACGGATCCTTTTCCAGGCTGCCACGCCCATACAACAGCTGGACCGGA

TCACGGACCTGTTAGGGACACCTGACCCCACCGAGGTGCGCCACACAGCTGAGGGCGCCCGTAACCACGTCCTCCGCAGA

CCACGCAAACCGCCCGCCATCAATACCCTCTATAACCTGGGGGCCAATGCCACTCAGGACGCCGTTCACCTCCTCTCCAT

GATGCTCACCTTTGACTATGAGAAGCGGTACAACGTGCTCCTCTGCCTCGACCACCCCTACCTCGATGAAGGCCGCCTCC

GCTACCACTCCTGCATGTGCTCCTGCTGTGGTCCTGATGCCAAGAACCCCCCACGACCCCGCTCCCCTAACTCTGCCGTC

CACACTTGGGAGGAGAGCATACGAAAAACCCCCGGCTTCAGAGGTGCTAATAACCCCAATCTGGAGCCCACTGCCTGTGA

ACCCTTCCGACACACCTACGAAGACGATATCCGATCGCTGCCTGAAGTGAAGGATCGCATGCACCGATTTATTAGCAAAC

AGATGAGCAGAGACAAAATTCCTGTCTGCATCAACCCTCAATCGGCCGCCTACAAGAGCTTCTCCAGCGAAGCTGTTACA

GTAAATAGTTCTTGATTGCTCGACTGTGGCCCACCCATCAGAGCTGCCGCCCTCACCTCATAATTGGGATTAACGCAGGG

AGAAGTAGGCCTTCTATTGTGTAGATATACCAACCGTGTGGGTGTGTGTGGCGGGTCCTGTCCCATCTCTGCTGCCCTCC

ATCGCTG

Pigment dispersing hormone receptor

>lcl|PH.k21.comp140546_seq0

CGAAGGAAATGGCTGGACGAACTATACCGTCTGCTTTACTCCAGACGCGCGTGATTTGCTGAATCAGCTTTACTCTGGAA

CCGAAGAAGAAGCTCAGATGAAATTCCTCGTAGCCAAAGGGTCCAGAGCTGTCGAAATCGTTGGACTCTCATTGTCCCTC

CTCAGTCTTCTGCCCAGCCCCTCCTCAGTCTTCTGCTCAGCCTCTTTATTTTTAGCTACTTCAAATTCGTTGACTTCAGA

GGACCGCTCGCACTGCCCTACTTCAGGAAACTGCGCAACAACCGATCACGGATCCACAAGAACCTGTTCGCGGCGATGTT

GGTGCAGGTGACGGTGCGCCTGGTGCTGTACACCGACCAGGCCATCGTCCGGGGGGACTCCATAGGTGTGGGGTCATCCG

TGGCCCCCGACAGGAGAGGCATCGACTCTACCCCTATATTGTGCGAAACTTTCTACATCGCCCTGGAGTACGGTCGCTCG

GCGATGTTCATGTGGATGTTCGTGGAGGGCATGTACCTGAATAACTTGATCAGCGTAGCCTTCTTCCAGGGGCCACCGAA

CTACAGCGCTTACTACATGATTGGCTGGGGTATTCCC

Par domain protein 1 ε

>lcl|PH.k27.comp15075_seq1

GCTACTTGTTAAATAAGGTTGGTCTGGTGCTGCAGGGCCAGTGTGTCGCGAGGTGTTGTAGGAGGTGTAGTGTACGTGTG

TCTCGTTGTGTGTCGCTCCTGTGGTCAGCGCTAGTGAATATCTTCACTTGTTGCGTTGCATGAGTTAACTTGTTGCGTCA

TATTGGGTGCTGAGAGTGTCTTCAGCGAGAAACCATGGTCGCCGAAACGGTGAAGAGCTACCCATACCCTGCCCTGCTGC

CGCTCGCTCAACACGGTGTCAACTACTCCTGCGCCTCGACCTCCTGTTCACCAGCACTTCTGCCCCAAAATCTTGGCCAG

TCTTACCCACAGGCCCCAGCCACCATGGATCCAAACCATCTGAGCCGAAGTAGAGCCCTCCCTAATGCTGGGTCCACACC

TTCTGGCTTAGGTGCTTCTGGGATGATGGCCGCCACTCCTGCTTTGCTCAAAGAAACCATGTTTGCTCAACGTAAGCAGA

GGGAGTTCATACCTGACAGTAAGAAGGATGATTCCTACTGGGACCGACGCAGAAGAAATAATGAAGCCGCAAAGCGCTCA

AGAGAAAAGAGGCGATTTAACGATATGATCCTGGAACAGCGAGTGCTGGAACTCTCCAAAGAAAATCACATACTTCGAGC

TCAATTGAGCGCTCTGGAGAATAAATTTCAAGTGAAAGGAGAAGGATTAGTAAATGAAGAACAAGTTCTTGCTTCCATGC

CTCAAGCGGATCAAATCTTATCTCTGACCCGAAGATCGAATTTATCGTTGCTTTCGATGACTCCTCGGACTTCTCTTTTA

TCCTCACCATCCATGCCCGCCTCGCCACCAGTGTCAGCCCCACAGCAGTCTCTCACAGAAGATGAACAATTTTCTATTCC

ACAATACAGTCAGCACCAAAGTCATCTTGAGACGCACCTTCCCTCCCCTTCTCAGAGCTACACCAGAGCCCACAGCCCCG

AGTACTACCCAACGGAGCCACCCGCTCCTTCTCACTCACACGTTAATTTCTCCTCAGAATCTCAAGAGATGTTCGAATCC

ACCGCATTAAACCTTTCTTCAAGATCAAGTCGTTCCCCCAGCAGCATGGACTGCTGTTACGAGCAAGCGCGCTCCCCTGA

TATGGGTGGGTCATGCCTTCCCCACAAACTCAGGCACAAGACTCATCACACTCATGCTCTCTCCAACCAATTCAACAACG

TTGCTAATACGTCAACACGACCACGGTCGGCATCACCGGACCAGCTGCCCCCAACTGCACCTCACTTCTACCAACAATCG

CCTTCGTCAAGCATAGTCAACGTGTCACAGTCCATACCGAGGTCGTCTACTTCCCCCCAAACTTCTCACCTACTTTTCCC

TATTAAAAGTGAACCCCTTTCCAGAGAAACTGGAGAAGAATCGCCAGGATCTTCGGACGATAGGGACTCTGGCATCAGCT

TAACATCTTCACCTCCATTGCCAGGAGAGCAGAGCTACCCGTCGTCCAACCGAGAGTCAACCGAAGACATGGAGTGTGAC

AGCGAGCAGCAGCTGCGCGCCGAACTTCACAAGCTCGCGTCGGAGGTGCGGTCACTCAAGTCTTACTTGAGTCGAAATGC

CGACCCTCACCGGCACCAGCAGCAGGACACTCGATAGCTTCGTTACTCTAGACTTAACTTTATATTCTTGTTGCTCATCA

TAGGTGTGAAGTGTGAACTAGTAATTAGGTTTTGATGTTATCAAATGTCGCTCAAACATGCACGTCAGTCCAATTTGCTA

CTAACTTAGCAATGATGAGAGGACTGATGCTGTGGTATCTTTCATTTTTAGCAAGTCCCACCTATTGTAATCTGCTTAAG

GCAGTCGGTTCAGTGAACTAGTTCACTTTTTTGCTTCGTATTAACATTACTTTTGCCCATCACTATGTATTACTATGACT

ATGTAGAGATGTTGATCTCCTCTGTTGCAGCTCTCGCTCATATATAAAAGCGTAGTTGAAGTGACGTTAGAATTTTTAAG

CACTGCACTGTTTTAAAGCGATGGCAATGTGATATGAACGAAATTTCAACTCTTCCAGTGATGTCGCTCTCACCATTTCG

CCACGGTGAGGGGCGCCCTCCATCCTCGGCTGCGAGATCTAACAGTGATGCGCTCGCGTGCTCTAGTACTCAAAATCGCC

TCTCTGGTTGACAGGAAATTCTTGCAATTTATGATTTGCGGCGTCAGCATTTCTGGTGTTCACACTCTATTTCACGAGTG

GGGACCCTAACCAGAAATTAGGCCTATCGGAAGGTCGTAGTTGTCCTATGATCCTCTTGCCCCCTATGCCATAGCTCTGG

CATTCTGTGGTTTCCTGGCACCATGCATTCCTCTGGCACACAGCGGGCGTCTATTGACAACAGACACACGCGCTTATGCA

GTTCCCGCTTGGGGTTAACTTTACCTTACCATTAGTAGTCCGTTCTTTAACCTACGGATCCTCCAACTGACACTTTATGG

TGCCACTCGAGTAGTCTGCTCCTCAACCAACTGATCCTGCGACTAAGGCTTGAACATTTCAACTGTAAGCGCTGCAACAT

TAGTATTAGGCTTGTGCTACAGCTGCAGCAGCTGTAACTGAGCTGATGGTGTGCAGCAGCTGTCACAACAAAGATCTGGA

AGCTCAAGAGCGTAGCACTGTTGTGTAGGTCTACGAACTTCTCCTGCGCAATTGAATGAGTTGCACCAATGCATTACTAT

GGTAATATTTATTTTAGTCTTTGTGTTCCACTGTTTCGGCTGTAGGGCGCAACTTCCTTATGATCTAATTATTATTGTAT

TGTAGGCGTATGTGTGTGTGTGTGAACTTTATCTGTACTTATGTAAGCTTGCTTGTGTTGGGCCCAGACTCTGTACTACT

TGACAGTCCCCTTCAGATGTTGATGCCCAGACTGTGCCCTGCCTGACAGTCCCCTTCAGATGTCGGGGTCCAGACTGTGC

CCTGCTTAACAGTCCCCTCCAGATGTTAGGGTAAAGACTGTGCACTACTTGAGAGTTTACTGTAGTTAGGACTTGTTGCT

ATTGAGTTCTTTCTTTCTTTCATTTAGGTACACTGCGGTTACATCGTTTACGATGCCATACAAACAGCGAGCAAACGGGT

GCTGGTGCTGCATGCAGTGTACTGGTCCTGGTGCTGCATGCAATGTACTGGTCCTGTTGCGGCATGCAGCTTACTAATGA

GCAGCAACTGCCTTACGTTACAACAGGCAGGCCCTATTACGACACCGAATTGATATAAACTTGATCAAAATCTTCGTACA

TAATTTTTTTTTTCTTATGAACTATCATATAAATATAGACTTATCGAATAGGTATTCCAGCTCAACAACGTGGTACATAT

AACCTAAATCAAAGAACTAATTATCTTATAAACTTGATCGTTCAATATATACTTTTATTAAAATTTAATTTTTACATCTT

ATTATTATTATTTCTTAAATACAAAAGTCTAATTAACTTCTGAATGTCTCAAGAAAACTTTCTAACTTGCAAACTTGCAA

GCCGTTCCTTAACCAAGCCTCCAGGAAGCTGCATTACAGGGCCCAGCCTCGGATTATGCTAACTGCGACCCACCTGTCCC

ACTGTCCTCCTCTCACCCGTTACTGACTAACTTAGCACGGCAACTGATCGCGTGCCGACTCTGCGCATCTCCTTGCATCG

CGTCGCTTGCAGTGCAGTGACTGGGAGCTCGCGCCCTTAGTTATCCAGACCAATTACGAATATTTCTATTGAAATGAGGT

GCTTTTTGATGTTCAGATATTCTAGGAAAAATCTAGTTAAATTTTTTTTCTGAAATCATTAAAAGCCGTAGTTATACTGT

TCTCGTAGTTTTTTATGTTTTGCCAGTGAATGATATTTTCTACCGCATTTTTTCATTTGGATGATATGTTTATTCGAAAT

TGCCAACACTTTATACCGAGCGATAATAATTTTCCCTATTTGTTTGTGGTGAAAGCCATGCAGTTCTCAGCGTATTTAAA

ATGCAAGTGCAAAGAATGATCAAATTCATTGAGTTTACTGTCGATCCACGTGACGTCAGAGCTAGGGAAGATGGGGGCGC

TTACGATGCTGCCTGCGCTACTGTCGTTCCACGTGACGTCAGAGCTAGGGAAGATGGGGGGCGCTCACGATGCTGCCTGC

GCTACACTGAGGGAATTTTTGTCCCAAAACCGAAACACTCCGCTGCTGCTGCTGCTAGAAAGTGGTAGCAAACTCCCCTA

CTGCTGCAAGATAGTTGTAGCAAACACAGAGTGCGGAAGGAGAGAGGT

Protein phosphatase 1A

>lcl|PH.k31.comp909_seq2

TTTTGTCATTTTTATCAGTGGCGGGTAGTGTGTATGGTCTAATTTGTGCACGAATTAGGCGTTTCCTAAATATAATTTTG

ATTTTACATACGTGGTCGGTAGACCAATAATAAATATTATAATGGCTGAAGCAGATAAACTGAACATAGACAGTATAATA

GCAAGATTATTAGAAGTTCGAGGCTCACGCCCGGGCAAGAATGTCCAGCTCACAGAGAATGAAATCCGAGGCTTGTGCCT

CAAGAGCCGAGAAATTTTCCTGTCGCAGCCAATCCTGCTGGAGCTCGAGGCTCCTTTGAAAATTTGTGGTGATATTCACG

GCCAATACTACGATTTGCTGCGCCTCTTCGAGTACGGAGGATTCCCGCCAGAGAGCAACTATCTCTTCCTCGGTGACTAT

GTTGATCGTGGCAAGCAGTCGCTAGAGACCATCTGCCTCCTCCTCGCTTACAAGATTAAATATCCCGAAAATTTCTTCCT

CCTCCGCGGCAATCACGAGTGCGCCTCTATCAACAGGATATACGGCTTCTACGATGAATGCAAGCGCCGGTACAATATCA

AGCTATGGAAGACGTTCACCGACTGCTTCAACTGCCTGCCTGTTGCCGCCATTGTGGACGAGAAGATTTTCTGCTGCCAC

GGGGGCCTGAGCCCTGACCTGCAGAGCATGGAGCAGATCCGCAGGATTATGCGGCCCACCGACGTCCCCGACCAGGGCTT

GCTGTGTGATCTATTGTGGTCTGACCCCGATAAGGACACAATGGGCTGGGGTGAGAACGACCGCGGTGTATCCTTCACAT

TCGGCGCAGAGGTGGTGGCCAAGTTCCTACACAAGCACGACTTCGACTTGATCTGTCGTGCCCATCAGGTTGTGGAGGAC

GGCTATGAGTTCTTCGCCAAGCGGCAGCTGGTGACGCTGTTCTCCGCACCCAACTACTGCGGGGAGTTCGACAACGCCGG

CGCCATGATGAGCGTTGATGAGACACTCATGTGCTCCTTCCAAATACTTAAGCCTGCAGACAAGAAGAAATTCTCTTATG

TCAGTCTCAACTCAGGACGACCCGTGACGCCGCCACGGGGTGCAGCCAATCAGAAGCCTAAGAAGAAGTAAATAGGTGGA

GCTCCTCGCTCATTGGCTCTCGCCAGTGACCAATGACGACCGAGATCCGCCGTCATTGGGTGTAACTTATGCAAACGGGG

TGCAAAATGCTCGCTATTGGTCGATGCGAAGCCCATGGGGAATAGGCAATTGTGATTTGTTCATGCTTAGCCAATGGGGA

TATGGGTGCGTGTGATTGGTCGTTGCTTAGCCAATGGGGCATAGGCATTTGTGATTCGTTGATGCTTAGCCAATGGGAAA

ATGGTGATGCATGATTGTTTGATGCTTGGCCAATGGTGAAATAACCGCGCATCTTTGATGTGCCTCTCATTTGTAGGTGA

AAGTGCTAAAATTAGTCCTAGTACCAGCATCAACATGCCTGGCTTACGGTATACTGAAGTGAATGTTTGTCGGCAATTTG

TGTTTTTCTGAACTGTCATTGGTTAATGTATCTACCAATTTTAATTCTTAGCGCTGTCATTAGTTGCTCTATTAAATAAT

GGAATCATTCTGAAGCAACGTACGTATTACTTGTAAGTGTCTGTGGGCTATATTTGATCATTACGTGAGACCCGAAAAGC

TTGCCATTGACTGTGCTTTGAGCCACTAGCGTGTTTTTGGTATCACATGGAGAGGGGGGCACTGAGGCTCCCCATTGGCT

GAAGTTCCGACGAAAAGCAATGCGCGACCTTGTGCTGGTCGGTCTTTGGTTCGCCTCCGGTCTGCAACGTTTGCCATTGG

CTACTGCCTCGACCAATAGCAAGGTGTGACTGTGTTTTGACGATAGCGTACTCAATGACCTGTAAATGACCGACATGTGT

TGCATAGGGAATCAAACAACCAATGTCTTCATGTATAATTCACTACGTTACTCTTCTGGGATAATGACCTTAGCTACACT

GTTGAATTTGATCCTTTAAAATATCAGATTTACATACACCATTAGATGCTTTCGGTCATTTCTACGTTATAGATATGTAT

AGCAACTGCATTCAGGTGAAAACGATGTGCAGTCAATTGAACCTGACGTGCGTTCAAATGAACACGACGTACATTCGAAA

GAACCCGTCGTGCGTTCAGATGAACATGATGTGCATTCAAATGGCTATGTACATTCATCTACACGCAATGCGCCCCAATA

TTTACACAGTGCACGTTCATATTCACCAATGCGCATTCGGATTTAATTAAACAATGCACTTACAGATCAACTTTGGTATC

AATTCAGATTAACAAATTCTTGCTCTTTGTTTCCACGCACTGTAGATACGTGACAACTGCGTACCTGTAGACACCATTTT

CAACTCTTCTATCTCCATAAATATATAGTAGATTTTCCTCCATAGTTCACAGACTGTACATGATTATTTTCTAGATGACG

ACTTGTGTTTTCCCGCGCAATACATCATCACGATTTCTTTTTTATGAACTTGAACGTTGATTTCCCCTTTTTCAACAGTA

GTTGTTACTCGCATTACTGTGTATTACTATTATTTCCTCCATACGACAAGACCTGGTAATTAGTTAGTAGCTAATTACTA

ATCTTGAATTATCTGATGCATGATTATTCTTTGGAAAACACGGCCGTTCATTGTTTATTCATCATGGTAGTTCATTATTT

ATTAATCTACTGAATACAATGATTTTCCTTAAAAAGTCGTTGACAGTTGTTTATTCTGAATCAGTCAATACAAGAATGTT

TCCTCCAAGAAACCTCGGAATTTGCTGATATTAAGTTAATGGTTCATTGTTGGCTATTTTATAGGCCGTCGCTATTTCTA

CTTTCGATAAATTAAGACAGTTTTCTCCAAGAGAAAACGTTTCTTGATTCCTAAATAAAGATATATAATAATTTAATCCC

ATCGTAGTCCGTTTTGTGGTGGGGAACCTGCTAAAGCGTGATTGGTTCTTAGGTCCCTGCCCATCAGAAGCTGGAAACGT

TGTGTGGTGGGGAACCTGCCAAAGCGTGATTGGTTCTTAGGTCCCAGCCAATCAGAAGCTGGAAACGCGTGTCTGTTTGT

GACGCAAGCATAGCTTTCTTTTGATCACGATAATGCGCTCATCTCTTGTCAATGAAGTTGTATTTATCCAATTATTATTG

TATGGGTATGTATTTATACACTACTGTGCTTTACTTGACACTCTCACTAATAGCATTTAGCCTACAAAAGAGTGGTGTTG

GTGGTGTCATGTTCGCTTTGTGTGGACAAATATAGCGCGTTCCCCCCTACGCTTGTCACTGATGCGCTAGGACTAGTAAA

CACGTATGTAGCTAAGGACCTTATCCCTCGAGGGTGTCGATACACAAGGTTTCCTGTAATCTTGCTATTGTATGTAGAGT

ACTGTAATATGTTACTACTATTATTATTATCATCAGGGATGGGGCGGGTTTTTTTGGTAGCAGTGTCTGTTTCCAGCTTT

TTCCTCCCTGCAGCTTTTTTTTTTTTTGGAAATTTGACAACCTTTCTAAGCCAAAATTTTCAACCGTGATCATGCACCTT

TTTTGTTAGAAAAGATTTTGCAACTCCTAAGGAAAATAGGTACCTATGCGGAAGGCAGTCTCCTGAAAACGATATTAAAT

AATTAGTGTAACAAAACGAAGGGACAGTGACCCTGGAGCCTATGTATGAATGGATCTCAGACCCTAATGCCTATATGACT

GAAAGGGCGGTTGACCTAAAGCCTATTAATTTATAACTGACAGGGCCGCTAACCCTAACGCCTATATAACTGATAGGGCC

ACTGACCCTAACGCCTATATACTGATAGGGCCACTGACCCTAAAGCTATATAACTGATGAAGGCGGTTGACCCTGGAGCC

TAGAAACTGAAAGGGCCGTTGACCTTGGAGCTTATATAACCGGAAAGGCTGCTAATCGTAATCCCTACTACGAATACGTC

GGGAGCCCTGGCCGCAGAGTTCCCAAAGAAGGTCGCAGGGCGAAGATAGCTAGAATCAAGACGTTAACTGCCATAGAGCC

GCACCCAACCTCCCATTTTTATTATTATTGATAATATAAGAGTAATGGCTGCGCTAGGGTCGTGGAGCTTTAGTTAAGCA

GTTTATTGTTGGCAGGGCGTGGTTCGTTTCCACTTCAGTGCTTGTACTCCTACTGATGAACCTTTTTTGCTGGTCTTCCT

CTTCTTTCTGGGAGTACCGGCTCACCTAAACTGTGTAATAGGCTATTGGGCCCCCAGGAGGTACACATTAGGTTGCCTGT

AGCCTCATTTTGGTCTTGGGCTAGCTTTATTTTGTGCCTACGTTGCTCCTTTCCCCTTTACTCCCTAGTATTTTTTGCTG

TATTGTTTCATCTTCTTCTTTCTGATGAAGTGATATTCCCGCTGCTCCTTTCCCATAATGATCTTTTTGCTATGGTCTCA

TTTTTTATATACTAGCTACATCCTGTGTTTCCAGACGCTAGTTTCTCCTGTACACTTGCCCCCCCATTTTCCCTTACGGT

CTTGACATGTTCCGGTAGCGCTAAAATCATGATAATATATAACCTTGGGCGCGATGCGGCGTGCAGTCATTCCTTTCTCT

TTCTGTATACCATACTGAGCTTCTTGGTCTGTCCTATCCTACGTCCCTTCTCACTCCGTCTAATCAACGAACGACCAACC

GACCGGAAAATTTTACGAACCAACTCCTGCCAGCAATAAGCTGCATTAGCGTGACATCTTCCACGACAAGTATCGTTATG

ATCTTCATGCGAACACGACGGCAAGCCGCCGAACTATGTATCTACCTTATAGAAGGCCAACTATGTACCTACCTTGTGCC

GGCCGGCTCTTTGCTTCTAAATTTGAAAATTTTCGCCAGTTCCACACCTGAGTATTTGCCATTCTATTATTAACCAGACA

ATGGCTGCTGATGTTAACGGGACATTGCTGTTGTTTGTAACGATCTTGGTAGGTCGTGAAATGCTTTCAAGTATTCTGCT

AAGGGACTTCTGTGTACTTTGTAGGCATCTTCCCGGTGAGATCCAGCGACGTCTTTCCTTGCAGGTTTATTGACTTGTGT

AGGTAGGGCAGAGAGGGTCATCTAACTTTTCATGCCCATATGGCCTGCTTTCATTTTTTCCTACGTCTAGTAATGGTTAT

TGCGAATTTCCTTTTTTTTCATGGCTTGCTGTTCCTTTACGTACCACTCTGTAGTATGTGGGTTCTTTTTCATTTGTTCA

AACTGTGTGTCCCTTCTCTTGCTTCCAAAACGGTGTCATGTTTAACGATACCGAAGCATGCATTCGATTTTGCCCCTCGT

TTACTTTCTTTCATTGCCATGCTGTTACTGAGACACTAACAGTCTTTCTTCTCGTCTACCTCTTCTTTTCCATTCTCTTC

ATACTTCTGTTTGTGTTTTTTCTTCTTCCTCTCCTTTGTTTCTTGTGCTCATTTCCATCTTGTTTCCTACATATACTGAT

AACTTTTTGTAGTGCTCTTCCTGTGCTTTTCTGGCGCTGTGGCCTTCCAACATGGGTTCCTTTTTCGTTCGATATGCGTC

TCCCTTGTTCCTCATCTGCTTGGATCACCATTATTGTTGATGGGATCCCCAACCTTCCTAATGACCCTATTAACACATGC

CCGTTACTTTTGCTGAAAGGGTTCTGTCCTATTGTAATCCTACAGCAAATTTAACTGTCCTAAATCCTATTCATACCAGG

CTGTTATTTTGCTACGAGGGCTCGATGTCCTATGGTATTCCTAAGGGCACAAAGGCTTCTTTCGTGAGAAGGCTCATCCT

GTTTCCTGATAGTAATAGGGAAAACGTATCACCCTTTCCTCTCTATATTCATACTTATCTCTTGGTGTTTGGTCCTGTCC

CTACTTACTGTTCCCTCTGGCGCTGATCCTGTGTGAGAACACCCTGTTGCACGGGTTCCTCGTAACTGTCAACACGCTAT

GTGTCCCCTATTCCTTCCCCTGTTAGACCGCCTTTGCAGTGGAGTCGGAGGCCCCAACTATTCAATATAAATAAGTGAAT

AAGTAAAGAACGCGAGGTCGACCCTTGGTGTAATAAACTGTATCAGGTCTTGCTGGCTTGGCTGCGCCCTGAGCTCCGGG

ACTAACTTGAGACGAGTTTTTTTTATGTTTAGCAGGCTGGTATATAAATATAGATGTAGTCTCCTTTTTGCCTGTGTATT

GCTTCTTTTCTGATCTTCAGACCCTGCCAGAGTTGTGGAGTGTGACCGTATGCCTATCGAGATCTGGAATAGTTGGGCCG

CTGGTTATTGAGTGTGGCCGTACGGATCTCCAGATCTGAAACAATCTGGACGCTCTTGTGACTAGACAAGGTTGTCTTGT

ATGGCTGTATGCTTGTCGAGTTCTGGAGCGTTAGGACCACGGTTTACAGAGTGTGGCCATATTCTTGCCTAGATTTGAAG

GTATTGGGCCACACTCGGTAATTATAGACACGTTTGCAAGGTGTGGCGAGATGATTGCCAAGACCAAGATCAGAAAATTG

ACGCTGCACTCGAAGACTAGGACCTCTGCTCGCAGGTCCCGCACTGTCACGTGTGGCTGTATATTTGTCGAGATCTTGAC

CAATAAGGCCACTACTATATATGATTAGGACTCCTACTGATGGTGTTTTTGCTGCCTTTCCTCCTCATTTTTCCTCTATT

TTCTCTTCTTAGTTGCGGTAGTCTTCCGTACTGTGGCCCATTTTGGTAAGCTATAAGAAAAACCCAGCTTATTTTGATGA

GCCATCTAGAGAGGCGTGGCGTGGTACAGTAACACCGCGGAGCTGTAACGCTCACTCAAGTAAATTGTTGGGATCACTCA

AGTAAATTGTTGGGAAAAATCACCCATTTTGATAGGTTTTTTGGAGGGGAGTCTCGAGTTTGTGGAGAGTCAACGGCAAT

TTCAATAGGGTATTTCGAGAGAAAGAAGAACTTCTGTTTTGACGAGGTGTCAGAAGAACAATGACCGCGTGTTGAGCCTT

AGGCAGAAAGAAACAGCCATTTGAATAAGCTGTCTAAAAATTGTTTCGAAGAACTGATCGAGAAATTCTAAGTTTCTAAA

GTTCCCTGCATTGTAGCGAACAGCGAGAAATTGTTTCGGCGAACTGATCGAGAAATTCTAAAGTTCCCTGCATTGTAGCA

AAAAGCCAGGCTATGTGGCTGACAGTACGGTTAATTTGAACATCGCTGTGTGTTTAATCAACGTCGGAGTGTTGTGTGAC

GTTGTGCTACGATGCTGGCTGTGGGTTGCCGCTAGGACTGCTTTTGTTGCAACAATTTTGAAGGGAGTTGTTGTTCATTT

CAGTGTAGAATGAATTTATTTTGGGCGCTTGTAAATGTGGCTCGCTCTGTGTTGTGTAAACTTTTTTTTAAAGAAAAAAC

TTGCCTATCACTACCAAAAAAAAAAAAAA

Microtubule star (PP2A)

>lcl|PH.k21.comp640_seq0

TGTCACGCTATGAGCGGCCATGTTGCACTAGAGGGGAGAAGGAGACGGTGTGAAGTGTGTAAGTGGGGTTCTAACTTGAA

CATATTTTCACATTGTAATTGACTAGCATAAGCATGGAAGAAAAAACGCAAATCAAAGAATTGGATCAATGGATCGATCA

GTTAATGGAATGTAAGCAATTAGGGGAAAACCAAGTGAAAACATTGTGTGAAAAGGCAAAGGAAGTTTTAGCAAAAGAAG

GCAATGTGCAAGAAGTCAAGAGTCCTGTCACAGTATGTGGTGATGTCCATGGACAGTTTCACGATCTCATGGAGCTGTTC

AAAATAGGTGGCCGTTCTCCTGATACCAACTATCTGTTCATGGGAGACTATGTTGACAGAGGCTATTACTCTGTTGAAAC

TGTTACCCTTCTAGTCACTCTCAAGGTTCGCTTCCGTGAAAGAATAACAATCCTACGTGGCAACCATGAGTCTCGACAAA

TTACTCAGGTGTATGGCTTCTATGATGAATGTCTGCGGAAATATGGAAATGCCAATGTATGGAAATATTTCACCGATCTC

TTCGATTATCTACCATTGACTGCACTTGTAGATGGACAAATATTCTGTCTTCATGGTGGATTATCTCCTTCCATTGATAC

CCTAGACCACATCAGGGCACTTGACAGGCTGCAGGAGGTGCCACATGAGGGCCCTATGTGTGACCTTCTGTGGTCGGATC

CTGACGACCGTGGTGGCTGGGGCATCTCCCCGCGAGGCGCTGGCTATACCTTTGGTCAGGACATCAGTGAAACATTCAAT

CACTCAAACGGCCTCACATTGGTGTCGCGGGCTCACCAACTCGTTATGGAAGGTTACAACTGGTGTCACGAGCGTAACGT

TGTCACCATCTTCTCCGCACCCAACTACTGCTACAGATGCGGTAACCAAGCTGCCATAATGGAACTGGACGACTCCCTCA

AATATTCCTTCCTACAGTTCGATCCGGCGCCGAGGAGAGGCGAGCCTCACGTGACGCGCCGCACACCAGACTACTTCCTG

TAATTGTGCGGCGGTGTGCCGAGTGCTGCCTGCGGCTCTCCCTGTGTGTGGCCGCTGCTGGTGTCTCATGTGTCACTTCT

GTACCATTGTACTGCCGTGTTGTTGGTGTCGACTGCTGCCTGCACGCCTATTGGTTCTTATGGAGAGTAAGGGGAGGGGA

CGCCTGTGCATTGTGTTGAGTCTTTAACTTTGTATTAAATGTGAGAGGCGTGTGTGTCTCTTCAGGGCTGTTCTTGACAG

GGCACAGATCTGGCCCGTCCTCTTCGGCGTATGCGATGCACAGCAGATGGAGTTATGCCGAGAGCTGTGGAGGCAGGTCT

GTGGCTGATCTATAAACTTATCGAGGGTATCGTATGATCCATTTGAAGTTGATTATCATTACATGGAGTTGCTGTGGAAA

TTTGTGTCTAGTTGCGTTAGCCAAAATATGTTAGGGTCGCGTTCTTCTATTATTCCACTTGTGTGATTAGCTCCATAGTT

GCGTGCTTGTCACTTCCCGCAGGATTTTAGTTACTTGTGCTTACTAGTTATTTTTTGATTTCTGCATTTTCTTTGTACTT

TCTATTATTTTTTGCCTTGCAGATTCTTACTGGTTCACAACTGTGCATTTTCTTACCTAAAGTCTAAATATGGTTGCTCC

TGGAGTGTACTGTGCCAGAGTGCACGTGTAATGTGTTTGTTGCTCTTGTTTGATTGTACCGCGTTGCCGGGGCGGTGCTG

GTCACGGGTCCTGACCCAGGGCCGCCTTGCTGCAACTCTGCGCTGTCTGCCTCTTCGTCGTCGTCTCAGCAGCTGCTGCT

GTGCGGCCCGGAGTTGGGCTGTGCTGCGCCTGCTGCTGTGCGTCCCTTGCTGCACGCTGCCTCTGCTGTGCTGTGTGGAG

GTGTTCCGTTCTAACTGGTGCTGCACCTCCCCGTTATTGGGCTGTTAACTTGTAAGTTCTGTAGTGTATATGTTACTCCT

CCCCAATGTTATTGTATTAATATATGCCCTCTTTACTCCAATTCCTGTAAGGAAAGGCTGGTCGTACCTTCTCCTCTATC

CTTTGGTTCATTGCTGTCGACGTTTATTCTAATTAATTTAATTTCTTCCCAAATGTATTGTAATATAGACCTGTAGTGTT

CTCGTTCATCTGATTTTAGTTCAAATAAAGTGATCCCCGATATTACGTAACAATTTTCTTTTGTGATTCGTTGCTTCTAA

TTCATCTTCGATGACTGTGACTCTGGTGCCGTATGGATAGAACTTCTGTTCTTGCTGGGTGTGTGTGCATGTCTATTTAC

ATTGTGCATTGAAGAACTTGAGCGCCTCCGAGAAGTAGCTCAGCTGATTAGTCCCGCGTTAATTTTATTGTCCCAACCTT

CGGCCGAGCGCCTTGAGTCGGTTATTCTCTATATAATGTACAGAAATTGGTGTTGAAATTATCCACAACAGTATAGATGG

CTCTCAAAATTTCTTCTAGCCAGCTTCATGGCACTAGAGTCTATTGTAATTTGTAGGTAATTGTGCGACTTGTTGGTGTT

GTTAACAGCTATGAGAAGGAGCGGTGGTCTTCGTTTTTGGTCTGGAGACTATCGCGTGTTGCACGGCTGCGTATTGTACG

AGACCGCTTATACCTAAATGCGCCTGTCCGCATTAAATAAATGTCTATTGTTTATATGCTTGGACATCAGAATACGTATG

TATATTATTGTCTTATCCTGGTTTTATGCCTAGTGTGCAAGTTAGTAGGGTTTGTGAAAGTGAAAGACTACGGAATTTTA

AAATGCAATTAAATTGTTGCACAACGTTGCGTATGCATCATTGTACTGATAGTTTGCCATGGAATATTTAGGAATTACTA

TTTGTTTTGAACCGATGGAAGTAATCTTATGAGTTTAAGTTGTATTGGAATACTGGTCTGAGGCTGCCTTACAGTGTCAT

CACGGCTGCTGAAAACTGTTACATTCCCGAAATGTGCGAATGTCAAAGTTTCGCCACCTCGTGTAGGTAGTGTTTCCGCT

GTATGGAGTGTTCCAGTCGCTTCTTAGATTGACTCGGTCTACTGGTACGTGAGCCTATCTACCCTACTTAGAGTATCAGC

GGAGAGTAGGTAGCTACAACAACTAGCTAACGGGGAGTTCGCCTGTGTGAGCTGACGTGGGTGGGCTCTAAGTTATTGTA

TTTTCAGTCGCAAATGGAGTAGGGGTCTGCCTGCCGACTATTGTAATGACCATGCTTCGTTCACTGGATGTGCTCCCGGT

GTATGTCAGCGATTGAAACCTCTCAAGGAGCTCAGCAGCATGTTTTCGAGCAAGGTCTAGGAACTTTGCGTTGTGAATGG

TTCGTGAGGTTGAAAAGCTTATTTCTCCTATATTGTAGAATGAATGATGGGCTTAACGCGGTAAAGAAAAACACTACTGC

AATTAATCTTTTTTCGAGTAAATCCTGTATTCAAAAAAAAAAAAAAAAAAAAAAAAAAAAAAAAAAAAAAAAAACTTGGA

AAAGTTTTTTATTTTTTT

Widerborst (PP2A)

>lcl|PH.k29.comp1930_seq3

TGTTGAAATCGCGATTGTGTGTTGTGTTAGTCTGTGAATATTAATCAAACTAATGCCGGCAAGCAGTCATCAATAAGTGC

ACTGTGTTGTAGATATAAAGGACTATATGTATATATTTTTATAAATGACCAAACCTGTTTTATTATTGAGTAATATAGCC

GAGTGTTCCGGTAACTTCGCTCCGTAGCAGCACCGTCACCACCGCCTTATTTACTTCTGGAAATAAAAATTGCAGTTGAC

TTGCACTCCGGCTGTTAACGATTGATACGGAGTCTGTTACTTACTTGGACATTATTGAGGTGGTTCGACATTACTAAGTT

TGTGGAGATACGCCCTATTGCCCGCGCTCTGCTTCCTAAATCCATTGTGCCACAATATTTGGATAAGGCTAGCTCAACTT

CGGCAAACCAATATTCTTGTAAGCTATAATAGACCGTAATTGCAGGCGGTTTCCATGTTTTGGATTCCTTTTTCATTATA

AAGTGCGTAAGTCGATTCCATATTTTTCATTTGATCAGTCAATCTTCACTTGGTCGAGATCATCATCCACCTGATAAACA

TTCAAATAAATATTGTGAAACTGATCTACCATTTGTGAATTGCAGTGACTGGTGATGGTAGTAGAATAAGTAGGCGGTGG

TGGCAGAGATGATGTGGGTTGACGGCTAGGCTGGTGCAACACAATTAAAAGAAACAACTTGAGCACTGTGGCACAGGTGC

CACATGCTGTAGGTAGCTATAATTTACTAAGAAGAAATACTAGACAACATGTCTGCTGGAAGTGGCAACTTTGTTGATAG

AATTGATCCGTTTGCTAAGCGGTCTCTGAAAAAGAAACCAAAACGATCTCAAGGCTCCTCTCGATACCGGACTGCAAATG

ATGTGGAATTGTCACCCCTGCCTTTGTTGAAAGATGTGCCTGGATCGGAGCAAGAAGATCTGTTCTTACGCAAGCTGCGG

CAATGTTGTGTTGGGTTTGATTTCCTGGACCCTGTGGCAGACCTCAAGGGAAAGGAGACCAAGCGCACCACGCTCAATGA

GCTTGTTGACTACATCACGGCTGGCCGTGGTGTCCTCACCGAGCCTGTATATCCTGAGATCATTTCCATGATTGCGTGTA

ACCTGTTCCGCACATTGCCTCCATCGGACAATCCTGACTTTGACCCTGAGGAGGATGATCCCACTCTTGAGGCATCGTGG

CCTCACCTCCAGCTGGTCTATGAATTCTTCTTGCGCTTTCTGGAAAGTCCAGACTTCCAGCCAGCCATTGGAAAGAAAGT

CATCGACCAAAAATTTGTCTTGCAGCTGCTGGAGTTGTTTGACAGCGAGGACCCTCGAGAGCGGGACTTCCTGAAGACAG

TGCTGCACAGGATATATGGCAAATTCCTCGGCCTCAGAGCATTCATCAGAAAACAGATCAACAATATATTTCTTAGGTTT

ATATATGAAACGGAACACTTCAATGGCGTCGGAGAATTACTTGAAATTCTTGGCAGCATCATCAACGGGTTTGCTCTACC

ACTGAAGGCGGAGCACAAGCAGTTCCTGATTAAAGTGCTGATTCCGCTGCACAAAGCAAAATGCCTGAGCCTCTACCACG

CGCAGTTGGCGTACTGCGTGGTGCAGTTTCTAGAGAAGGACCCTACGCTAACCGAGCCCGTAATAAAGGGGCTTCTTAAA

ATATGGCCAAAGACGTGCAGTCAGAAAGAGGTTATGTTTCTAGGTGAAATAGAAGAAATCTTGGATGTAATCGAGCCTAA

TCAGTTTGTGAAGGTACAGGAGCCTCTTTTTAAGCAGATAGCAAAATGCGTCTCTAGCCCCCATTTCCAGGTGGCTGAGC

GAGCCCTCTACTTCTGGAACAATGAATATATTATGAGTCTTATAGAAGAGAATTCCAATGTCATCTTACCCATCATGTTT

CCGGCGCTCTACAGAATTAGCAAGGAACATTGGAACCAGACGATCGTTGCACTTGTCTACAATGTTCTCAAAACCTTTAT

GGAGATGAACAGCAAACTTTTCGACGAACTCACTGCATCTTACAAGAGCGAGCGGCAGAGGGAGAAAAAGCGCGATCGCG

AAAGGGAGGAGCTATGGAAGAAACTGCAGGCCCTCGAACTGAACCGCACCCAACCTAAAGAGTAGTTACCTCCATGAGTG

TTTGTGTGCCAAACTCAGAGAACTCCCTTCTGATGCACTCATTTATCCTCCCCTCAACGTCGCCCTTCATTGCTTCTCTA

TCACTGCTATTGCTCATGCCCTGCCTGGAATCAGTCCCCTGCACACTCTTTAGTTTCTCCTAAACATTCTCTCATTTCCT

GTGTAATCATCATCGCCGCCCCCGCCATTGTTTTAGTTGTTTTTTCTTGTGTCTTTACTTCGTCGCGCAGCCGGTACGTA

TGTTGTGTTCTTGTCTCTTCTGTCCTGATCTTTCGAAGCTGAAATTCGCTAGCCGCGCTGAAAGTTTCAGTAGCGAGATG

ACTCTTGTAATGCTGTGCTACTTTGCCCAGCTTCACTTTTCACTTCGAGTTGCTGTTTCAGCCTCAATTTTCCTTCAATT

ATCTTTTTGCTTCCTTCGTCGATTTCTTTTAGTTGTACTTTTTTGTCAGTGGGTAGCCAGTATATCATTCGGATTTCCTT

CGAATCCTCATCATCAGTCATGGCACTCATCCCAGCACTTTATGCTACCCCTGGTGCTGCTGCACTTCCCATGTCAATCT

ATTTGCACCAAATTGGCGTATCTCCACCCAATTTTCTACTGTAATTTACTGTTGTGTGTCGAGCGATTGCAAGTCTGACA

TTGCTTTACCCCCCCCCCCCGTCCACTCCCATCAGAACCATAAGTCCGTTGTCGTCCCAGCCACTGATCCTTGCGGTTGA

CCATGACAGGACTTTACAATATTGAATGTTTGTGGTTACCGAGTTTTTTCTTCGATTTCAACTCCTAGCAACTGCCTCTG

CTGCCGTGATCCAGTGACATCCTTACGTACTCCCTAATCGCTTTTATGAGGTGCAAAAGTTCGGAGTGACTCTATTGACA

GTGCAAAGTGTTGCCAAGTGCGGACCTTATTTTCCTTGGAATGCATTTTCATTGCTTGGTTCTACTTAGAGCGTCTTACG

GGAATGTCTCATTAGATCAGTACTACAACAGACTTGTGTATTATTTAGAGCCCTTCATTTAATAGTGCAGCAAGTTTCGT

CTGGTAGACAGTGACTAGCTGCGGCCCTCCCGTACCGCGGTAATGTACGACAACTGTTTGGCAAATAGTGAGGTATCTCT

TTTTACCACAGAGTCATATAGCGAACTCTCAGTACAGTTTCCTTCTTTTCGATGCACTTCCACGACATTGCCGCTGGAGT

TCGATTGATTGGTGAGATTTGGTCTTTGGATATATACCTACCTATGACACAGTTGTCTCTTTGTGTACATCGTTTTTCTT

TTTACGTACCTACGATTTGCGTGCTTGCTAACTTTCAATAAGATGTGTGACACTTGTAGCGTGTTGTGTTCAGTAAACTC

TTGATAGCCTCTGCTGAGAGAACGCTTTGTCGTGAGGAGGGCTCATCCATTCCGCAGAACTTGCTCTGTGTACAGTCTGG

AGGAGGATGGCGAGTGTTAAACCGTAAAGTTTGTCTCACGTATGTAGCACCGCGTATATCAAAACAATGTTTTCTGTGGC

GCATCTCTTCTAATGGTAGGCTTCTCTTACCCTTGAGTCACCAGTTCCACCATCATTTAGAGGCTGCAGTAAGCTAGAGC

GTACGTCTTGTACGTTAGTAAGTTCGTAGTAAATTAATTTGTCGTTGTTTCGTACTTCACAAACTTATTTGAACGTTACC

TATTTACTCAATGATCAGAAGGGAATAATAGACTGTAATTTAAATATTCACCCTTTGACAGCGGAGTTGCTAATGAGATC

ATTTAAAGCTTCTGAAAGGAACGGTAGCACCCGCTTGTATAAAAATAGAAAGCTCATTTACAAGTCAAATTTCTGTACGT

TTGTAAACTTTATGCCTTCATGTGTATTGTAAAACATTTTCATGGGATATTAAGTTAGTATAGTTGACCAGGTACCCTGC

TCCTCCCTGTCAGTAGAGACCTCTCACCGTGAAATGGAAACTGGTAAGAACATGTGTGTGTGCACTACGAAAATCAGCGT

TTTCTTGCTGTCTGCTGTGGAACCGCCATACATTTTTTTTCTATAACTGCTTACCGTATCATTCGGTGCACAGTGAATAG

TTTCCTCTGATAAATAGGTTATTTGATGATTGGCTACTTGATGTTCACTTCTATGTGAGAAGTCCTCCGTTACGCCTCTT

TCATGTTGATGTTTCTCACGGTCAGAGGCTGGTATGTAGTTTTCTCTCGTGTTTCAGTGGGGTGGTTACGATGTTACCCT

CCGCTGCAACGCAGGAAACAAAGCCCGTCACTGCACAATTAAGCAAGGAACACATCGGCAGAGCGCTCTAAGTTGTCTAC

GCAATTGCTTCTGTCGAGTAAAATGTCGAGGTCTTGCTTATAACACTGCCAAGTCGCAATCAACATCTGTATATTTATTG

GAAAAAGACTTGTCTTGATTATATGGTACAATACTGCAGTGCATTAGCGGTCTGATCTCCACATTTTCTGACTCTAAATA

TGATGATTTAAAACCTATGACTGTCTTCGTATAAATTTATTATTGCATCGTGATTGTGATTTTGAATCTATTAACAGTGT

ATGGGCGTTGGTATAACTGGATAATTTCGAAAGTGTAGCTTTAGGTTCTCTCTTCGATTGTCCAATCGCCTGGGGCTTCA

CTAGGTATGTTGAATCGGCTGTTCGTCTGTTTTCTACATGACAGTTCATCTTTCGAACTTAGTTCCTCATTTTCTAAAAA

TTTCTATGACCACCACATTTAGATACTTTTATCTGCGTTACGAAATCGATTCCGTCATGGTTTTTCTATCACCAAGTCGA

GCACAGAATTCATCTGATGCGAACACATGAATGCAAATTCTATGGAAGCAAAATGTTCCCCGTTTCTCCCCATGATTCTG

GAATAATTCAGTAAGTAAAAGTTAGTCTTACTAGGCACGGTTTTATGAAAAACCTGTCGTAATGTAATCTAGTGTAATGT

AACAATGATATTCAGTTATCTGAGTTTTACGATTCCGCGATTATGACATTATAAGGTGGAATGCTTGGTTAATAATAGAC

CTGTTCGTATCGCCTTCGTTTTTCTATGAGCCTTCCAGTTTCTGTGCGACTAACTTACGAAATATTCGAAAAGAAAGGCA

GGCTCTGGGATCTTTGTTACCTCGACTGTAACTTGAAGCTAATACTCGTCAGGGCATCTGTATGCAGCCTGGCGCTCTGA

CTTAGAAACTAGTTATTTTCCCCACAGACTGAGCAGTGTGTTTTCTCTACTCCAGCGTAGCACGCGTTCCCGTGCATAGA

CGGGAAAGACCTATCGTCGCGGCTTCGGCTACTGCAAGTTTTTCCGTTGGTCGGTGCACGTGAAGACGTGCTGGTTGCGG

AGCAACACGTATCTCAGTGTTCAAGAGTTCCTTCCGCACACAAAATAGCCGTAAATTCACCATGTTCGCTCGTTTGCTTA

GGTAATTTTTTAATCCAACCGCTGCTATTGCGTCACAAAATTGCACTAGGCGACGAATAACGTCACTAGCCTTACTAGAA

AGACTTTTTTTTAAGAACCTGTCTGCCTTTATAGATCGCCATGCTTTGTTTCTTGCTTTCTTTTAGCATTCACCTACTTT

ATGAATGTGACTTCTTTACAAGTCGACTGGAGTACACTGGATACCGCGTGCTCTGGTGCTTCGGTTGTCATTGTTATTGC

ATAAAAAATAGCAGCGCGGGTACTACGTACGTAGGTACGTACCCAGCAGGATTTAATTAGCTACCTAGCATGTCTTTTCA

TCGGTTTGGCCCCTGTAGAAGTCTACGTGCATTTCAGAATTAGTGCTGCTTTTGAAACTTTCCCGATTCGGCGCGTTATG

TTGACACGTATTCCGTTTGCTTTGTGCATTTGGGAACACGTGAGTTTGTTTTACCGTAGTTAGTAATGTGTTATGAAGCG

GAGGACTGCATTGTTGCCCTGACGAGTACTGCGAGATACTAGTTTCTCCTCTTTCATGAGACAAACAGAGCTCGTTCTCG

CTGCACGTGTGACATAATCACGCATTTTTGGCCTCTTTTTCTAGATATTTCCTACCTATAAATTTAGTTAAACCTTCGTG

ATTGTGTAGTCACTTAATGATACTTTAGTGAAACTTGTACATTTGGCTGCATTGTTCGTTGTGACTCCAGAGTTGTTTTC

GTTGTTCATTTTGGCTCGTCCGATTTGTTGATCGAATTGATTTGTATCGACGGTATTTCCTTGAGCCACAAGATTTCGCG

TCACGAACTGTTCAATAGTCTGTGAGAGACTGTCAGTATGACGACGCATCTAATTTCTCTTCGAGCGATTAATTAATTTC

CTATATACTTGCTTAACTTACGCTGCGACGTATTGTTCTTTTTATCTTTTCGTCTTTTTTATATGCGCCCGAAAGACATT

TATTTCGGATTGATGTTTTTGTACAGATATAATTTGCAGTCCCTGTATTTTAATTGAGTGTACCATAGATAATATCAATT

TGATACCTCCATTTTTTTTATCTTGTGGTTGCGATGAAGCATTTTCTTACTAGCTGAGTTGGTAATGGTAGGGGCGGTTT

GTGAGAAGTGGTTGCTTCTCAAAATGACCCTCATCATCACAATGCCAGCTTGTGTTCGCTAAACCAATTTGTCTTGCGGC

AACTTCTTTTGTTTGTAAGATTTGCTATTGACAGCGTGTTGAATTGTTTCTGTATGAACTTCACTTCATTAACAAGTTGA

GTAAGAATTCTTATCTATGCTGAAGTGATGATGACTCTCATTCACAGCATCGCCGAATTTATTCAAGGCCAGGCAATCGC

ATCAGTTTGCAAGTTTTATCCTGCTCGCTTAGAACGCTGTAATTGCTTTAACGCGCGTTTTTTAAGCGAAAATTAGGGCC

AGGTGCGGGAGTCAGTAACATTTGAACAGCAACTCTGTAATCAGTAACTCCTAGTTTGAGCTCGTGCTACATCTGCGCCA

TGTGTGGTGAATAAAGCTTGATATGAGGGTTTTATTCTTTAACTAGTGTGTAGCTAACAAATCGACTTTCCCCCGCAGTG

TTTTCCATATTATTTCTCAGTTCCTCGTAACAGTCGCTCGCTTGCAGTTCCCCAGCACTTGCGTACTTTGGTCACAAAGA

AACAACAAGGTTGGTCACCTGTGTCTTGGTGTCTAGTCATTACGAACATGTCGGTGCTTAGGCAGGTAGATCTTCTGCTT

GCTTTTTCTTCTCATATTAGCAACTTCACTAATTTGCAACTCTCGCCCCTTTCTTCCGCTGGTAAATTCGACTCCAATTG

TAGTCCTTTACGGTCACTCAATTTAGTAGCCGACGGTGTAAACAAATTCCCAAATTTTGATAGGAAGAAGAATGTGACGG

ACCTCATGGTGCTTCATGTCTGGATTATTTTCCTAGAAGCCGATGTCCGAGAAATAGCACAGGCTCGTTGAAAGCTTTCG

CAGCATCAGCGATACATGGATGAGAGCGATGCATATTAGGCGTGTGTCCGCGGCCTGCATGCAGTATCTGTTTGGCATTC

AGTGCCTCTCTGACTCTAGTCGTGGAGGTTGTGCCATGCCGCGGACTGTGGGTTTTCCCCGAGGACTCTGGCCTTTATTG

TATCGATGTAGTCCTCGAAATTGTCTCTATCTTTAAAGTGAAACTAACGGCAAAAGAAGGCGGTAGAAACACTCTGTGAA

TACCTTGGTAGTCGCTGCACCGGTGGATTTCTTTATATTACTCTTGGTTTTTAGAATTTGATATTTTTTTTTTGGACGTG

TGCCCGTGGCGATAACCTTACTCTTAGGCATGTGCGTGATGATGTCAAGTCAACTCCATCCCTTTCTATGACCTTGCTTC

GCTCCTCTCTTAGGAATTAGGTTACATAGATCCGTAACGACGTGTCTTTCTTAGTAGAGCAGCAGATAGGAGCACTTTTT

TTTCCGAAAGTGCGACTTCATTTCCCGAAAAATGTAATTACTTTAATATCTCTTTGCTGCTCCTGATCAGTAGTAGTGTT

GAACAGATAAGGTATCTGACATTGTAACCATTTCTATGATAGAATGTTCGTTACGCCTTGCTTTCTTAGAAAGATATACC

TTGTTTAGGTCATAATGTACGTGTAGAACATCATTGCTTTAGAGAGGTGTGTGCTCAATATAATTTGGCATTACTATGTA

AAGTGTCTGTGCTATAATTTACTATGACTTTTACTCTTACTATACCTACCAAAATTGTAGCTTTGACAAAACAATTCCAA

ATAAAAGTGATAGGAATGAACCGTTGATGAGAGGTGTTTATATTATATATATACCTGTATATATAACTAACTATTAACAA

GCTGGCCATATTGTGAAATGGAGCGTTCCACAATGTATGAATTATGTTACAGTGCACTCTTAAAGACTGTTCTCAGTACC

ATTATGTACAATTCTTTAAGATAGTAGGTCATCTCAAATAAAATGATGCACACATTCTCTCGCGGAGGTACACGGAAATG

TACTCTGTACTCAGAACGCACGGCTCCGCCTTGAAATTACTCGTAAATTATATCAACGCGAGAGCTGCAGTTCTATGGCA

TAAATTGCACAGTCATATAGACGTCCTGTATTCGTAGCGGCGTCGCTAAAACTTTTCTAAGCACATACATACCTACTGAT

AGGACTGGCTGGTTACGTTTCTAATTCACTTGTTGTACTTTAATAACGATCAGGCGAATGGGCGTTTGTTGTACCTACTA

CAATTCGTCTATTACTTTATATTCATGGATTGATGTTACTAGGCTTTATCGTACAGCATTTTCATTATGCGCACTGAGTT

GCTATAAGCGAGGTACCCTTTCTCTTCTCTGGCGAGGTGAACACATGGTTCTCTCCGGGTGCCTCGGTGGTGAGTTGGAG

TGCAATAAGATTCGTCTTTGTATAATCATCCTTCTGTTGAGATGCCTCCTAGTCTGTGCTTCTTTTTGCGTCATTTATGT

TCGGTGTACATAATATATTTTTTCCGATGACGTTGTAGTAGCGAAGTTTTCGTGCAATGTAATCTGATAAAAGTGAGCGG

GGCTGCCAAGCTTCGCCGACGCGTGCTGTTTCTTTCTATGGTCATCAAATTTCACTCATTATATGAAGGCGAGGCGCTTG

ATGCGCCCACGACATCACTGGTGACCACTTCATTAGAATTTTGAATTACCTGCCGGAGTTTTGTGCTCTGATGGGGACAA

TTCAAGAATTGTGTGAAATATCGTCTTGACATTAGTGGTTGGTAAGTGGCACTTTTGTAATAAAATAACAAATTACGCAG

TGTATCATTATTCAGGTGCAGTTTATTAGAATATGTTCTAATTAAGAAAGTCAAAATGTAGGCAAGGGTTCTTGACTGAT

AACTAGTTTCAAGCGAATGCGTTCTTTTAGGTTGCAGTCGACGTCTTGACTATTTAGTGATCTGATGTATGAGCCAAGTG

TGCAGAGCTCTCGAGTTCGTACATTGTTGAACTTGCTGATGTGCAGTGTCACAGTGTGTTAGCTGATCTACGTGTGCTCT

ATAATGGCTGACAGTAGAGTTTCAATTGCTAAATTGCGACTTCGCTTCATTATGCTTGGTTTGCAAAAAAAAAGAACATG

CATGTTGAAATAATTTTTAATATGAGCTCCACCACTTCTCCAAAATTAATGGTTCAACTATAATCAAAACAAGT

Twins (PP2A)

>lcl|PH.k31.comp30016_seq0

CATAAATATAGTGAATGCAGTGTCCGCCATAGACTCCGAATTTTATGTATTTTCTGTGTTTTGTGTATCGTTGATGAATA

TCATCAATGACTAAGGTCATTGGTCGAGCTGAATTTCTTAATAGCAAGATTAGACGTGTATCCTATTGCATGAACGAGTG

CAATGGCCGGCAATGGAGCAGACACACAGTGGTGCTTTTCACAAGTGAAAGGAACTCTCGATGATGAAATAACAGATGCT

GATGTTATCTCCTGTGTGGAGTTCAGTCACGATGGTGACCTCTTGGCCACTGGCGACAAGGGCGGTCGTGTTGTCATATT

TCAGAGAGACCCTCTATCCAAGGGCTGCTCCCCAACGAGAGGGGAGTACAACGTATACAGCACCTTCCAGAGCCACGAGC

CGGAGTTCGACTACCTCAAGTCTTTAGAAATAGAAGAAAAGATTAATAAGATTAGATGGCTTAAGAGGAAAAATCCTGCT

CATTTTTTATTATCCACTAATGATAAGACTATAAAGCTGTGGAAAGTCTCTGAACGAGACAAGCGAGCAGAAGGCTACAA

TCTTAGGGATGATTCCGGCCAACCTAGAGACTCTGCTACCATCACTTCTCTTAGGGTACCAATACTGAAGCAAATGGAGC

TGATGGTGGAGGCATCTCCTCGACGCATCTTCGCTAATGCTCACACCTACCACATCAACAGCATCTCCATCAACTCCGAT

CAGGAGACCTACCTTAGTGCTGACGACCTTAGGATAAATTTATGGCACCTTGAGGTCACGGATCAGTCGTTCAATATAGT

GGACATCAAGCCTAGCAACATGGAGGAGCTCACAGAAGTGATCACGGCTGCAGAGTTCCACCCCCGGGACTGCAATGCTT

TCGTGTACAGTAGCAGCAAAGGCACCATCAGACTCTGTGACATGCGAGCGGCCGCCCTCTGTGATACCCACGCTAAAATG

TTTGAGGAGGCTGAGGATCCCAGTAACAGAAGCTTCTTCAGTGAGATCATCTCAAGCATCTCAGACGTGAAGTTCAGCAA

CAACGGTAGTCTTATGATCTCCCGAGACTACCTCACCATCAAGGTGTGGGACCTGCGCAAGGAGAATCAGCCTCTTGAGA

CCTACTCCGTGCACGACTACCTCCGGTCAAAGCTGTGTTCGCTGTATGAGAATGACTGCATCTTTGACAAGTTCGAGTGC

TGCTGGAGTGGCAACGATAAACACATCATGACTGGGTCGTACAATAACTTCTTTAGGATGTTTGACCGTGAGAACAAGAA

GGATGTTACTCTGGAGGCCGCCAGAGACATAGCTAAGCCTAGGACGGTACTAAAGCCCAGAAAGATTGGCGGTGGTGGGA

GCAAAAGAAAGAAGGATGAAATCAACGTTGACTGTTTGGACTTTGGCAAGAAGATCTTACACACTGCGTGGCACCCCTCG

GAGAACGTCATAGCCGTCGCAGCTACCAACAATCTGTACATCTTCCAGGACAAGTAGTAGCTTGTGTGTGTGTGTGCCAT

CGGTGCTGCCCTTAGTATTGTGCTGCGGTGACTCTGTGCTGCACTTGCCGCTCCAGTGCTATTGTCTCTCACCTGTGACT

GCTGCTGGGGTTGTGGGTTGTTGCCACGTGCGGTTGTGCTTTGCCGTCTCATTCATTGGCAGCATACTTGACAAACGCAG

CGCGTGCTGCAGTACTTTGCTGCCTGTAAGGCCTGTGTTGCAGCTGCTGCAGCAGTAAGATATAGTGTGCGCACGTAGTT

GGGCAACGAACTTTTTACCTACCGTACGCAGTTTCTGAAGTGGTAAAATTTTACTTGCACAGGCCGTGTGACAGCGAGCG

AATGCACGCCGTGCCGTGCAGACTCCATGGCAGAGAACAATAGCACATTGCGTGCTGCGCACGCTGAAGCAGGAAATTAT

TGCAGTTGATGAGACGGTAGGTACCCTAGCAACTTTATTTGCCGCACCCGAGCTGTTTTATAATTTAGGGTTCTGGGGCA

GCCAACAATGTTGCTAGGGTAGACTGTTCAATACGCAGTATCGTGTCCCAAACGTGGCCGTTTCGATCGCAACCTCAGCC

CCTCGATGCAGTTACAGGTGAAATGAAGACGAAAACGACTGACTCATCTAACACCCTCCCACTGATATTTCTGGCTGTTA

GATGACTCTGCCGTTTGTTCCCTTCATACATGAGCACTTGTCGCAACTGTGATGTACAGCCATACTGTCGGTGCCCTCCC

AGAGCGATTGTATTTTACTACTGATGTACTTTATTTCCTACCCCTGAAGTTGTCCTCCGTCTGATGATTTATTTGTCTGT

TCGAGGTGCGCCTGGTTGCAGTTCTGACGACGTCGTCGTGGTGACGTCAGCTATAGTACCAGTGCTGTTAACGTTCCACT

GACGTCTGTTTCGTTACTGTCTGCTATCCTGGTGATGTCTTGCCGACGTCACTTAAGGTCTCAGTGCTGTTAACGTTCCA

CCGACGTCCGTTTCGTTACGTCTAGTAGACGTCTACATCCACTGCCATTCTTCTTCTAACCTTTGGTGTGCTATCACTCA

ATCCACTTTTGTGAAACCCGCTTTTAGAAGCCACTTCCTCCATTAGAAACGTCCCGATAACTTTCACCTTCCTTTGTAAA

GAAATTCATCTCACATTCACGATCCTCTTCCTGAAACGTGCACTTATTCACTTTTATAGTGTAATTTGTGCTTTGAACAT

TCACTCCCCATAGAAACGTGCATCGAACTTGCACCTCCTTTGTGAAATATGCGTCTTACATTCACCATCCCGATGTGAAA

ATGTGCTTTATAGTTGACCCTTGCAGACAATATCTCACGCACACCTCTTTCAAGTGGTAAACCGTAGCTATGTCATTATT

TACTGACACTCCTGAGTAGGGGGAACATGGTGTATGGATAGTGCTACTCGGTAGTAGCTCTCTGGTTATAGGGCGCTCCA

AGTATTTATTTAGCTACATTGCGATACTGCTGCTGCCGCCTCCGCCGCCGCAGTTTTTCTAAGAGCTTGGCCAGTTCTTG

GCCCTAAACCGTTACCTGAAGCTCCAAGCTTGCTGGTTTCTCTTTCCTTCCTTTGCTTTGGAACCTCCCCTTAGTGTTAT

TGCTGGTGTTGGCAATCATGCCTGAGTTAACTCAATTGTCACGCCTTGTAGGATGAATGAACTGGAGGTCTACTCGCACA

AATGCGATAACCCTCGTAAGAATCACTATGATATTTTGTTGAAATTTACAGGGTTACCGTACTTTGTCAAGTTATCCCTC

CCAACCGTGTTGTTGCATAGAGGTAAAACTGGTTTGTGCTGGTCGTGCCCTCTGACCTCAGTAAAGGCCTTCTGATTCTC

CCATGTTCATCGATGAAATCAGGCCCAGTCCGTGTCAGAACTTGTCTGGAACTTTGCTTCTGAGCGCTCGATATGCGACG

TTGAATCGAATGATAGGCGTGTGCCGTTAATCAATTATGCCTGTAACTAAATCCATTGTGTTGCGGCGGATGCTTGGCAT

TGCAGTCTGGACAAGTTAAATTTAAACTTAATCAGATTGGCTTATCTCGATAGAAGTGGGCAAGTCAAGGGCACTTTAAC

TGAAGTAGCTGGCAGAAGATGAGTTGAACTTGGCTGTCCTAAAGAGGCGCTTAGTGATATTTTTGTGAGAAAGAAATATT

TAAAATTGTTAGGTATTCCTACTGCCTTCGCTTGTAGTTGCTTCGGATATCCCTCTGTAAGGGCGGATGCTTTACGTGAT

CCGTGTGTGACTCTGCCAACACAGTATAGGGATCTGGAACGCTGGTAGGGCTTTCTTGCAGTGTCTTCTATAATTGGTAG

GTTCATAGTTATTGATATTGTCAGTGCAGAGCAAATTAATGATGGTGAAATAATTTGTACCTACTGTCGGTGTGGCGTCG

TAAGGTTGTTATTGATGCTATTTGATAGTCTGTGGAGAAAGTTTCCCTATGTTGACATTCTAAGTTATGCTGTCTGTACG

GAAAGAAAGTATTTCCTAGTTATTCATACTGTCTGTGCCGAAAAATTAAGGTGGTGTAGTAATTTATTTACACTGTTTGT

ACGGAAAGAATTCTAGCAGAGTTATTGAGTTGTCTCGAATTGAATAGGTTATGCCGTATGTTTTGCCAATTAGGGAAATG

CTGTGTCGTCTTGGTGTATATAAGTACTTATCGATGCAGAATTCGTGTATTGGTTTTGCTAACAGTGTATTGATGCGAAA

TTTAAGTACAGTACTGGTTTGCCTACAGCAAATTCTCGATACTGAATTGGAGTAGCTACTAGTTTAGTTTTTGCATCAAA

TTGTTGACAGAGAAGAGAATCCAGTGCTTGTTTTTGCTCACCGCAACTTGTGGATAGCTCCAAAATATACCTAAGTATCG

GTCTTACTAAGAATAAACTTTTGGTACAAAGTTGGAGTATAAATTGGTTTGTTTACGTCTTGTTTCAGACAAAGGTTGAC

TGTTCCCTTCTACCCAAAGGCTGTCGTGGATATAACCCTCGAAGTTCTCGGTGTTGGCGATGGCCGCAATTGGTGCGGCA

AGCGTGTGAATTCAGTCGTAACTTTGGTTCTGTCGTTGGTGCCATGCAGCATAATGCCCCGTACCTTTTTTCAGATAGCT

GTAGTTCAGTTGGCTGCATTTTGACGATGTTTACTTGAAATACTTTTGTGTGAGAGCACCCCTGATTCTTGATTCTCGGT

TTCTGTTGTAATGAGCGTCGGGCCGCTACGGTCAATCCTTCGTATCATCGCTTCAGTAAACGGGATTCCTGTATGTTTGT

ATCTATATATATAATTTGTTCTTTTGCTGGAGGTGATTTTGCCCACTCGTCTGTAGCTCTTTTGTTTCCATTGTTGCTTG

TGCTGTGGAGGGCCTCTCTTGTGATTCATTTCGTTTACCACCGCTTGTATGGCCTCCCTCTGGCTGCTCTAGAACCCTGT

GCCCACTTAGTCTGGTTTCTCTACGACATTTCCGCTCTCATCTTGGCCAGCCTTCAATATCTGATTTTGAAGCGCTACCT

CTCTTCAGTTGCTGTCTTGAAAGTCCCGTTTACAGAATGTATGGTGTGCTCTGCTGTTAATGTGCAAGACCGCAGCTTTG

CAGTTTGGCCCATATTACATATTGTGCTTCAGCTGCTGATGCCCGTGCTTGCGATTCGTCTCCCCGCTTTATTCTACCCA

CCTTTGTGTCGGCGGCCGCCCTATTGGCGCTTCACTTTAGCATCAACGTAACGTAAGTG

Shaggy

>lcl|PH.k29.comp7576_seq1

AGTTCAGGTAGCAGCATCACAGCTCTAGCGGCTATTTTTGTTCCGACGTTAAGGTGAAAGAGTGAGAGTGAGAGGCGGCG

TGGCCAAGCTTCTGGTCCCAAGCAGTTGCTCGTTGCCCTCACACACCTCGCGCTGTTCCCCTCTGTCTCTCTCTAGCACT

GTCCCTCAGGCTTCTGCCATGAGCCTCCTAACCTTAGATTTACTGGATAAGAAGCCAGAACCTTACCACTATCTTAATTT

TGTGGAGGAGGAAGATTGGGATTTGGAAGATCATGACGAAGACGCTCTCACTGACGAGCAGCTTGAGCCTCAGTATATAG

AGGATCCCTCGGATGAGTTAGAGCGAAGAAAACGAGGAGGTAAAGATGGTAACAAAGTCACGACTGTGGTGGCGACAGCC

GGCCAGGGCTCGGATCGGCCACAGGAGGTGTCGTATATGGACACTAAAGTCATAGGTAACGGATCCTTCGGCGTGGTGTT

CCAAGCCAAACTTGTCGAGTCTGGCGAACTCGTTGCTATTAAGAAAGTTCTGCAAGACAAACGCTTCAAGAATCGGGAAC

TGCAGATAATGAGAAGACTGGAGCATTGTAATATAGTCGAGCTCAAATACTTCTTTTATTCCTGCGGGGACAAGAAGGAT

GAAGTCTTCTTGAATCTCGTATTAGAATACGTACCAGAGACCATATATAAGGTAGCCAGGCATCATAGTAAACAGAAGCA

AACTATACCAATCAGCTATATTAAGTTGTATATGTACCAGCTGTTCCGGTCCCTGGCATACATACACGCCCTCGGTGTGT

GTCATCGCGATATCAAACCCCAGAATCTGCTGCTTGACCCGGAGACTGGAGTGCTTAAGTTGTGCGATTTCGGGAGCGCG

AAACATCTCGTGAGGGGCGAGCCTAACGTCTCCTACATCTGTTCCCGATACTACCGTGCCCCCGAGCTCATCTTCGGTGC

CACAGATTACACCACCAATATCGATGTGTGGAGTGCAGGATGCGTACTGGCAGAACTCCTGCTGGGGCAGCCCATCTTCC

CCGGCGACAGCGGTGTCGACCAGCTCGTGGAGATCATAAAAGTTTTGGGGACGCCGACGCGTGACCAAATCAGAGAGATG

AACCCTAATTATACAGAGTTTAAATTCCCCCAGATCAAAAGTCATCCGTGGCAGAAGGTGTTCCGGCAGCGAACTCCTGA

AGACGCGATCAACTTGGTGTCGCGACTGCTGGAGTACACACCGAGTGCACGCATCTCGCCGCTGCAGGCCTGCACCCACC

GGTTCTTTGATGAACTACGTGAGCCCAACACCAGACTACCCAACAACAGGCAACTGCCTCCTCTCTTCAATTTCACCGAC

TTTGAACTGAAAATCCAACCGGAGTTGGCAAGCAAGCTGATTCCCAGCCACTATCATCCAGAGGGCAAGAACAGCGGTGG

CGGTAGTGGTGGTGAGACGTCCTCAGGCACTGGTGGTGGTGGAGGTGGAGGTGGTGGGAGCGCCACCGGTGTCACTAGCG

CCACAGCAGCCACTGCTGCTGGAGCAGAGTAGCTCGCGCCTCATATGGCTTTGCTGCCTCACATTCTTTTAATGACTTAT

AGTTTTCTTTTGTATTAAATTTCTATCCTCTTGTGAATATTTTTTGTTTTAATTGCAATTGCCGGACGCTTGGCTTTAAT

TTTGTGTGTGACATAATAGTGACGCAAAAGAAGTTACACCCGGGTAAGCATGTCTGTGTGTGTGTGTGTGTGTGAAGCCG

CTGATCTGTCGTTGCTGTAGCTGCAATGGCACCACAGCTTCGTGGTGTTACGAGAAGTTTTTGTGCGTGCGCACGCAACT

TGTGTGCTCGGTAGCTGTTGTGTAGGACTTGAGTGTGGCCGGCGGCTGACCAGCGGCTCTTGTAATTGCCTCTTTCTGTT

CTTACTTTGGTGTAAGCGAGCTCGCTCTGACGCGCGTCTCACCAGCGACTGTGTATTGCATTAGCCGTACGTTACACGCG

GCCTGCTCGTAGCGACTTAGCGTTGCTCGTACAAACTCTGTGTGTGTGTGCTGCGTGTGTCCCTGTCGAGTGTGTACGAC

TTCCGTCAATTGCAATTTAGGTGGCAAGGGGATAAGAGACAAGGTAACTTTCGGTCTCGATCCCCTATAAGTGAATGGCA

GTCACTCTAAGCTAAACTCAGGTATACATGTAAAATTAGAGCGTGTCGAGGCATTTCATTGCTGGACTGGGCCTGTTTTT

CCAGTAAATATTATGCTTTAGTTCACGGTTTCGTCGGCAGACCATAGCATGCCTGCGGAATCGGCAGTCTCGGCGTATAA

GCTTGAGTGGACGCACTCCTCAGTTACATGGTAGAAAACAAAAACTTCAACGTGTATGTAGTGTTCCATTGTTGGTCCTG

ACACCTATGAAGACTATTAGGCATCTAGGCACTAATATTTACATCTCCTTATGTCCTCGTTAGAAATACTATGTTGTTAT

TACATTCCCAAGACGAGACTTTGACGAGGTTTCGCGGTGCCGTGTCGTTGCACTGGAGCGCTCACTTCATACGTCTACAG

TAGTTCCAGATTGAATTAAATTTAGCTAAATTTTCTGACCCGCCGTATGCTATCGTAATTTTTGTACCGGAGCAACGATG

TAACCACCACGCGCAGTCGCTGATTAAGAACTTGCTACTTTAAAAAAAAAAAAAAAATTTTGTCACATTCTTCCTGTACT

CGTTCTCTCATAGCGTGGGTCTACGTGCATCATTGCAAATAGCGCTGTAGCTACCTACGAGAGAGGCGGGATAATTCATG

AGAGAAAGGTTAAGAAGAAGACTATTTTATAAATTCCACCACTGTAACAATCCAAAGTTTTATGCGTGGATTCTTCAGCG

TTGATTTTGCTGCTGTTGTCTGGCTGCCTCATACATCGTCTCCTGCAGTACTGTTATATTTATTGAAGCGAAATTCCAAT

ATATGCAGTTTTCATCTTCAGTAATCTGATCTTCATACAAATGAGTCAGTTCATAAAATGTCTTTCTTTCAAGGTTAAAA

CATCTTCTTTGTAGACGAGGGTCTCCTGTAGCTCTTTGATCTGTCGAACTAAACCGTACTGAACGACACCCACAACAACA

ACACATCCTAACACACCTGAAATTCGCTGCATCCTTTAGCAGATTCCTGTGGCTGTTTGTCTGTTGCTGAATGAGAGCAT

CCGGTGCCGTGTCTACTTTAAAGTCCATGTCGGCTGACCTCAGATGGTAGGACTTAGGGTCACCACGTCACTAGACTAAA

TAGTCCTTACAGTTTCGTCTCGACTAGTCAGGCGATCACCGCCGTGCCTCGCCTCGCCTCCTATTCTTCATTCATTAATA

GAAGGATTCAAATCTCGACGTGAAGTATAAACTTCTCAGGCACTTTTAAATCTGCCAGACGTAGTGTAAACGACTATCAC

GCCTACAGGCAATGAAAAACAACAACCCTTCCCCCATAAGTGAAACACTGCCCTATATAGCCTATATATCCACCAACTGT

TACAGATAAAAGACCACTCATATAGAATAGTTTCCCTGACGATACCACTTGTACTGCTGAGCTCTTAGTAATATCAGGAA

AAAATTGGCAGTTTTTCACAGAGGATGAGTTTTTTGGAATAATCAAAATTTTCTTCCAAGATGAGCGTTATATTTCACTC

ATTTTGAAGTTACCAACGGGCATCCAACTAGTCCAAACCAGCTAATCGGTCGGATTCAATTCGGTGTAGCGTCATCCTTC

AGCTGCATTTGATTTCGGCCTGGAGACGTTTTGAACAAGAAATGTGCGGGCATGTATGTAGTGCACTATAACTTGGATTA

TTTCATTCTGTGTTCAAACGAGTGCTGGTTTCAGAGCAGAGAAATAGCTGGGATTTTAATAAATGAGTAACTTTTATTAA

GAACACATAGTTTCTTATCTAACTTTTCAGCTCTTCCAATTAAATCATTAAGAACACAGTGGGTTCAAAAAATACGATTT

TGCCCATGTCTGTTGCTCCAAAGATGAACCATGTTGAGAAGTCCATTGGTACAGATCAGCCTATAAGGGGGGGGGGAGGC

AGAAATGTTTGTCATTGAAAGAACCGAAAGATTGTAAGAGATTTCTTAACCTAGTTTGGGTCCTAGTCTGCGAAGGAGCT

CCTTCTTGTTGGAGAATTTATTGGCGGATTTGAAGTGCTGTTGAGCTCTAACTAGCTTGAGAATCTTTTTGTTAAACTTT

CTGAGAATTAAAAAACTGCATAACATGGCTCACCATTCTGCACTTTCGAAAATATGGCTTCTATCTGAGAAAAATTTTAG

GCTATCCGCCAAAGCAAACATTTGAAATAGGCTACTTTTGGGAGTTGTTACTGCACAGTCAATACTGAAAAATTGCACGG

AGAGGTACCGGGTGCAGCGACCCCTGCAGGTTAGGATCCTTCTTGCGTAACGCGTTATCCTACTTGTAACTCGCTCGGTT

CTTCAAAAGGCGGCCTATTTTTATAGTTTTTTTTATTATTTTTCAACCAAGCTAACGCAACGTGTGAGTACTGCGACCAG

ACTCCTGTGTCTCCTTACATTTTCACTATATTCGACTACCATTTGCCGCTGGTTACCCTGTAGTTTGTCTCGAGTGAGCT

AGCAATAGTTCAGAGAGAAACAGCCTGGGCTTAGATCTCGTTAGGAATGGAAATTCAAGGTCTGTTGTTCAGTATCTGAT

GTACCACTGCAACGCGCGACCTCTCAAGCTTATTTGCCTGGACTGTACGTTGCCTATATTCCTACTCGCCTTACAGTCTT

GCCACTTATAGACTGTATATACGTTGCCTCTATTCCCAGTCTCCATCGCTGTTATATAGTCTTTGTATCTAAAATACGGC

GGGAAAACCCACAATATAAGAATTCCTAGTCTCCATTGCATTGCCACCCAAGGTATGATAGAGCCGCTGTCTTCATCTGT

TCCTATACTTGTGTTGTAAGTTTCACTACTGCTGCTCATTTGTTATTAAAAACTAGAAGTTCTCCTGTCCTTTCATTGTT

GCTGCTCTTGATGATTAACTTGGCCGGCTGCCGCCACCAGTAAACGACGCTTTTTTAGATCGTGGAGTACATACAGTTTC

TACTGCATGTCTAGGATTGCCGGAGGTGCTGCATCTGTATGATTCGATAACATGTGCGTGTGTGTGTTGTAGCAAGTGGG

TCTCACATTGTGTCATAGCATCTTAATATGCAGTTGTGTACAGA

Supernumerary limbs

>lcl|PH.k25.comp29022_seq0

TTCACAGTCCACAGAAACTTGTTGGATAGTGAGTTAGTCTCGATTGCACGGTGGTAGTGAGCAATGAATCCAAGTTGCTT

TTCTTGAAACTAATTCAAACTTGTGATATTGGTTTTAGTATTTATTAATTAATCGCGGCAATGGATGTCGATCCAATATT

AGAAGATAGTAGAAGTCTAGAATCTTTAGAGGGTAGTGGGCTGAACACAGTACTGTACGACAGCAACAGTCGGCACTCGG

CAATGGACACAAGCACGCCAGTTCTAGATGACAGTCACAATGATAGCAATACAAACATTTACACTGGCTCAATTGTTCCA

CTCTGCACCACTACTCCTGCTGCTACCACCACCATCACCAACAATGCCAATAACAGCAGACGAAAGAAGGAAAGCAGAGG

AGAGTACACAAGCCAGAGAGAATCCTGCCTCCAGATGTTCGACCATTGGAGCGAGCAGGATCAGCTGGAGTTTATGGAAC

ACTTGCTCTCCAGAATGTGTCATTACCAGCATGGCCACATCAATGCCTTTCTTAAACCTATGTTGCAACGAGATTTTATA

TCTTTATTGCCAAAAAAAGGATTAGACCACGTAGCAGAGAAGATTTTGAGCTATTTGGATGGCAAGAGTTTACGAGATGC

AGAGCTCGTGTGCAAGGAATGGCAAAGAGTAATAGCTGATGGCGTCCTGTGGAAAAAGCTCATTGAAAGAAAAGTCAGAA

CAGATCCTCTGTGGAAAGGTCTCTCCGAGAGACGAGGATGGGGTCAATTCTTGTTCAAACCTCGACCAGGAGAGCAGCAC

CCCGGCCATTCCTATTACCGAAAAATGTACCCTAAAATTATTCAGAACATCAAAACAATAGAAGCCAACTGGCGAATGGG

ACGACACAACCTGCAGAAAATCAATTGCCGGTCCGAAACATCCAAAGGTGTATACTGCCTTCAGTACGACGACCACAAGA

TCGTGTCCGGCCTTCGAGATAACACCATCAAGATGTGGGACCGCAATACTCTGCAGTGCTATAAGGTATTGACGGGCCAC

ACGGGCTCAGTGCTGTGCCTTCAGTATGACGAACGAGTCATCATCAGTGGGTCATCAGACTCGACAGTGCGTGTGTGGGA

CGTACACACCGGCGAGATGACCAACACTCTCATCCACCACTGTGAAGCTGTGCTCCACCTCCGCTTTACCAATGGACTAC

TCGTCACTTGCTCAAAGGACCGCAGCATTGCCGTCTGGGACATGGTCTCTCCTAGCGAAATAAACCTGAGGAGGGTTCTT

GTTGGACACAGGGCTGCTGTCAATGTTGTCGACTTCGATGAGAAGTACATTGTGAGTGCCTCAGGTGACCGAACCATCAA

GGTCTGGGGCACCAGCACCTGCGAATTTGTCCGCACTCTAAATGGTCACAAGAGAGGCATTGCCTGCCTGCAGTACAGGG

AGCGTCTCGTTGTCAGTGGATCGTCAGACAACACAATACGGCTTTGGGACATCGAGTATGGTGCCTGTCTCCGCATCCTC

GACGGCCACGAGGAGCTAGTGCGATGCATTCGCTTCGATAACAAACGAATCGTCTCTGGAGCCTATGATGGCAAAATAAA

AGTCTGGGACCTGAATGCTGCTCTCGACCCACGGTCTCCAGCTGGCACGTTGTGTCTGAGGACGCTTGTGGAACACTCAG

GTCGGGTCTTCAGACTACAATTCGACGAATTCCAGATCGTGTCTTCCTCTCATGACGACACAATCCTAATATGGGACTTC

CTTAACTGCTCTCCGCCTGACACGCCCCCTCCCCAGGTGCCTCTCGGTGGTACCACGTCGCCTTCGGTCGCCCCCGAGTC

TAGTGTAAATCCCTCTTCCCATAATACCGTAGCCTCATCTAATATGGCCTCAGCAGCCTCCAGTGCTTCCCCATCTGCTG

CCTGTCCTGCAAATCCCAACCTCGGGGCTCCTTCCAACTCTATGTTTTTCGGCGGCGCTGAAGGTGTGCCGGCCCCGCTG

CTGTCGCCTCCGCCTCCACTACCGTCCAATGACTCTATGGATAGGTCAATAGATGAAGGGCAGGACTGATGATGTGTGGG

CGTTCACTTGTAGATAGCTACGATCGTCAATGGTTAAGTGCACTCTCATTGTTATTGATGTTGTTTTAGTGGCAGATTTG

GTAACTGTACAATGCGCACTTGTGCTGTTCTATTGCTGTACTCACAATCATTATCCGCGTCGCTAAGAAATCTTTCCACA

TCATGCCTGTCTCACCGCTTCATTACCGCGACTGGTGCAGTGTATTCCAGTAACAATCATAGCCGCCGCCTATGTATCAT

TTTGACATTACCCTCATCACTAAAGTATCGTTCAGTATCTCGCCTCTTCATTAGCTACCACAACTGGTGCAGTGTATTCG

AGTTACAACTGTTTCCTTCACCGATGTATCAAGTTGGGGCGCTTCATCCCAATCATTCGTTAGGTACCATAAACACTCAC

GTAAAACATTTCATCACCATAACCGTGTCAGCACTACCTTACCAGCCCTCTCCACCGTGATTGTTCCAGTCACATTATTA

CCTTTGCACTGGGGACTGTTGAGTTACAAACTTGTCGCTTTTTCGTGCCCCTCATACCGGTATGCGTGTCTTAACGTGGA

ATTCATTGGCGGGATCTGTGTTCAGCCGAGCTGGTCTCTGCAGGCGTTCAAATCATCCGTGGAGACTTTGCTGAGTTCTT

CTTCTTCTTTTTGAGCAGCGCTTTGGGCGGCCAGGGGTTGTGGCTGAATGCTTCATTTGGGTTTTTGTGACCAGTCTCTT

CACGCCCTGGCATATAATTTCTATTGATTAGCTATAATATAATAATCGTATTTACGTCATCTCCGTCTATTCCCGCTTAA

CGTGAACGTTATCTTAACACTTTGCCGACCTTCTGGGGGCGGTATTCCGCTGTTAAATAGTGTTCTTGGAACGGTGAATA

ATTCAGGCTGTAGAAGGTTAAATGGATCAGTAAAGTACACCAATTCTTGTCCTGCTCTTCCTCCGTTGACAGACTACATA

GTTGCGCCGAGTCTTTCACTACCGCGCAGCGTAGCGAACGCACCAGTTACACTTGCTGTATCTGTTATCTCATTACCTAC

ATTTGCTGCATCGCCATCACACAACGTGCACTTGCTACATCTACATCATCCTCTCACCTACACTTGCTGCATCTACATCA

TCTATGGAGTCTGCATCTCCACTGAACTACATCTTCATCATCCTGTCACCTACATTCTCCTCACCTGCTCCACTCCAGCG

GCCACTCAGCTTCAGCACTCGCGTATCACGTCGCTTCTCTCTCTGCTCAATCCTTCTCAACTTGATCTTCTTCAACTTCA

AGGCCGCTGCCATCTTCTTCCCATTTACGAAACTCATCGTTTTCTAACTCATCAGACAGAGGCAAGAGAAAGCTTATAAA

TTAGCTCTTAGGTTTGCTAGCGTCTCTCATTGCTCTAGGTAAAGTTTCTTTATACCGTGTAAGTTAAGATCCATTGATTG

GAGATCATGTGCTCCAAATATTACTGTTAGTAGGTACTTGTGCTGTGCTGGGGAAGTGTAAAGTGTTGCGTTAAACGCCT

TCTTTAGAATATTATCTCGATATTTTTTCTATTTGCTTTTGGCTTAACATTGCTTAAACTATTTATATTGAAACTTGTTT

TCCGAAAACGCACCGTCGTTAGTTTTCGTAACTGCGCTTGCGGTGGTTTTTGGAACAGAAACAATGATAGGAACATTGCG

TGCAAGATATCATTTACTGTTCATGTGGTGTCACATTATTGACGTGTAGGGTAGGAAAACGTGTATCAGGTGGTAATGGA

AAAAGTTTGTGTGGTTTTGTAGTGGATTGTTAAACTGAAAAGCGTGAGAAGTTAATGGGATAACTACCTGTAACACGCTG

TGTGATGAAATAGGTATTGTGCTCATCCCAACTACGACTGAACTTGGAAGTGACCTCACTTCACTAGAACCTGCTACCTT

CACGATAAAGCCTAGTCTCGGTTCCATGTCGATAAATCTGGCAAGGATTCAAGCTGTTCTTTCTTCGAAGACGAACCTTT

GCTGTCAATTATCGTGATGGCGCGCGGGGCATCAAACTTTGAAGGAAGTCCTTGCCGTTTGTAATGCGGTCGACATCTAC

GTCATTGTATCGCAACCGAAGTACCACACCCCAAGCATAGCTATAACTTTCCATTTCTTATCTACCAATCTAAAATGGTT

TCGTGCTCTTGGGTCATTTTAGATTTGTAGAAAGGAAAAGTTTTAGCGTAGTAAAAAGCTGAGGTCTTTTCCTCCCTCAA

GCATTGCAACATTTAGTCTACTTGAATTTGAGGCCGTCTTTTGTGTCTAGATACCAGAATAATGGAATATCTGTATACGT

ATTATTTATCATCGCAGGTGTTATTAACAAATATGTTTTTTTGTTTTTAAGAGCAAAGTCTCACGCCAAGTTCAGAAGCG

TGTCTCCGTCACAGTCTAGGTACTTAGATTTCTGTAACCTCCCAAGTCGCTTACGGTGTCGACAAGCGGTTGCTGTTAGC

TGGTCCCCTCAGGGGGCATAGGATGATGGGGAGCAGCGCCCCAGGGGGGACCCTGTTAGTTACTTTAGCCATTAGCTAGG

CTGACGGGACGCCTGTCTTTCGTGCCCAATTTGGTTCTTTTGATTTTCATATCACACTCTTCCTGATGATGCTTTGTTCT

GTTCTGCCTCTTCTTTCCCGACTTGCTGTTCCTATGCGATGC

Timeout

>phaw_30_tra_m.019341

MTAAIIEAELVAACSTLGYSDGKKYVRDPECLEVIRDLLRYLRRDDNYHQVRLALGETRVLQTDLVPLLREHHTDFALLELLLRLLVNLTTPALLIFHQEIPEDKAGMQRYIKMQTQQQGFKEAFTEAAVWASIATVLSSRLQDGANRDPESENIIEMCLVLLRNVLSVPPSRQDSLRTSDDADVHDQVLWSLHLAGFPDLLLYLSSSTEESDLSLHTLEIISLMLRQQDPQALATSALHRSAEEQKKDEAALVRARESEKARRQQTVRKHCSSRHSRFGGTYYVRNMKSISDRDIITHKPLTDISAINFDENKRGKKVPKNRAPLPDSTTTRRSTLAIRLFLQEFCVEFLNGAYNNIMNIVKSNLDRARSQEHDESYYLWAMKFFMEFNRHHEFKVELVSETLSIQSVHYVQTNIETYHEMMTTERKKIPLWGRRMHNGLRAYQEILMTLTAMDRCQDTAVRESSLALKSKLFYVVEYRELPLVLLLNYDPAKMSKGYLKDLVETTHVFLKLLEGMCKKTRQLMVQLPEKKKNSAKKSKSTKKIIAPPTPEELEEKWGSLADELSAILQGEAGDLPVVVPFDALSELSEDEQKEKAMRKVNALLRQSDLREAVALFRASREVWPEGDIFGAQDVDAPAEFACLREVFMAELNPVEQPENPEESEPEEDEEDELAASAVAESSAHISERELDFPAFVRRFAHIKILQSYSWLLKSYATNAPLTNHYIIKLFHRIAWDSKLPAVFFQSSLFVTLHAAMTDPAKSSNEIIGQIAKFGKYIVRQFFQVAETNPKVFMELLFWKNYKEAVDIECGYDAPVATKAVKSLWSEEEEDELTRLYEEFKEKVDPEGAKDLADHIMENLIRQDRSRRLVIKKLKDLGLISGLRELRNKPARVKGNQWSDIELEELRVLFEEHKNDIDPMARILDFMVNPKPKHRVVEKLLELGIIQDKKEVRKKKIPKPKQQKAKKKGNNFGEQFLAANRGSDDEKSDVDTDEDSSSEDEEQSASTRNAPTPIPVVTPHLVSAALTKVRSLGFQESIQWLVEIMTEVADDREADNDFSAVPILAISEEQSKAIEDEDFQALMKIIGIQPPQSHEEMFWRVPEKLSVDNLRRRAQYLTQGLEGGLPVSSSQDQSMSTSAPSAGSGDSGGAAAEVCSLVDGDQSQDDKENRNILNVGSVLNESSDELSDEQLPVSSGRDNSRASVVMPKRPLEDHSANFDALLSSQPPKKTKKRRIVVDNSDDDDDDDDDDEDEEDSDEKNGESDAASLAKKKENSDADEEDNAESELPRERVTRQPSIIVVEQMEHGDSGCPNSASDGRYLSTSASCDHLNYNSASDTSERYFTADGNGIDTGDDLDTPKRRSLSKECLLKASTDEELARRLCELKLHRKSSSKESLHRLNFKRPSAETLQTFRSCERLTEEDDYGLTMEANPKKFKQEWLARSLTTLSPKYQAKAQECSVMSCLAKFTAPELLASNNKIICQNCTKLRNISGNNPEKDGPVRSPARKQLMVVCPPAVLTLQLKRFHHDGVHLAKANRFVHFPLVLDLAPFTSTIAMSPGTRLLYGLYGIVQHTGRLHNGHYTAFVRSRPITANRPPPTAYLTQHPLDSASLETDPPPRLPPSKTATTVPAVEELEAEIRKDQWYHVSDAHVTHVTIDRVLKAQAYLLFYERIV

Vrille

>lcl|PH.k27.comp15075_seq1

GCTACTTGTTAAATAAGGTTGGTCTGGTGCTGCAGGGCCAGTGTGTCGCGAGGTGTTGTAGGAGGTGTAGTGTACGTGTG

TCTCGTTGTGTGTCGCTCCTGTGGTCAGCGCTAGTGAATATCTTCACTTGTTGCGTTGCATGAGTTAACTTGTTGCGTCA

TATTGGGTGCTGAGAGTGTCTTCAGCGAGAAACCATGGTCGCCGAAACGGTGAAGAGCTACCCATACCCTGCCCTGCTGC

CGCTCGCTCAACACGGTGTCAACTACTCCTGCGCCTCGACCTCCTGTTCACCAGCACTTCTGCCCCAAAATCTTGGCCAG

TCTTACCCACAGGCCCCAGCCACCATGGATCCAAACCATCTGAGCCGAAGTAGAGCCCTCCCTAATGCTGGGTCCACACC

TTCTGGCTTAGGTGCTTCTGGGATGATGGCCGCCACTCCTGCTTTGCTCAAAGAAACCATGTTTGCTCAACGTAAGCAGA

GGGAGTTCATACCTGACAGTAAGAAGGATGATTCCTACTGGGACCGACGCAGAAGAAATAATGAAGCCGCAAAGCGCTCA

AGAGAAAAGAGGCGATTTAACGATATGATCCTGGAACAGCGAGTGCTGGAACTCTCCAAAGAAAATCACATACTTCGAGC

TCAATTGAGCGCTCTGGAGAATAAATTTCAAGTGAAAGGAGAAGGATTAGTAAATGAAGAACAAGTTCTTGCTTCCATGC

CTCAAGCGGATCAAATCTTATCTCTGACCCGAAGATCGAATTTATCGTTGCTTTCGATGACTCCTCGGACTTCTCTTTTA

TCCTCACCATCCATGCCCGCCTCGCCACCAGTGTCAGCCCCACAGCAGTCTCTCACAGAAGATGAACAATTTTCTATTCC

ACAATACAGTCAGCACCAAAGTCATCTTGAGACGCACCTTCCCTCCCCTTCTCAGAGCTACACCAGAGCCCACAGCCCCG

AGTACTACCCAACGGAGCCACCCGCTCCTTCTCACTCACACGTTAATTTCTCCTCAGAATCTCAAGAGATGTTCGAATCC

ACCGCATTAAACCTTTCTTCAAGATCAAGTCGTTCCCCCAGCAGCATGGACTGCTGTTACGAGCAAGCGCGCTCCCCTGA

TATGGGTGGGTCATGCCTTCCCCACAAACTCAGGCACAAGACTCATCACACTCATGCTCTCTCCAACCAATTCAACAACG

TTGCTAATACGTCAACACGACCACGGTCGGCATCACCGGACCAGCTGCCCCCAACTGCACCTCACTTCTACCAACAATCG

CCTTCGTCAAGCATAGTCAACGTGTCACAGTCCATACCGAGGTCGTCTACTTCCCCCCAAACTTCTCACCTACTTTTCCC

TATTAAAAGTGAACCCCTTTCCAGAGAAACTGGAGAAGAATCGCCAGGATCTTCGGACGATAGGGACTCTGGCATCAGCT

TAACATCTTCACCTCCATTGCCAGGAGAGCAGAGCTACCCGTCGTCCAACCGAGAGTCAACCGAAGACATGGAGTGTGAC

AGCGAGCAGCAGCTGCGCGCCGAACTTCACAAGCTCGCGTCGGAGGTGCGGTCACTCAAGTCTTACTTGAGTCGAAATGC

CGACCCTCACCGGCACCAGCAGCAGGACACTCGATAGCTTCGTTACTCTAGACTTAACTTTATATTCTTGTTGCTCATCA

TAGGTGTGAAGTGTGAACTAGTAATTAGGTTTTGATGTTATCAAATGTCGCTCAAACATGCACGTCAGTCCAATTTGCTA

CTAACTTAGCAATGATGAGAGGACTGATGCTGTGGTATCTTTCATTTTTAGCAAGTCCCACCTATTGTAATCTGCTTAAG

GCAGTCGGTTCAGTGAACTAGTTCACTTTTTTGCTTCGTATTAACATTACTTTTGCCCATCACTATGTATTACTATGACT

ATGTAGAGATGTTGATCTCCTCTGTTGCAGCTCTCGCTCATATATAAAAGCGTAGTTGAAGTGACGTTAGAATTTTTAAG

CACTGCACTGTTTTAAAGCGATGGCAATGTGATATGAACGAAATTTCAACTCTTCCAGTGATGTCGCTCTCACCATTTCG

CCACGGTGAGGGGCGCCCTCCATCCTCGGCTGCGAGATCTAACAGTGATGCGCTCGCGTGCTCTAGTACTCAAAATCGCC

TCTCTGGTTGACAGGAAATTCTTGCAATTTATGATTTGCGGCGTCAGCATTTCTGGTGTTCACACTCTATTTCACGAGTG

GGGACCCTAACCAGAAATTAGGCCTATCGGAAGGTCGTAGTTGTCCTATGATCCTCTTGCCCCCTATGCCATAGCTCTGG

CATTCTGTGGTTTCCTGGCACCATGCATTCCTCTGGCACACAGCGGGCGTCTATTGACAACAGACACACGCGCTTATGCA

GTTCCCGCTTGGGGTTAACTTTACCTTACCATTAGTAGTCCGTTCTTTAACCTACGGATCCTCCAACTGACACTTTATGG

TGCCACTCGAGTAGTCTGCTCCTCAACCAACTGATCCTGCGACTAAGGCTTGAACATTTCAACTGTAAGCGCTGCAACAT

TAGTATTAGGCTTGTGCTACAGCTGCAGCAGCTGTAACTGAGCTGATGGTGTGCAGCAGCTGTCACAACAAAGATCTGGA

AGCTCAAGAGCGTAGCACTGTTGTGTAGGTCTACGAACTTCTCCTGCGCAATTGAATGAGTTGCACCAATGCATTACTAT

GGTAATATTTATTTTAGTCTTTGTGTTCCACTGTTTCGGCTGTAGGGCGCAACTTCCTTATGATCTAATTATTATTGTAT

TGTAGGCGTATGTGTGTGTGTGTGAACTTTATCTGTACTTATGTAAGCTTGCTTGTGTTGGGCCCAGACTCTGTACTACT

TGACAGTCCCCTTCAGATGTTGATGCCCAGACTGTGCCCTGCCTGACAGTCCCCTTCAGATGTCGGGGTCCAGACTGTGC

CCTGCTTAACAGTCCCCTCCAGATGTTAGGGTAAAGACTGTGCACTACTTGAGAGTTTACTGTAGTTAGGACTTGTTGCT

ATTGAGTTCTTTCTTTCTTTCATTTAGGTACACTGCGGTTACATCGTTTACGATGCCATACAAACAGCGAGCAAACGGGT

GCTGGTGCTGCATGCAGTGTACTGGTCCTGGTGCTGCATGCAATGTACTGGTCCTGTTGCGGCATGCAGCTTACTAATGA

GCAGCAACTGCCTTACGTTACAACAGGCAGGCCCTATTACGACACCGAATTGATATAAACTTGATCAAAATCTTCGTACA

TAATTTTTTTTTTCTTATGAACTATCATATAAATATAGACTTATCGAATAGGTATTCCAGCTCAACAACGTGGTACATAT

AACCTAAATCAAAGAACTAATTATCTTATAAACTTGATCGTTCAATATATACTTTTATTAAAATTTAATTTTTACATCTT

ATTATTATTATTTCTTAAATACAAAAGTCTAATTAACTTCTGAATGTCTCAAGAAAACTTTCTAACTTGCAAACTTGCAA

GCCGTTCCTTAACCAAGCCTCCAGGAAGCTGCATTACAGGGCCCAGCCTCGGATTATGCTAACTGCGACCCACCTGTCCC

ACTGTCCTCCTCTCACCCGTTACTGACTAACTTAGCACGGCAACTGATCGCGTGCCGACTCTGCGCATCTCCTTGCATCG

CGTCGCTTGCAGTGCAGTGACTGGGAGCTCGCGCCCTTAGTTATCCAGACCAATTACGAATATTTCTATTGAAATGAGGT

GCTTTTTGATGTTCAGATATTCTAGGAAAAATCTAGTTAAATTTTTTTTCTGAAATCATTAAAAGCCGTAGTTATACTGT

TCTCGTAGTTTTTTATGTTTTGCCAGTGAATGATATTTTCTACCGCATTTTTTCATTTGGATGATATGTTTATTCGAAAT

TGCCAACACTTTATACCGAGCGATAATAATTTTCCCTATTTGTTTGTGGTGAAAGCCATGCAGTTCTCAGCGTATTTAAA

ATGCAAGTGCAAAGAATGATCAAATTCATTGAGTTTACTGTCGATCCACGTGACGTCAGAGCTAGGGAAGATGGGGGCGC

TTACGATGCTGCCTGCGCTACTGTCGTTCCACGTGACGTCAGAGCTAGGGAAGATGGGGGGCGCTCACGATGCTGCCTGC

GCTACACTGAGGGAATTTTTGTCCCAAAACCGAAACACTCCGCTGCTGCTGCTGCTAGAAAGTGGTAGCAAACTCCCCTA

CTGCTGCAAGATAGTTGTAGCAAACACAGAGTGCGGAAGGAGAGAGGT

Methoprene tolerant

>PH.k25.comp147496_seq0

CATAGGCTCAACATCACTGCACTCAGAGGATGATTCCTCGGGCACAGAAAGTGAGGACCGCAAGCCCTTCCATGGTCAGG

ACACTCACCTCCTCGTTGCGTTCGTGCAGCTCATCAAGGAGCGCCCCATTACGGAGCTGTCCAGACTGGAAACCCTGAAG

GATGAATACATTACCCGACACGATGTCTATGGCACCATACTGTACACCGATCACAGGATATCATTCGTTACCGGCTACAT

GCCTGAGAGTGTCAGCGGCCTGTCTGCCTTCCTATACGTGCACC

Pigment dispersing hormone

>PH.k41.S6600835

CCGACCTCTGTCACTCGACGCCTGGAGTTCTTGGTGTTCATAGCCTCTTCGTCTCGCCGCCAACTACGCTGCTTGCCTTG

TACCCATGATGATGTTGCAGCTGGCCGCAGTTCCTGCTGATAAGATATCACATCGCAGTATCCTCGCCCTGATGCTCCTG

GGAGTGCTGGCATGCATGCACTTCGTTTATGCTCAACCCAGACGACATCACGAGCTCTACACTGACGACGATATGAACAC

GATGGAGGGCGAAGGCGTGCCGTACAGCAGCAGTGTAGCTGATGATCTCGCCACTTGGATCGTACGCAACAGGATGCCGA

AGCGCAACGCTGAACTGCTCAACACTCTTCTCGGCTCCAGGAACTTGGTGGCGCTGCGGGCAGCAGGCCGTCGATAAGCA

AACATTTCTCCAAATATATTTCGCAGAGTGCAAGGAGCTCCCACTGCAAGATACCTTTATTAGTGGCAAAATAATCCAAA

GTATTACAAAACTTTTAAAAAAATAATTATAATTACTACTAGGACGTCAAATAATTACCGGTAATGGAAATAATCATTGC

TGGAGCGAAACTCAAACTTTGTAACAAGCTTCCCAGATGCAATTACTGTTAGAGCGTTCTGCTCTGGAAATAGCGTTGAG

ATTTGCTAGTCTCACAGCCTCGTGCACTTCCAGCACAGCACTGAGCATAGCAAGAGATACTACCCAGCAGCACAATATTG

CGGTGTATCACACACAGCATGTCGTCGTCCTGTTGACACGTTAAAATACAACTTTTTCAATACACGTGGATAATA

Predicted structures

Red – signal peptide

Yellow – signal peptide cleavage site

Blue – dibasic cleavage site

Underlined – mature peptide.

RPLSLDAWSSWCS-PLRLAANYAACLVPMMMLQLAAVPADKISHRSILALMLLGVLACMHFVYAQPRRHHELYTDDDMNTMEGEGVPYSSSVADDLATWIVRNRMPKRNAELLNTLLGSRNLVALRAAGRR-ANISPNIFRRVQGAPTARYLY-WQNNPKYYKTFKKIIIITTRTSNNYR

-WK-SLLERNSNFVTSFPDAITVRAFCSGNSVEIC-SHSLVHFQHSTEHSKRYYPAAQYC

GVSHTACRRPVDTLKYNFFNTRG-
